# Explainable t-SNE for Single-Cell RNA-seq Data Analysis: A Cell-Driven Information Fusion Framework

**DOI:** 10.64898/2026.09.29.755455

**Authors:** Henry Han, Tianyu Zhang, Luyu Niu, Jiacun Wang, Fei Han, Chun Li, Junyi Li, Huaixin Cao

## Abstract

Single-cell RNA sequencing fundamentally depends on defining cell–cell similarity to resolve distinct cell states. However, conventional dimensional-reduction frameworks such as t-SNE and UMAP inherently rely on Euclidean geometry, a construct ill-suited for high-dimensional, sparse count matrices, and produce cluster boundaries that lack biological interpretability. Here, we introduce cell-driven information fusion, an end-to-end framework that constructs cell affinity directly from raw counts by integrating three biological signals: gene detection patterns, expression magnitude of dominant transcripts, and global profile correlation. A sparsity-adaptive operator merges these signals into a unified affinity matrix, while an entry-usage diagnostic traces which biological component governs local cluster boundaries. The framework comprises two complementary implementations: c-TSNE for low-to-moderate sparsity (*<* 80%), which consistently outperforms Euclidean baselines across six benchmarks, and c-UMAP for high-sparsity regimes. Evaluated across 29 datasets and two protocols against 11 baseline methods, c-UMAP achieved the highest mean normalized and adjusted mutual information (NMI and AMI) in every setting, with all improvements remaining significant after Bonferroni correction. Because a single fixed weight vector was applied across all 29 datasets per protocol, these gains reflect intrinsic algorithmic robustness rather than dataset-specific hyperparameter tuning. Furthermore, c-UMAP scales seamlessly to 1.14 million cells, running up to 22-fold faster than the leading baseline. Crucially, by operating directly on raw counts without variance-based gene selection, our approach avoids the median 46.9% loss of top differentially expressed genes per cell class, preserving essential lineage markers such as *IAPP* and *FOXP3*. Together, these results demonstrate that atlas-scale efficiency, raw-data preservation, and mechanistic interpretability are fully compatible.

## 1. Introduction

Single-cell RNA sequencing (scRNA-seq) reports the transcriptome of one cell at a time. Bulk sequencing averages over a tissue and hides the variation inside it, whereas scRNA-seq resolves that variation directly. The technology has revealed cell subtypes that bulk data could not separate, and it has clarified how immune responses, neural circuits, and tumors are organized. A typical experiment yields a count matrix whose rows are cells and whose columns are genes. Turning that matrix into biological knowledge requires grouping the cells into populations, so clustering is the step on which most downstream interpretation rests.

Clustering rests in turn on a single upstream choice: how the similarity between two cells is measured. Analysis pipelines almost always embed the count matrix into two dimensions with t-SNE [6] or UMAP [8], then cluster the embedding. Both methods build their neighborhoods from Euclidean distance. That choice is the problem this work addresses. Euclidean distance weights every gene equally and assumes the coordinates are dense and informative. scRNA-seq count matrices violate both assumptions. They carry tens of thousands of genes for a few thousand cells, and most entries are zero, partly because a gene is genuinely silent and partly because sequencing failed to capture it. As dimension grows, Euclidean distances between points concentrate around a common value, so near and far neighbors become hard to tell apart [5]; Lemma 1 states this effect precisely. A similarity measure that flattens in exactly the regime where the data live will place cells by geometric accident rather than by biology.

The consequences matter well beyond embedding quality. When the distance is not tied to any biological quantity, a cluster boundary cannot be traced back to the transcriptional evidence that produced it. A biologist who sees two adjacent groups in a t-SNE plot has no principled way to ask which genes made them adjacent. Cluster identities are then assigned after the fact by inspecting marker genes, which turns the embedding into a black box that happens to sit in the middle of an otherwise evidence-based workflow. Rare populations and disease-associated states are the cases most exposed to this gap, because they are the ones a distorted geometry is most likely to merge into a neighbor.

Three obstacles keep current methods from closing the gap. The first is that improvements have targeted accuracy rather than explanation. Imputation methods reduce apparent zero inflation, yet systematic evaluation finds their effect on downstream clustering inconsistent [18]. Deep generative models handle zero inflation and batch effects well, but their latent dimensions correspond to no named biological quantity. SIMLR [4] learns a similarity through multi-kernel integration and performs strongly, though what the learned kernel measures biologically remains open. Recent models including scDCCA [34], scGAC [35], and scICE [33] raise accuracy further while adding depth that makes the mapping from expression to cluster harder to trace.

The second obstacle is that standard pipelines discard evidence before any method runs. Selecting a few thousand highly variable genes is a variance-based decision made without reference to labels, and Section 4.12 shows it removes a large share of the genes that define the very populations being sought.

The third obstacle is scale. Published atlases now hold 10^5^ to 10^6^ cells, and several otherwise strong methods are bounded by an *N × N* matrix they must construct.

This work extends our earlier explainable t-SNE for single-cell data [49], which introduced the presence/absence reading of raw counts, to a full fusion framework with a second backbone and atlas-scale evidence. We address these obstacles with *cell-driven information fusion*, a framework that builds cell similarity from biological signals intrinsic to single-cell expression instead of importing a metric from another domain. A cell is represented by its raw count vector, and three named views are computed from it. The presence/absence view asks which genes are detected and is measured by Yule’s Y. The dominant-signal view asks how strongly the top-expressed genes are expressed and is measured by the L-Chebyshev distance introduced here. The holistic view compares whole expression profiles and is measured by the fractional distance. A sparsity-adaptive operator fuses the three into one affinity, and the resulting embedding carries a diagnostic that attributes each fused entry back to the view that produced it. Two instantiations follow the same principle: *c-TSNE* for data below 80% sparsity, and *c-UMAP* at or above that level. Here sparsity means the proportion of zero entries in the count matrix. We make the following contributions.

1. *Biologically explainable metric fusion*. c-TSNE/c-UMAP fuses three complementary biological factors: (i) gene presence/absence, modeled by Yule’s Y; (ii) the dominant transcriptional signal of top-expressed genes, modeled by the proposed L-Chebyshev distance; and (iii) the holistic expression profile, modeled by fractional distance. Each metric encodes a distinct, non-redundant biological perspective, ensuring that no single similarity bias dominates the embedding.
2. *Preprocessing-free design*. c-TSNE/c-UMAP operates directly on raw counts, reducing dependence on normalization and other preprocessing steps that may suppress marker-gene contributions and obscure true biological heterogeneity. By directly capturing presence–absence signals, highlighting top-expressed genes, and preserving the data’s intrinsic biological heterogeneity, c-TSNE/c-UMAP generates more robust and interpretable embeddings than methods reliant on preprocessed inputs. This eliminates preprocessing dependencies and enhances reproducibility. It also removes the need to correct for library size. Sequencing depth scales every count in a cell by one factor. Binarization does not see that factor. Under the Raw protocol the fused distance puts 0.90 of its weight on the presence/absence view (Section 4.9), so nine tenths of it is untouched by depth. By retaining raw-data stochasticity, c-TSNE/c-UMAP captures transcriptional bursting and subtle biological distinctions, offering a streamlined, biologically faithful, and explainable alternative to conventional workflows.
3. *Sparsity-adaptive fusion*. The framework automatically selects max-fusion or weighted sum-fusion according to data sparsity, ensuring that the dominant biological signal governs the embedding across different sparsity regimes.
4. *Performance at scale, with one fixed setting*. c-TSNE leads every Euclidean-geometry baseline on six well-characterized benchmarks. We also built a 29-dataset benchmark spanning 225 to 1,136,218 cells. Its sparsity ranges from 45.0% to 98.0%. Evaluated there under two protocols against 11 baselines and two indices, c-UMAP attains the highest mean NMI and AMI in every setting. It leads all eleven baselines on both indices, and all 22 Wilcoxon signed-rank tests survive Bonferroni correction. This result uses *one fixed weight vector per protocol*, applied unchanged to all 29 datasets, so it reflects the fusion principle rather than per-dataset tuning.
5. *Atlas-scale scalability*. The lean implementation contains no *N × N* step and reaches 1.14 *×* 10^6^ cells with a measured *t* ∝ *n*^1.01^, running up to 22*×* faster end to end than the strongest baseline method.
6. *Measured cost of preprocessing*. We quantify what standard pipelines remove: variance-based selection of 2,000 genes discards a median 46.9% of each class’s top-50 differentially expressed genes. That amounts to 15,121 (dataset, class, gene) instances over 4,988 distinct genes, including canonical markers such as *IAPP, GCG, CTRB1, FOXP3* and *MS4A1*. This supplies the empirical case for a preprocessing-free design.
7. *Theoretical guarantees*. We prove that high sparsity induces distance flattening and weakens local neighborhood contrast, thereby explaining the crowding and reduced segregation observed in sparse scRNA-seq embeddings. We further prove that max-fusion perturbs the fused distance less than sum-fusion does at high sparsity (Theorem 4), and we characterize the high-dimensional behavior of Euclidean distance (Lemma 1).

Broadly, cell-driven information fusion provides a biologically explainable framework that redefines cell similarity in scRNA-seq analysis using interpretable biological signals extracted directly from raw scRNA-seq data, rather than relying on generic, data-agnostic machine-learning pipelines. By improving the biological interpretability of cell-similarity estimation, c-TSNE and c-UMAP enable more reliable identification of cellular heterogeneity, rare cell populations, and disease-relevant transcriptional patterns, thereby enhancing the biological reliability of machine-learning-driven single-cell analysis. Beyond single-cell clustering, this work offers a general principle for designing data-specific and explainable information-fusion methods for high-dimensional biomedical data. Explainability is not an add-on to machine learning. It is a prerequisite for learning that stays biologically faithful.

## 2. Related Work

Single-cell clustering has evolved rapidly over the past decade, advancing from classical statistical pipelines to graph-based methods, similarity learning, imputation, and deep learning frameworks. We organize existing methods into five categories and relate each to the central goal of this work, which is making cell similarity biologically explainable.

### Classical dimensionality reduction

Early single-cell pipelines applied PCA followed by k-means or hierarchical clustering. That route is transparent, but it captures nonlinear biological structure poorly. t-SNE [6] became a dominant visualization method for scRNA-seq because of its ability to preserve local neighborhood structure, with parameter settings later optimized for large data [27], and UMAP [8] subsequently emerged as a faster and more scalable alternative that was quickly adopted for single-cell work [3]. PHATE [32] introduced diffusion-based geometry to capture developmental trajectories, while FIt-SNE [2] accelerated t-SNE through grid interpolation for large datasets. Despite their broad use, these methods are commonly applied in scRNA-seq pipelines with Euclidean, PCA-derived, or other generic distances to construct pairwise affinities or neighborhood graphs. Such distances are often poorly aligned with the high-dimensional and sparse structure of scRNA-seq data [5]. As a result, the geometry of the embedding is not explicitly tied to biological factors such as gene detection patterns, dominant expression signals, or global transcriptional profiles, making cluster boundaries difficult to validate biologically.

### Graph-based and similarity-learning methods

Seurat [40] and Scanpy [41] build shared nearest-neighbor graphs from PCA-reduced representations and apply community detection, commonly using Louvain or Leiden algorithms [39], to identify clusters. These pipelines are powerful and flexible, but their outcomes depend on multiple interacting choices, including normalization, highly variable gene selection, PCA dimensionality, neighborhood size, and clustering resolution. The combined effects of these choices are often difficult to disentangle biologically. Semisoft clustering [16] addresses intermediate settings, and SIMLR [4] learns cell similarity through multi-kernel learning. Although these methods can achieve strong clustering performance, the resulting similarity measures are not explicitly decomposed into named biological factors. Consequently, it remains difficult to determine whether a cluster boundary is driven by gene detection, dominant marker expression, global transcriptional differences, or preprocessing-induced effects.

### Imputation-based approaches

A distinct line of work addresses sparsity by estimating missing expression values before clustering. MAGIC [42] propagates expression signals across a cell-similarity graph, and related methods fit zero-inflated factor models or regression models to recover dropout values. Although these approaches can reduce apparent zero inflation, systematic evaluations have shown that imputation does not consistently improve downstream clustering and may introduce spurious co-expression patterns [18]. More fundamentally, imputation can blur the distinction between true biological zeros and technical dropouts, thereby removing sparsity patterns that may themselves carry biological information. In contrast, our framework operates directly on raw counts and treats sparsity not as a nuisance to be erased, but as an explicit factor governing information fusion.

### Deep learning methods

Deep learning methods form a major class of high-performing approaches for modeling the statistical complexity of scRNA-seq data. Variational and count autoencoders model the zero-inflated negative binomial likelihood of the counts, and scDeepCluster [43] jointly optimizes an autoencoder and a clustering objective. More recent methods further incorporate contrastive learning, graph attention, and consensus refinement: scDCCA [34] uses a deep contrastive autoencoder, scGAC [35] aggregates neighborhood information through graph attention, and scICE [33] combines multi-resolution Leiden clustering with consensus refinement. These methods can achieve strong quantitative performance, but their multi-layer nonlinear architectures often lack an inherent, transparent mapping from raw gene expression to cluster assignments. The learned latent dimensions usually do not correspond to named biological factors, and cluster boundaries are difficult to trace back to interpretable expression mechanisms. Thus, as model complexity increases, the gap between predictive performance and biological explanation often widens [7, 20].

### Scalable methods

The growth of atlas-scale scRNA-seq datasets has motivated scalable methods capable of handling tens of thousands to millions of cells. UMAP [8] achieves efficient optimization on a *k*-nearest-neighbor graph; and scICE [33] improves scalability through multi-resolution consensus clustering. These methods address computational cost and enable large-scale analysis, but scalability alone does not resolve the biological interpretability of cell similarity. In many cases, scalable pipelines still rely on PCA-derived representations, Euclidean neighborhoods, or learned latent features whose connection to scRNA-seq-specific biological signals is indirect.

### Explainability gap and positioning of this work

Across these five categories a common opportunity emerges. Existing methods generally treat cell similarity as a computational quantity to be optimized, which leaves open the possibility of treating it as a biological concept to be explained. They may improve visualization quality, clustering accuracy, denoising, or scalability. Few tie their similarity measures to named biological factors. Fewer still expose how several biological signals are fused, or quantify which axis drives each embedding. Our cell-driven framework addresses this gap directly. c-TSNE and c-UMAP fuse three biologically named dissimilarity views under a sparsity-adaptive operator; the entry-usage diagnostic attributes the fused affinity back to its biological sources; and the full pipeline operates on raw counts without preprocessing. The framework therefore aims at embeddings that are competitive in accuracy and, at the same time, interpretable through the single-cell mechanisms that define biologically faithful cell similarity.

## 3. Cell-driven information fusion: cell-driven explainable t-SNE (c-TSNE) and UMAP (c-UMAP)

The central idea of the cell-driven framework is to define cell similarity through biological factors intrinsic to single-cell gene expression, rather than importing a generic metric from other domains. A cell is represented by its raw count vector, from which three complementary views are constructed. The presence/absence view records which genes are detected. The dominant-signal view records how strongly the leading genes are expressed. The holistic view compares all expressed genes at the global level. These three views are fused into a single cell-driven affinity that governs the low-dimensional embedding. The fusion operator is sparsity-adaptive: at high sparsity, max-fusion lets the dropout-robust Yule view dominate; at lower sparsity, weighted sum-fusion draws on all three views equally. The resulting framework is fully preprocessing-free, operating on raw counts without normalization, imputation, or gene selection. It is also explainable by design: each component of the fused distance maps directly to a named biological factor. Figure 1 gives an end-to-end overview of the pipeline.

**Figure 1:**
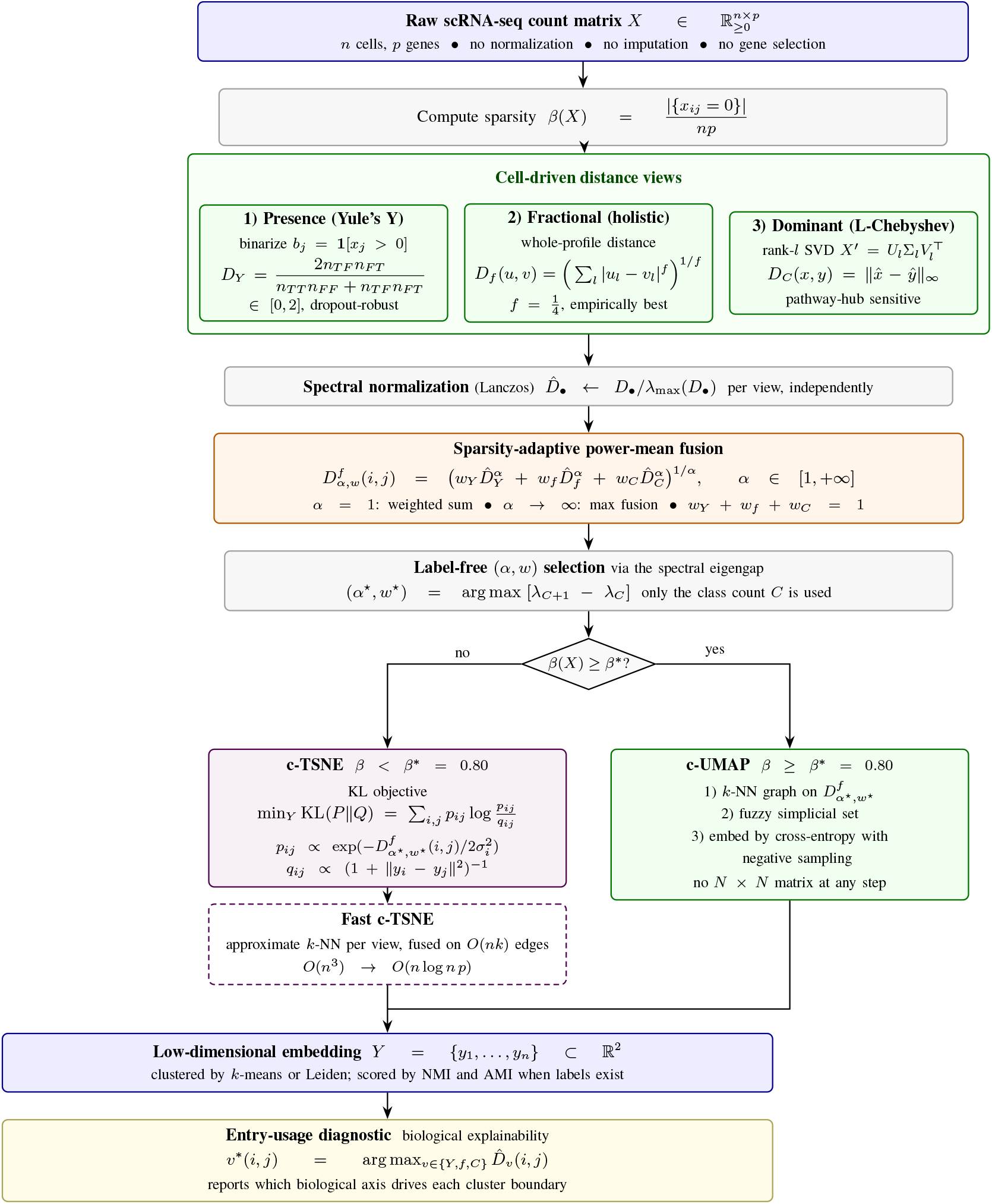
Cell-driven information fusion pipeline (c-TSNE and c-UMAP). Three biologically named dissimilarity views: Yule’s Y (presence/absence), Fractional (holistic expression), and L-Chebyshev (dominant signal) are fused via a sparsity-adaptive power-mean operator with label-free (*α, w*) selection. Sparsity *β*^∗^ = 80% routes computation: **c-TSNE** (KL divergence, gradient descent) for *β <* 80%, with fast c-TSNE reducing *O*(*n*^3^) to *O*(*n* log *n p*); **c-UMAP** (fuzzy simplicial set, SGD) for *β* ≥ 80%. The entry-usage diagnostic attributes each fused distance to its dominant biological source (*Y, f*, or *C*), exposing which axis drives each cluster boundary.

### 3.1 Notation

Table 1 collects every symbol used in this work. Each symbol is also defined where it first appears in the text.

**Table 1:**
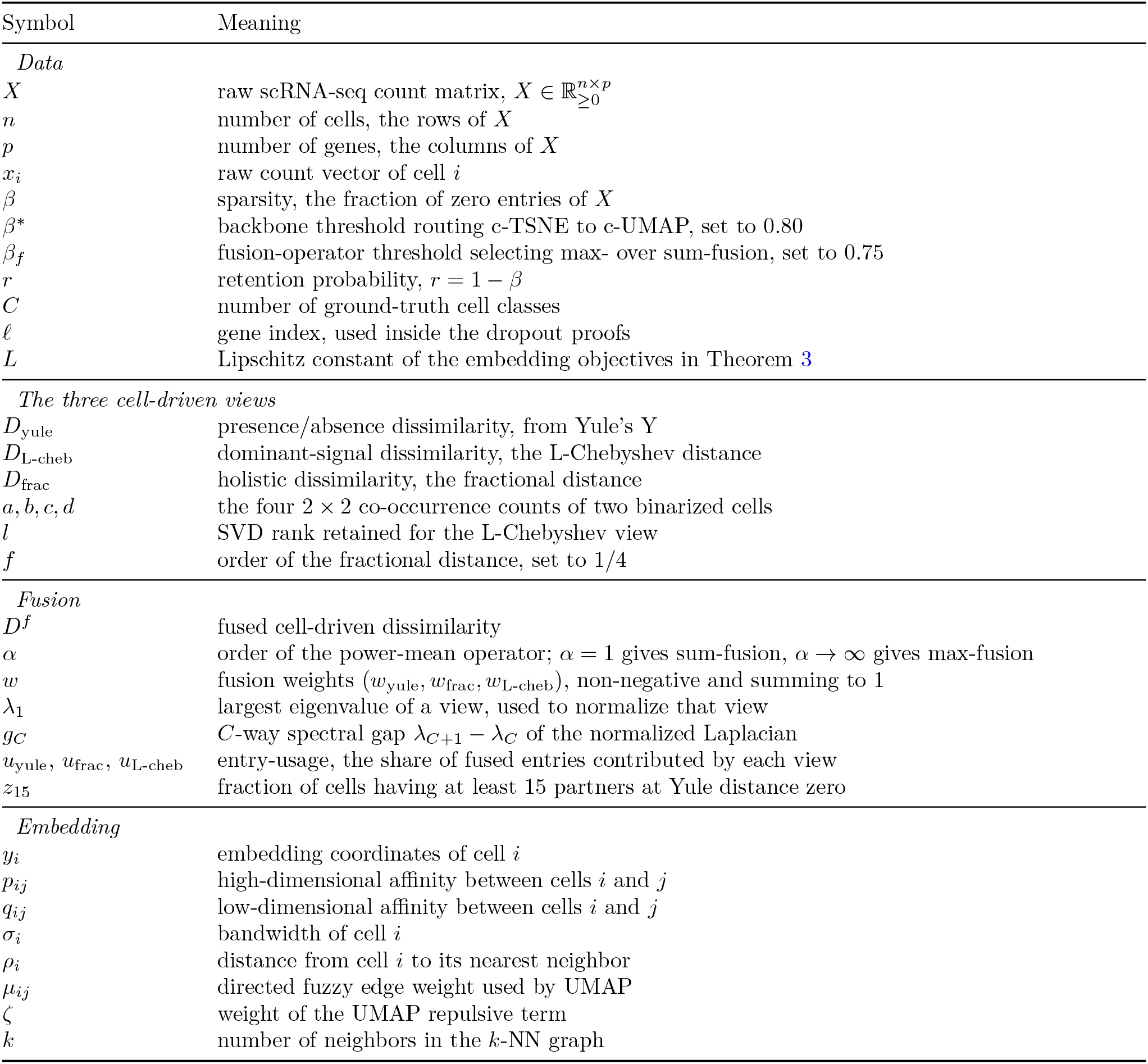
Notation used throughout this work.

**Table 2:** Implementation complexity of c-TSNE. Fast c-TSNE reaches *O*(*n* log *n p*) with fidelity bounded by Theorem 3.

| Stage | Naive | Fast c-TSNE ($k = O(\log n)$) |
| --- | --- | --- |
| $k$-NN graph (per view) | $O(n^2 p)$ | $O(n \log n p)$ |
| Fused distance on edges | $O(n^2 p)$ | $O(n k p)$ |
| L-Chebyshev low-rank SVD | $O(n p^2)/O(n^3)$ | $O(n p r)$ randomized |
| Per-view normalization | $O(n^3)$ full eigen | $O(n k)$ median-of-edges |
| Fusion + $(\alpha, w)$ selection | $O(n^2)$ | $O(n k)$ on edges |
| Affinity / perplexity calib. | $O(n^2)$ | $O(n k)$ (sparse) |
| t-SNE gradient opt. | $O(n^2)/\text{iter}$ | $O(n \log n)/\text{iter}$ (BH) |
| **Total time** | **$O(n^3)$** | **$O(n \log n p)$** |
| **Memory** | **$O(n^2)$** | **$O(n k + \text{nnz}(X))$** |

### 3.2 scRNA-seq data: high-dimensional and sparse

Single-cell RNA sequencing produces a count matrix 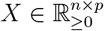, where rows are cells and columns are genes. Two structural features distinguish scRNA-seq from standard multivariate data. First, the data are **high-dimensional**: *p* ≫ *n*, with typical gene panels of 10,000–30,000 features and only hundreds to thousands of cells. Second, the data are **extremely sparse**: most entries are zero. Two causes mix. A gene may be genuinely silent in that cell type, or it may be expressed but missed by the sequencing reaction. These two properties jointly invalidate standard Euclidean distance as a cell-similarity measure (Lemma 1) and motivate the three cell-driven dissimilarities defined in the subsections below.

#### Data sparsity

For *X* = *{x*_1_, *x*_2_, …, *x*_*n*_*}, x*_*i*_ ∈ ℝ^*p*^, sparsity is the fraction of zero entries,

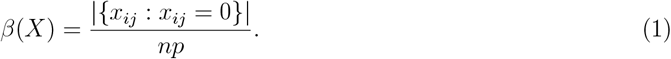

Sparsity is rarely reported yet is intrinsic to scRNA-seq, being closely tied to dropout: larger *β* implies more zeros and more potential dropout. Because *β* routes the backbone and selects the fusion operator (and the Yule view directly targets sparsity), it deserves explicit attention; without imputation it spans 45.0–98.0% across the 29 datasets benchmarked here.

#### Main implication

Theorem 1 shows that independent dropout introduces a dominant, pair-independent masking background into observed squared distances. Conditional on the latent expression matrix,

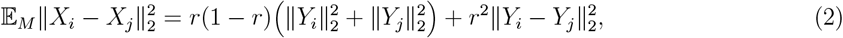

where *r* = 1 − *β* is the retention probability. Hence the biologically meaningful distance contrast is suppressed by an additional factor of *r*. Under standard concentration conditions, specifically *rp* ≫ log *n*, and assuming that the resulting between/within distance gap satisfies 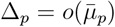, all observed pairwise distances concentrate around the same scale. Consequently, the data-dependent component of the t-SNE KL gradient collapses: if *P* is close to the uniform affinity matrix *U*, then

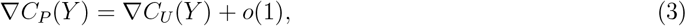

uniformly over embeddings. Thus t-SNE optimization is not necessarily stationary, but it becomes driven by a data-blind uniform-affinity objective rather than by biological cluster structure.

UMAP behaves differently because its optimization is governed primarily by the sparse *k*-nearest-neighbor support graph. Non-edges retain repulsive weight, while the local-connectivity constraint enforces at least one high-confidence edge per point. Therefore, if the *k*-NN graph remains biologically assortative under sparsity, UMAP retains a nonvanishing cluster-separation signal even when continuous edge weights flatten. This establishes a formal distinction: t-SNE fails when dense affinity magnitudes collapse, whereas UMAP remains robust to magnitude collapse provided binary graph topology survives. The proposed 80% switch should therefore be interpreted as a calibrated finite-sample threshold at which dense-affinity signal becomes negligible while sparse graph topology remains informative.

The following theorem says that dropout makes every pair of cells look about equally far apart, so the biological part of the distance stops being visible.

##### Theorem 1

(Sparsity-induced distance flattening). *When expression vectors are observed through independent dropout masks, the observed squared distance between two cells is dominated by a masking background that is nearly the same for all pairs. The true biological distance term is multiplied by r*^2^, *while the dropout background is of order r. As sparsity increases, between-type and within-type distances become harder to distinguish, and all observed distances concentrate near one common scale*.

*Let r*_*p*_ = 1 − *β*_*p*_ *be the retention probability. For i* = 1, …, *n*_*p*_, *let*

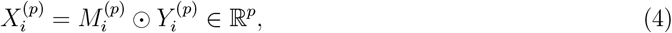

*where* 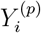 *is the latent expression vector and, conditional on* 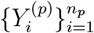, *the entries* 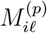 *are independent Bernoulli*(*r*_*p*_) *random variables. Define*

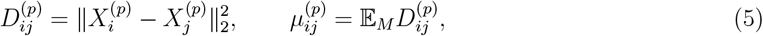

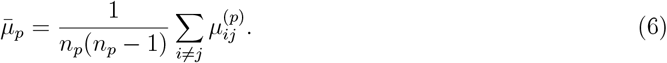

*Then, conditional on the latent vectors*,

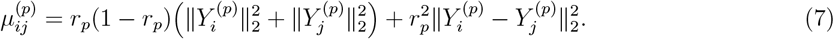

*Assume:* 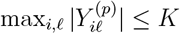 *for some constant* 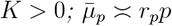 *and r*_*p*_*p/* log *n*_*p*_ → ∞; *and*

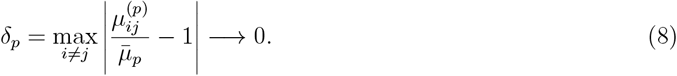

*Then*

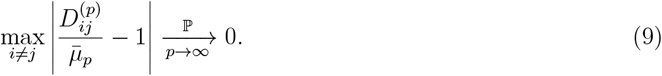

*In particular, if the label-level between/within distance contrast*

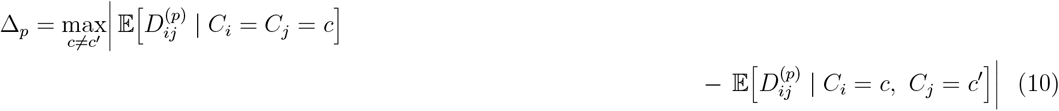

*satisfies* 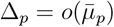, *then biological distance contrast is asymptotically negligible relative to the dominant dropout-induced distance scale*.

*Proof*. For each coordinate 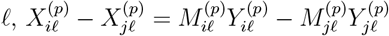, so

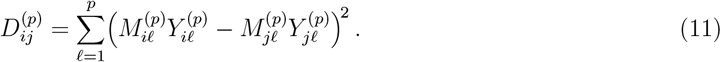

Conditional on latent vectors, summands are independent across *ℓ*. Since

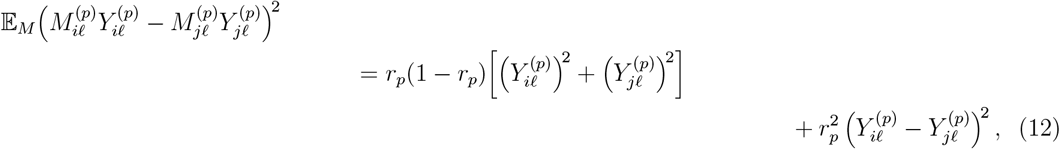

summing over *ℓ* gives the stated formula for 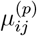. Because 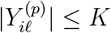, each summand is bounded by 4*K*^2^ and has conditional variance *O*(*r*_*p*_), so 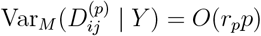. By Bernstein’s inequality,

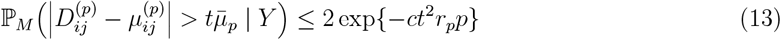

for a constant *c >* 0 depending only on *K*. A union bound over all *n*^2^ ordered pairs gives

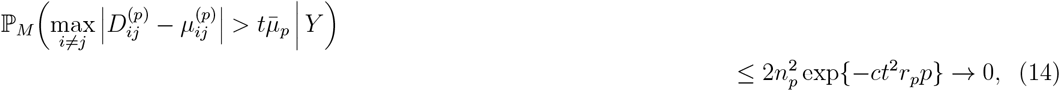

since *r*_*p*_*p/* log *n*_*p*_ → ∞. Combining with *δ*_*p*_ → 0 gives 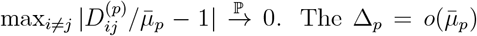 condition then implies that biological contrast vanishes after normalization by 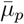.

The benchmark shows the predicted shift. As sparsity rises the fused distance leans on the view that survives flattening: the presence/absence view supplies 41.7% of fused entries on Usoskin at 78.1% sparsity, against 25.6% on Kolod at 27.9% (Table 7).

Theorem 1 motivates replacing Euclidean distance with the cell-driven dissimilarities defined below, which are constructed so that their within-class and between-class gap holds up as sparsity rises. The measurements agree with the theorem. In Table 5 the Euclidean view trails the best cell-driven view on all four benchmarks, and the gap is widest on Buettner, at 0.303 *±* 0.010 against 0.824 *±* 0.013. The effect also tracks sparsity as predicted (Table 7).

> The following theorem says that once distances flatten, the t-SNE objective stops depending on the data, while UMAP keeps a separating force because its loss reads the sparse neighbor graph rather than the distance magnitudes.

##### Theorem 2

(Sparsity-induced KL-divergence bias and UMAP repulsive robustness). *Once dropout makes the high-dimensional t-SNE affinity matrix P close to the uniform affinity matrix U, the data-dependent part of the t-SNE gradient becomes negligible: t-SNE may still move points, but according to a data-blind uniform-affinity force field. UMAP retains a cluster-separation force because its loss depends on the binary support structure of the sparse k-NN graph. Therefore, UMAP remains robust to weight flattening only when k-NN graph topology itself remains biologically informative*.

*Let N*_*p*_ = *n*_*p*_(*n*_*p*_ − 1) *and* 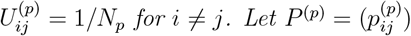 *be the symmetrized t-SNE affinity matrix, and suppose*

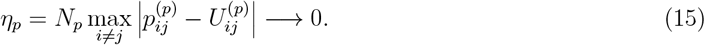

*For an embedding Y* = (*y*_1_, …, *y*_*n*_), *define*

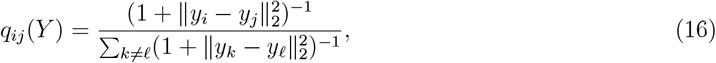

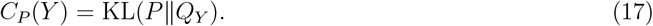

*Then* 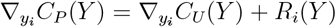 *where C*_*U*_ (*Y*) = KL(*U*∥*Q*_*Y*_) *and*

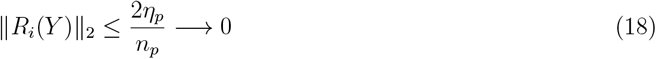

*uniformly over all embeddings Y*. *Thus the data-dependent component of the t-SNE KL gradient collapses as P* → *U*.

*For UMAP, let G*_*p*_ = (*V*_*p*_, *E*_*p*_, *w*_*p*_) *be the fuzzy k-NN graph with* 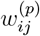 ∈ [0, 1], *and let*

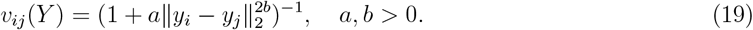

*Consider the cross-entropy*

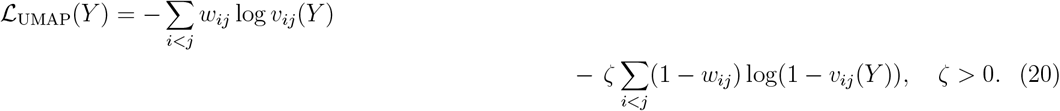

*Assume the cells admit a partition V*_*p*_ = *C*_1_ ∪ … ∪ *C*_*K*_ *satisfying:*

1. ***Within-cluster support survives***. *Each G*_*p*_[*C*_*a*_] *contains a connected spanning subgraph T*_*a*_ *with* 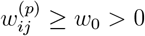 *on T*_*a*_.
2. ***Cross-cluster attraction is absent***. *For* 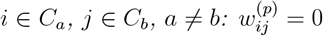.

*Define ℓ*_−_(*r*) = − log(1 − (1 + *ar*^2*b*^)^−1^), *strictly decreasing in r. For two embeddings Y* ^near^, *Y* ^far^ *with identical within-cluster geometry but* 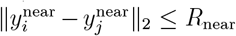 *and* 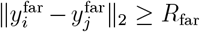 *for all cross-cluster pairs, with R*_far_ *> R*_near_ *>* 0,

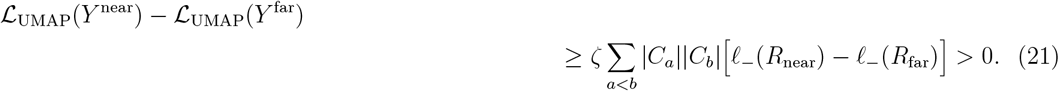

*Hence, if the sparse k-NN graph remains biologically assortative, UMAP retains a nonvanishing cluster-separation signal even when continuous edge weights flatten*.

*Proof*. **t-SNE claim**. Write *p*_*ij*_ = *U*_*ij*_ + *δ*_*ij*_. The standard symmetric t-SNE gradient is

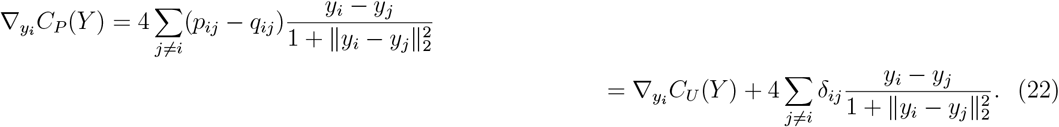

Using 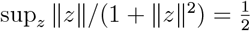 and |*δ*_*ij*_| ≤ *η*_*p*_*/N*_*p*_,

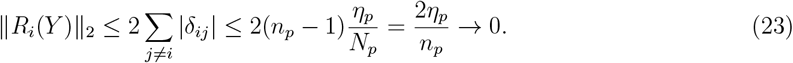

**UMAP claim.** Cross-cluster pairs have *w*_*ij*_ = 0, contributing purely repulsive loss ζ *ℓ*_−_(∥*y*_*i*_ − *y*_*j*_∥_2_). Since *ℓ*_−_ is strictly decreasing and within-cluster geometry is fixed,

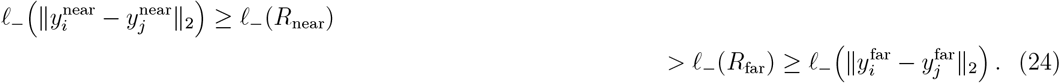

Summing ζ[*ℓ* (*R*_near_) *ℓ* (*R*_far_)] *>* 0 over all ∑_*a<b*_ *C*_*a*_ *C*_*b*_ cross-cluster pairs gives the bound. Positivity follows from *R*_far_ *> R*_near_ and strict monotonicity of *ℓ*_−_.

The benchmark supports the distinction the theorem draws. Across the 29 datasets, whose sparsity reaches 98%, the UMAP-backbone instantiation attains the highest mean NMI under both protocols, at 0.618 and 0.696 (Table 11). This is the regime in which the theorem predicts that graph topology, rather than affinity magnitude, carries the cluster signal.

> The following corollary says that the choice between the two backbones should follow a measurable property of the data rather than a fixed constant.

##### Corollary 1

(Sparsity-adaptive routing principle). *The theory does not make* 80% *a universal mathematical constant. Instead, it justifies a sparsity-dependent switching principle: use c-TSNE while dense affinity magnitudes still contain biological signal, and switch to c-UMAP once dense affinities collapse but sparse graph topology remains informative. The value* 80% *is the empirically calibrated crossing point for the datasets in this work*.

*Let*

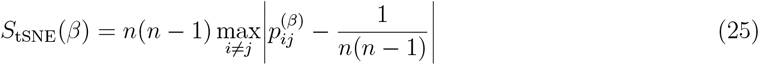

*measure dense-affinity deviation from uniformity, and*

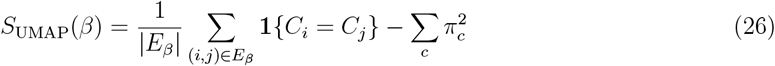

*measure biological assortativity of the k-NN graph (π*_*c*_: *class fraction). The routing rule is: choose* ***c-TSNE*** *if S*_tSNE_(*β*) *> τ*_*t*_; *choose* ***c-UMAP*** *if S*_tSNE_(*β*) ≤ *τ*_*t*_ *and S*_UMAP_(*β*) ≥ *τ*_*g*_. *The empirical threshold β*^∗^ = 80% *is the finite-sample point at which dense t-SNE affinity signal becomes negligible while UMAP graph topology remains sufficiently assortative*.

The two halves of our evaluation bear the routing rule out. The six c-TSNE benchmarks span 27.9% to 78.1% sparsity and sit below the threshold, where c-TSNE leads every Euclidean-geometry baseline (Tables 6 and 9). The 29-dataset collection reaches 98.0% sparsity, and there the c-UMAP instantiation leads every baseline on both indices (Table 11).

### 3.3. Yule’s Y: the presence/absence view

Yule’s Y (the coefficient of colligation, Yule 1912 [24]) measures association between two binary variables. We binarize each cell *x* = (*x*_1_, …, *x*_*p*_)^⊤^ by 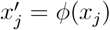 with

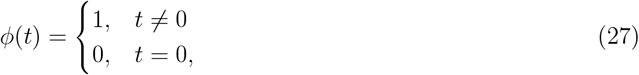

mapping every detected gene to 1 (“expressed”) and every zero to 0 (“not expressed”). Binarization is what frees this view from library-size bias. Sequencing depth multiplies a cell’s counts by a positive factor. That factor does not change which entries are non-zero. So *ϕ* is unchanged, and *D*_yule_ with it. Correcting for depth is what a normalization step is for, which is why the raw-count pipeline does not need one. Given two binarized cells *U* = (*u*_1_, …, *u*_*p*_)^⊤^, *V* = (*v*_1_, …, *v*_*p*_)^⊤^ with *u*_*i*_, *v*_*i*_ ∈ *{*0, 1*}*, define the 2 *×* 2 counts

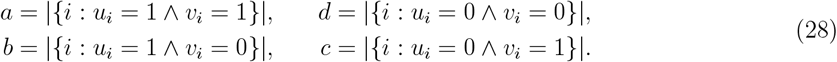

Yule’s Y is

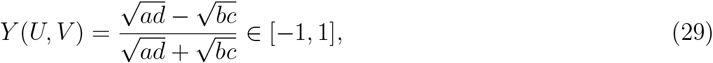

where *±*1 denote total positive/negative association and 0 no association.

Yule’s Y compares the global pattern of presences and absences. Genes that stay silent across related cells therefore emerge as true signal, while sporadic isolated zeros behave like dropout. That is what makes this view robust to zero inflation. For scalability, the c-UMAP implementation uses Yule’s *Q* = (*ad* − *bc*)*/*(*ad* + *bc*) in dissimilarity form, 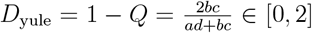 (Eq. (40)); *Q* and *Y* are monotonically related binary-association coefficients computed from the same 2 *×* 2 counts.

### 3.4. L-Chebyshev: the dominant-signal view

Top-expressed genes contribute much of the data variance and often act as pathway hubs. To isolate them we first low-rank-approximate *X* via SVD [25]; efficient updates of a truncated SVD are available when the matrix evolves [19]. For *X* = *{x*_1_, …, *x*_*n*_*}, x*_*i*_ ∈ ℝ^*p*^, *p* ≫ *n*, the rank-*l* approximation is

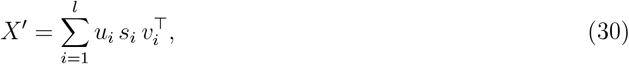

with *l* the smallest rank capturing at least 80% of the variance. We then take the Chebyshev (*L*) distance between approximated cells:

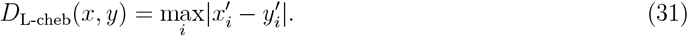

We use Chebyshev rather than Euclidean because, on the still high-dimensional approximations, the Chebyshev distance avoids the amplification that biases Euclidean distances in high dimensions, giving a sharper read on the dominant-gene contribution.

### 3.5. Fractional dissimilarity: the holistic view

Aggarwal *et al*. showed that in high dimensions, standard (*L*_2_) metrics become poorly discriminative [5]. Extending the Minkowski distance to a fractional order gives a more discriminative holistic measure. For two cells *U, V* ∈ ℝ^*p*^,

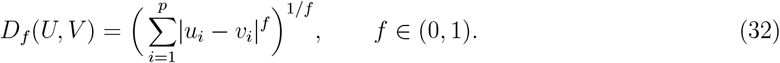

We use *f* = 1*/*4, which empirically gives the best sensitivity. *D*_*f*_ is a *dissimilarity* (a semimetric): it can violate the triangle inequality for degenerate points, but such all-or-nothing cells do not occur in real scRNA-seq, and *D*_*f*_ discriminates cells well in practice.

### 3.6. Cell-driven information fusion

The three views yield three pairwise dissimilarity matrices reflecting three biological factors; because these factors act jointly, we fuse the matrices into one affinity. We first normalize each matrix by its largest eigenvalue [8, 26]: given *D* with eigen-decomposition *D* = *Q*Λ*Q*^−1^,

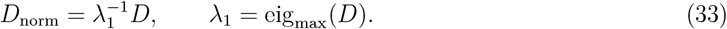

**Max-fusion** takes the entrywise maximum:

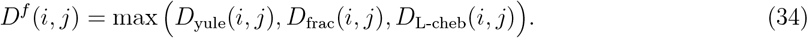

**Sum-fusion** takes a weighted sum:

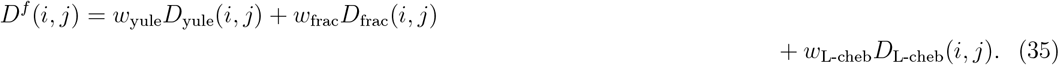

Default weights are 1*/*3 each; for high-sparsity data (*β >* 0.5) we up-weight Yule (e.g., *w*_yule_ = 0.8).

#### A unifying operator: power-mean fusion

Rather than treat max- and sum-fusion as two disjoint heuristics, we embed them in a single one-parameter family, the weighted *power-mean*:

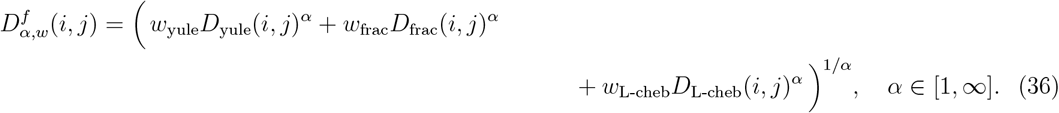

The order *α* interpolates the two operators continuously. Setting *α* = 1 recovers sum-fusion (Eq. (35)), and letting *α* → ∞ recovers max-fusion (Eq. (34)). The choice between max and sum is therefore a position on a continuum rather than a binary decision. Stability under perturbation rises monotonically from *α* = 1 to *α* = ∞, so the design rule “larger *α* for sparser data” is the continuous form of the sparsity-adaptive rule.

### 3.7. What separates c-TSNE from c-UMAP

The fusion is identical in both instantiations: the same three views, the same operator, the same weights. They differ only in what consumes the fused distance, and that difference is forced by sparsity.

c-TSNE keeps the dense affinity matrix and minimizes a Kullback–Leibler objective by gradient descent. It therefore reads the magnitudes of the fused distance, which is what makes it the more interpretable of the two. Those magnitudes flatten once most entries are zero (Theorem 1).

c-UMAP keeps only the edges of an approximate *k*-nearest-neighbor graph and minimizes a fuzzy cross-entropy by stochastic gradient descent. It reads graph topology rather than magnitude, which survives the flattening. Cost follows the same split: c-TSNE builds an *n × n* object, c-UMAP builds none. That is what carries one fusion rule from 225 cells to 1.14 million.

#### Which to use

c-UMAP suits large, highly sparse data: it is the instantiation we recommend at or above 80% sparsity, and it is the only one that reaches atlas scale. c-TSNE suits smaller data below 80% sparsity, where the dense affinities still carry signal and its magnitudes remain readable.

### 3.8. Cell-driven t-SNE (c-TSNE)

c-TSNE follows the t-SNE framework but replaces the Euclidean affinity with the fused, cell-driven affinity *D*^*f*^. Given *X* =*{x*_1_, …, *x*_*n*_*}, x*_*i*_ ∈ ℝ^*p*^, *p* ≫ *n*, c-TSNE computes the embedding *Y* = *{y*_1_, …, *y*_*n*_ *}, y*_*i*_ ∈ ℝ^*l*^ (typically *l* = 2), by minimizing the Kullback–Leibler divergence between a Gaussian affinity *P* and a Student-*t* affinity *Q*:

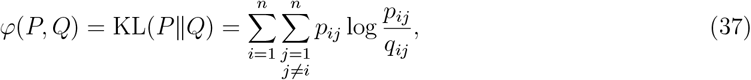

with 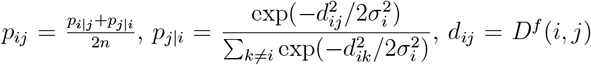, and 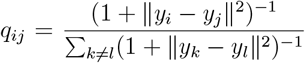. The embedding is optimized by gradient descent,

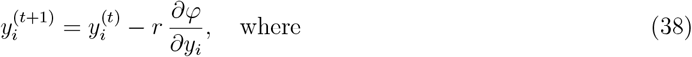

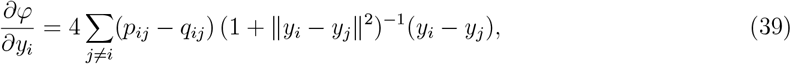

with learning rate *r* set to about 1*/*10 of classic t-SNE’s.

### 3.9. Fast c-TSNE: from O(n^3^) to O(n log np)

The naive *O*(*n*^3^) cost of c-TSNE comes from the full eigen-decomposition and the dense SVD (L-Chebyshev), not from t-SNE itself. **Fast c-TSNE** never forms the full matrix; it builds the fused distance only on an approximate *k*-nearest-neighbor graph.

Fast c-TSNE proceeds in five steps.

1. Obtain candidate neighbors by approximate-NN search under each view.
2. Evaluate and fuse the three cell-driven dissimilarities (Eq. (36)) only on the *O*(*nk*) candidate edges.
3. Replace the *O*(*n*^3^) spectral normalization by the median-of-edges scale.
4. Calibrate the per-cell bandwidth *σ*_*i*_ on the sparse neighborhood.
5. Optimize the KL objective with Barnes–Hut t-SNE, at *O*(*n* log *n*) per iteration, or with FFT-accelerated FIt-SNE [2].

The approximation is not heuristic: its objective deviation from exact, dense-affinity c-TSNE is at most *Lε* by Theorem 3(i).

### 3.10. Scaling the fusion: cell-driven UMAP (c-UMAP)

The atlas experiments adopt the UMAP backbone for its mature, GPU-optimized precomputed-metric implementation. c-UMAP keeps the cell-driven fusion verbatim but feeds it to UMAP, which scales near-linearly and accepts a precomputed metric.

#### Fused cell-driven distance

c-UMAP reads the raw count matrix 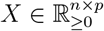 directly, with no normalization, log-transform, scaling, imputation, or gene selection. Let *B* = **1**[*X >*0] be its binarization, and let *n*_*T T*_, *n*_*FF*_, *n*_*T F*_, *n*_*FT*_ be the per-pair co-occurrence counts. The Yule dissimilarity is then

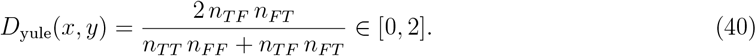

The dominant-signal view uses the rank-*r* truncated SVD 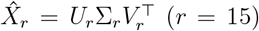 followed by the Chebyshev distance, 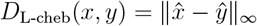, and the holistic view is 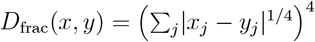.

#### Spectral normalization and convex fusion

Each matrix is divided by its largest-magnitude eigenvalue (Lanczos iteration). The fused distance is the convex combination

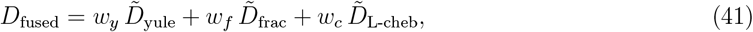

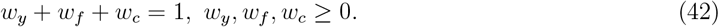

#### UMAP embedding

Given *D*_fused_, UMAP builds a fuzzy simplicial set with directed edge weight

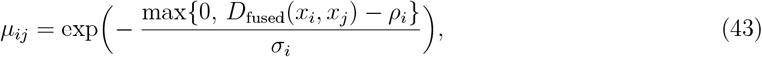

where *ρ*_*i*_ is the nearest-neighbor distance and *σ*_*i*_ solves ∑_*j*_ *µ*_*ij*_ = log_2_ *k* (*k* = 50). The graph is symmetrized and the embedding minimizes the fuzzy cross-entropy by stochastic gradient descent with negative sampling.

### 3.11. When is the UMAP backbone provably better?

#### Assumption 1

(*κ*-neighbor concentration). There is *κ* = *κ*(*n, β, ε*) such that for every cell *i*, φ *d*_*i*,(*κ*+1)_ */*φ *d*_*i*,(1)_ ≤ *ε/n*.

#### Assumption 2

(Planted sparse-support mixture *M* (*β, C*)). Each cell has one of *C* types; type *c* has a gene-support *S*_*c*_ ⊆ [*N*]. A gene in *S*_*c*_ is detected independently with probability 1 − *β*_0_; a gene outside *S*_*c*_ is detected with probability *γ*(*β*), where *γ*(*β*) ↓ 0 as *β* ↑ 1. Distinct supports overlap in at most a fraction *ω <* 1 of their genes.

> The following theorem says that restricting the fused distance to a *k*-nearest-neighbor graph costs almost nothing in objective value, while removing the cubic and quadratic steps that make the dense computation expensive.

#### Theorem 3

(c-UMAP matches c-TSNE at near-linear cost on large, highly sparse data). *Fix a target accuracy ε* ∈ (0, 1) *and let the k-NN graph use any k* ≥ *κ*.

i. ***(Fidelity.)*** *Under Assumption 1, restricting the affinity to the k-NN graph perturbs each cell’s conditional distribution by at most ε in total variation. Consequently the c-UMAP cross-entropy and the c-TSNE objective change by at most Lε, for an absolute constant L*.
ii. ***(Cost.)*** *c-TSNE requires a full eigen-decomposition for normalization (*Θ(*n*^3^)*) and dense Student-t repulsion (*Θ(*n*^2^) *per iteration). c-UMAP avoids both: it needs only the* largest *eigenvalue per view (Lanczos), and its UMAP embedding costs O*(*nk*) *per epoch. c-UMAP is thus strictly cheaper: no* Θ(*n*^3^) *step and no dense softmax*.
iii. ***(Sparsity monotonicity.)*** *Under Assumption 2, κ*(*n, β, ε*) *is non-increasing in β and bounded* independently of *n once sparsity is sufficiently high*.

*Therefore, for any fixed ε there is a threshold n*_0_(*β*) *such that for all n > n*_0_(*β*), *c-UMAP attains the same objective value up to Lε while removing c-TSNE’s* Θ(*n*^3^) *bottleneck; and n*_0_(*β*) decreases *as sparsity β increases*.

*Proof.*

i. For each *i*, the affinity mass beyond rank *κ* is ∑ _*r>κ*_ φ(*d*_*i*,(*r*)_)*/Z*_*i*_. Since *Z*_*i*_ ≥φ(*d*_*i*,(1)_) and φ is decreasing, each tail term is at most φ(*d*_*i*,(*κ*+1)_)*/*φ(*d*_*i*,(1)_) ≤ *ε/n* by Assumption 1; summing the fewer than *n* tail terms gives tail mass ≤*ε*. Both objectives are Lipschitz in their affinities, giving the *Lε* bound.
ii. For c-TSNE, the eigen-normalization is a full eigen-decomposition (Θ(*n*^3^)) and the Student-*t* repulsion sums over all pairs (Θ(*n*^2^) per iteration). c-UMAP replaces the decomposition by a single largest-eigenvalue computation (Lanczos); the UMAP fuzzy graph keeps only the *O*(*nk*) nearest-neighbor edges, and SGD costs *O*(*nkm*) per epoch.
iii. Under *M*(*β, C*), same-type cells have **E**[*O*_*ij*_] ≥ *µ*_in_ and cross-type cells have **E**[*O*_*ij*_] ≤ *µ*_out_. By Hoeffding’s inequality, whenever 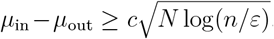, Assumption 1 holds with *κ* ≤ *n*_*c*_ independent of *n*. Because *γ*(*β*) decreases in *β*, the gap increases in *β*, so *κ*(*β*) is non-increasing in *β*.

The benchmark bears the bound out. Fast c-TSNE, which is exactly the *k*-nearest-neighbor restriction the theorem analyzes, reaches a mean NMI of 0.782 against 0.758 for the exact dense computation over the six primary benchmarks, and matches or exceeds it on every one of them (Table A.18). The predicted loss is therefore not visible in the scores.

Theorem 3 formalizes the design rule: use c-TSNE for interpretable analysis at modest *n*, and c-UMAP for large, highly sparse atlases. Measurement confirms both halves of the claim. On fidelity, the *k*-NN restriction proves lossless in practice, with Fast c-TSNE matching or exceeding exact c-TSNE on every dataset tested at a mean NMI of 0.782 against 0.758 (Table A.18). On cost, subsampling one dataset with all else held fixed gives a measured exponent of *n*^1.01^ (Section 4.11), and the lean path reaches 1.14 million cells.

### 3.12. Why max-fusion is more robust than sum-fusion at high sparsity

#### Assumption 3

(Dominant, dropout-robust presence/absence view). Fix a perturbation that changes each normalized magnitude-view entry by at most *η*, i.e. 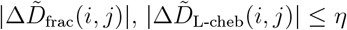. The Yule view is dropout-robust: 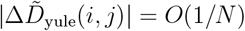. Moreover, at sparsity *β* the Yule view dominates with margin *m*(*β*) on a fraction *π*(*β*) of cell pairs, where both *m*(*β*) and *π*(*β*) are non-decreasing in *β* with *π*(*β*) → 1 as *β* → 1.

> The following theorem says that when the presence/absence view dominates, taking the maximum passes a perturbation through on only a few cell pairs, whereas a weighted sum passes it through on every pair.

#### Theorem 4

(Max-fusion robustness at high sparsity). *Under Assumption 3, for any η < m*(*β*):

a. *On the dominant fraction π*(*β*) *of pairs*, 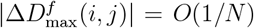; *over all pairs*, 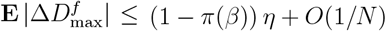.
b. *The weighted sum-fused distance shifts on* every *pair by up to* 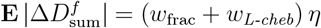.

*Whenever* 1 − *π*(*β*) ≤ *w*_frac_ + *w*_*L-cheb*_ *(i.e*., *for β > β*^⋆^ ≈ 0.8 *on our data), max-fusion has strictly smaller worst-case and expected perturbation of the fused distance than any sum-fusion with w*_*y*_ *<* 1. *The quantity bounded here is the* stability of the fused distance under perturbation, *which is the property the sparsity-adaptive rule is designed around; downstream clustering scores depend additionally on the embedding and the clustering step, and are reported separately in Section 4.10 and Table 11*.

*Proof*. By Assumption 3, on the *π*(*β*)-fraction of dominant pairs the Yule view exceeds the next-largest view by *m*(*β*) *> η*; since the perturbation moves the magnitude views by at most *η < m*(*β*) and the Yule view by *O*(1*/N*), the entrywise argmax stays the (essentially unchanged) Yule view, giving 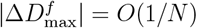 there. On the remaining (1 − *π*(*β*)) fraction the change is at most *η*, yielding (a). For (b), 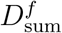 is linear in the views, so the perturbation propagates with its weights, yielding (b). Comparing (a) and (b) gives the claimed superiority for *β > β*^⋆^.

Two measurements support the assumption on which the theorem rests. The presence/absence view does become dominant as sparsity grows, supplying 41.7% of fused entries on the sparsest primary benchmark (Table 7). Independently, the weight selected for the 29-dataset collection places 0.90 on that view under the Raw protocol (Section 4.9), which is the weighting the theorem identifies as stable at high sparsity.

### 3.13. Label-free joint selection of the fusion operator (α, w)

We select (*α, w*) jointly and label-free, by maximizing the *C*-way spectral gap (Davis–Kahan eigengap) of the normalized Laplacian:

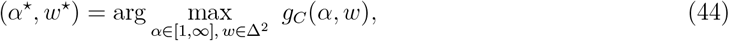

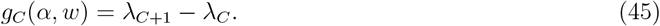

A larger eigengap yields a more stable, better-separated *C*-way partition. The program is low-dimensional; each *g*_*C*_ needs only *λ*_*C*_, *λ*_*C*+1_ (Lanczos), and we solve it by a coarse grid (*α* ∈ *{*1, 2, 4, 8,, *w*∞*}* over an 11-point simplex grid). It is label-free: only *C* is used, which K-means needs anyway, so it cannot over-fit NMI.

#### Validation

Table 3 compares the eigengap-selected (*α*^⋆^, *w*^⋆^) against the two fixed endpoints and the oracle (grid-best NMI). Joint selection attains mean NMI 0.560, **beating equal-weight sum (**0.434**) and max-fusion (**0.406) and doing so with no manual tuning.

**Table 3:** Label-free joint (*α, w*) selection vs. fixed endpoints and oracle (grid-best), scored by spectral clustering (NMI_max_; a selection proxy). Best non-oracle per row in **bold**.

| Dataset | sum ($\alpha=1$, equal $w$) | max ($\alpha=\infty$) | eigengap ($\alpha^*, w^*$) | oracle |
| --- | --- | --- | --- | --- |
| Buettner | 0.279 | **0.381** | 0.279 | 0.770 |
| Kolod | 0.745 | 0.516 | **0.805** | 0.809 |
| Pollen | 0.904 | 0.861 | **0.908** | 0.933 |
| Usoskin | 0.090 | 0.026 | **0.593** | 0.610 |
| Patel | 0.551 | 0.572 | **0.681** | 0.750 |
| Goolam | 0.401 | 0.429 | **0.584** | 0.709 |
| Zeisel | **0.070** | 0.058 | 0.069 | 0.607 |
| Mean | 0.434 | 0.406 | **0.560** | 0.741 |

### 3.14. c-TSNE clustering and evaluation

c-TSNE clustering applies K-means to the c-TSNE embedding. Clustering quality is measured by normalized mutual information (NMI) [28, 29]. For two labelings *U, V*,

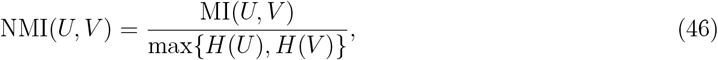

with 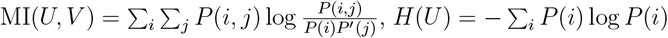, and 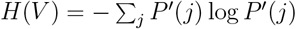.

We use the maximum-entropy normalization for its stability across datasets.

### 3.15. Algorithm

#### Algorithm 1

**Cell-driven t-SNE (c-TSNE) clustering**

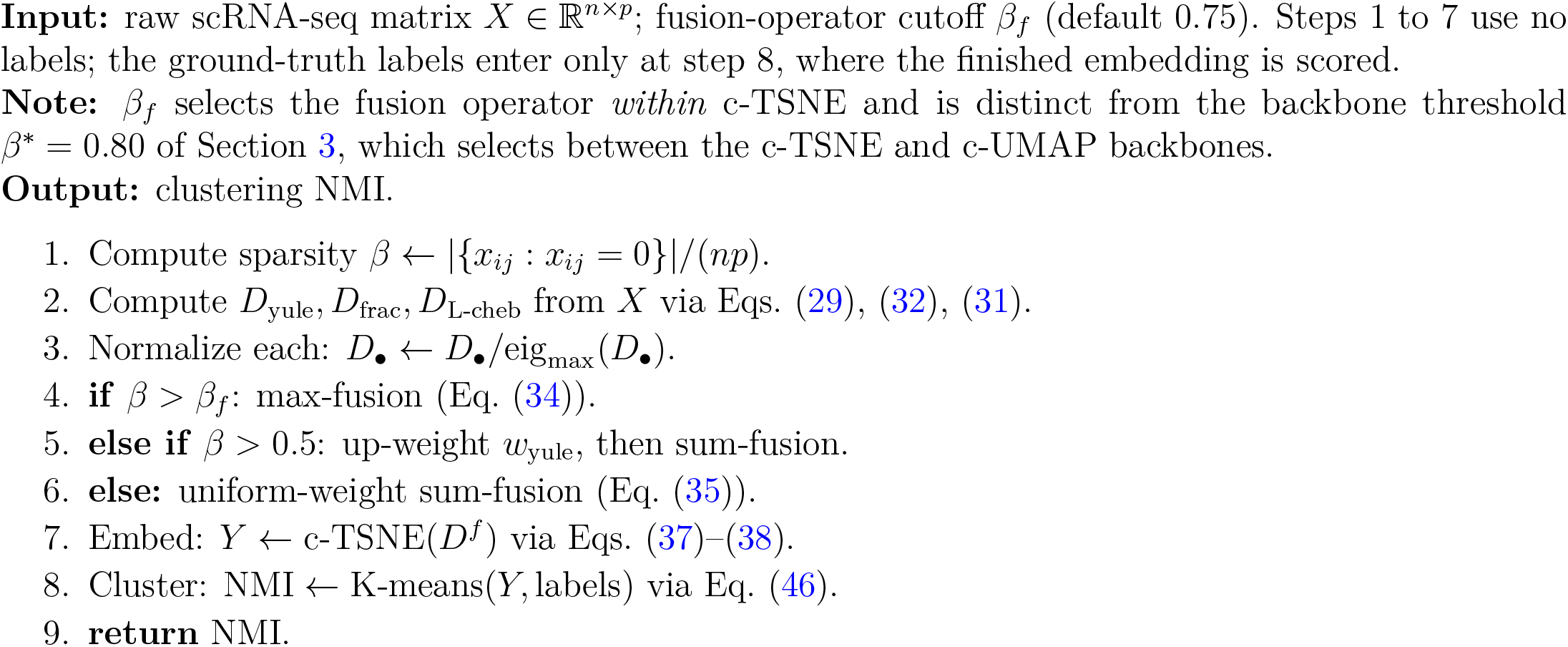

### 3.16. High-dimensional bias of the Euclidean metric

The following lemma says that in high dimensions all Euclidean distances crowd toward one value, so nearest and farthest neighbors stop being distinguishable.

#### Lemma 1.

*The Euclidean metric is biased for evaluating similarity in high-dimensional space: the relative spread of pairwise distances vanishes as dimension grows*.

*Proof*. Draw *n* points uniformly from the *p*-dimensional unit hypercube. For *a* = (*a*_1_, …, *a*_*p*_), *b* = (*b*_1_, …, *b*_*p*_), the squared distance is 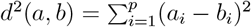. With *a*_*i*_, *b*_*i*_ ~ U[0, 1], **E**[*d*^2^(*a, b*)] = *p/*6 and **Var**[**E**[*d*^2^(*a, b*)] *p/*180, so the coefficient of variation is

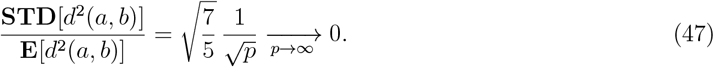

As *p* grows, pairwise distances concentrate at their mean and the observed range narrows like 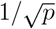; points become near-equidistant, erasing the distinction between nearest and farthest neighbors.

Table 5 shows this effect on real data. On each of the four benchmarks the Euclidean view trails the best cell-driven view. On Kolod the presence/absence view reaches 1.000 *±* 0.000 while the Euclidean view reaches 0.654 *±* 0.079.

## 4. Results

### Overview

We benchmarked c-TSNE against four widely used methods (t-SNE, UMAP, KPCA and SIMLR) on six well-characterized datasets spanning diverse cell types and sparsity 27.9–78.1%: Buettner, Kolod, Pollen, Usoskin, Patel, and Goolam. Normalized mutual information (NMI) is the primary metric. We additionally stress-test robustness to dropout and noise, challenges that remain under-explored in the single-cell literature. The c-TSNE workflow is two steps: (i) compute the c-TSNE embedding by fusing the three cell-driven dissimilarities; (ii) cluster the embedding with K-means (*k* set to the known number of classes, for comparability [4]). The same protocol is applied to every baseline method by substituting its embedding.

### 4.1. Baseline methods

We compare c-TSNE with t-SNE, UMAP, KPCA, and SIMLR [4, 6, 8, 9, 27]. The first three use Euclidean distance for cell similarity; in preliminary tests both UMAP and KPCA outperformed PCA on all four benchmarks, so we report the stronger pair. UMAP can be viewed as a manifold-learning variant of t-SNE addressing slow convergence [8]. For KPCA we use the cosine kernel *k*(*x, y*) = (*x* · *y*)*/*(∥*x*∥ ∥ *y*∥) for its automatic *L*_2_ normalization [9]. SIMLR learns a metric by multi-kernel integration and is a classic single-cell benchmark [4].

### 4.2. Benchmark datasets

Table 4 summarizes the four primary datasets; all values (cells, genes, classes, sparsity) were recomputed directly from the raw data.

**Table 4:** Primary benchmark scRNA-seq datasets (computed from raw data).

| Dataset | Cells | Genes | Classes | Sparsity (%) |
| --- | --- | --- | --- | --- |
| Buettner | 182 | 8,989 | 3 | 37.9 |
| Kolod | 704 | 10,685 | 3 | 27.9 |
| Pollen | 249 | 14,805 | 11 | 51.0 |
| Usoskin | 622 | 17,772 | 4 | 78.1 |

1. The **Buettner** dataset has 182 mouse embryonic stem cells (mESCs) in 3 cell-cycle stages (G1, S, G2M) [11].
2. The **Kolod** (Kolodziejczyk) dataset has 704 mESCs profiled under 3 culture conditions (2i, alternative-2i, serum), i.e. 3 classes [10].
3. The **Pollen** dataset has 249 cells across 11 populations (including skin, pluripotent stem, blood, and neural cells), designed to test low-coverage scRNA-seq for population identification [12].
4. The **Usoskin** dataset has 622 mouse dorsal-root-ganglion neurons in 4 neuronal types, ~1.14M reads/cell [13].

### 4.3. Cell-driven dissimilarities improve t-SNE visualization

The cell-driven views clearly outperform the default Euclidean metric in t-SNE visualization (Fig. 2). Each point is colored by its true label. Under Yule and L-Chebyshev, Buettner and Kolod show three well-separated clusters, while the Euclidean view keeps them closer together. The cell-driven views therefore give both a more accurate and a more explainable embedding. The advantage persists on Usoskin and Pollen, where the cell-driven views produce more separable clusters; the fractional view shows similar gains (Supplementary Material).

**Figure 2:**
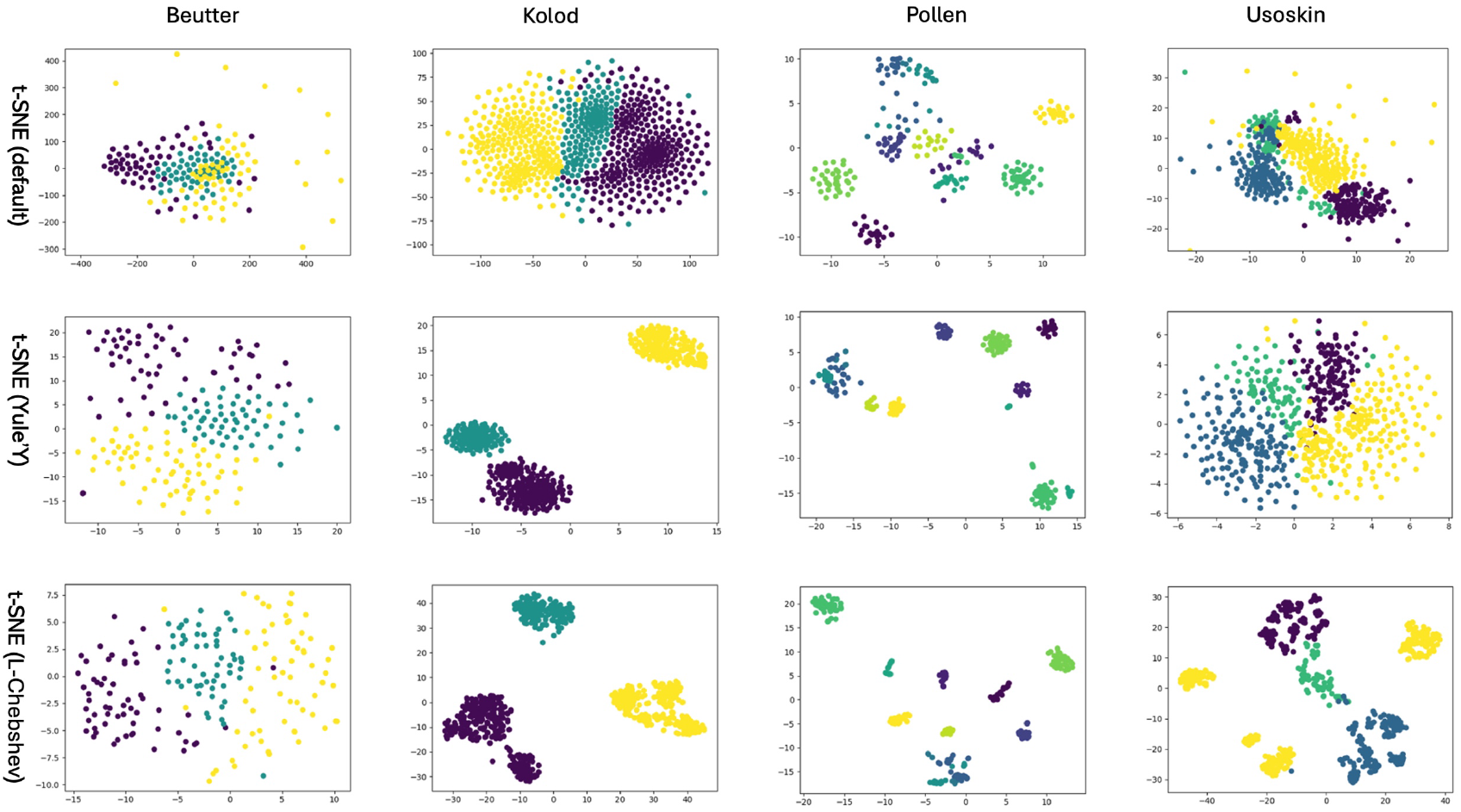
t-SNE visualizations under the Yule, L-Chebyshev, and (default) Euclidean dissimilarities on the four benchmark datasets (3, 3, 11, 4 clusters). The cell-driven views yield more meaningful subpopulation separation than Euclidean.

Table 5 quantifies this over 10 seeds. NMI lies in [0, 1] (Eq. (46)), and higher is better. On every dataset the best cell-driven view leads the Euclidean view. On Kolod, NMI reaches 1.000 *±* 0.000 under Yule against 0.654 *±* 0.079 for Euclidean, and on Buettner the fractional view reaches 0.824 *±* 0.013 against 0.303 *±* 0.010. This ordering is consistent with the high-dimensional behavior of Euclidean distance [5] established in Lemma 1. Which view leads moves with the data: Yule on Kolod, L-Chebyshev on Pollen and Usoskin, fractional on Buettner. That variation is precisely what motivates fusing them.

**Table 5:** NMI of t-SNE clustering under each dissimilarity, reported as mean *±*SD over 10 random seeds under a single fixed protocol. Best per row in bold.

| Dataset | Yule | L-Chebyshev | Euclidean | Fractional |
| --- | --- | --- | --- | --- |
| Buettner | 0.645 ± 0.070 | 0.572 ± 0.072 | 0.303 ± 0.010 | **0.824 ± 0.013** |
| Kolod | **1.000 ± 0.000** | 0.996 ± 0.005 | 0.654 ± 0.079 | 0.693 ± 0.032 |
| Pollen | 0.908 ± 0.010 | **0.954 ± 0.000** | 0.890 ± 0.015 | 0.935 ± 0.012 |
| Usoskin | 0.532 ± 0.081 | **0.779 ± 0.008** | 0.545 ± 0.034 | 0.540 ± 0.026 |

### 4.4. c-TSNE fusion improves visualization and clustering

c-TSNE visualizations are more representative than both classic t-SNE and t-SNE on any single cell-driven view (Fig. 3). Fusing the views yields more orthogonal, better-bounded clusters on all four datasets, showing that fusion, not any single source, drives the gain. Max-fusion is preferable for highly sparse data (e.g., the four Usoskin clusters at 78.1% sparsity), while sum-fusion is preferable at lower sparsity, foreshadowing the sparsity-adaptive rule.

**Figure 3:**
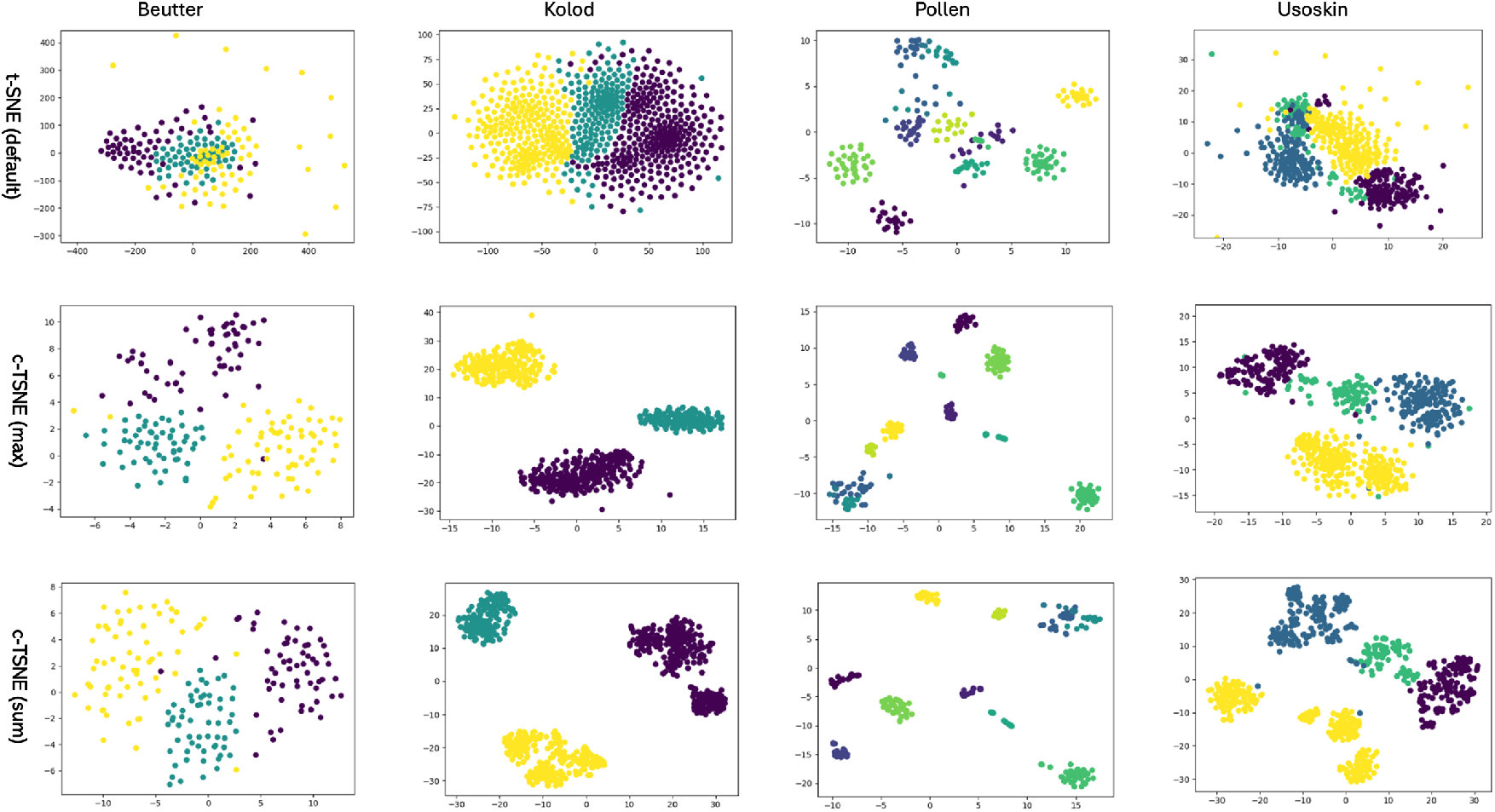
c-TSNE versus classic t-SNE (Euclidean) on the four benchmarks (3, 3, 11, 4 clusters). c-TSNE produces more separable clusters under both sum- and max-fusion, and also improves over t-SNE using any single cell-driven view.

Table 6 compares clustering NMI under the seeded protocol. Here c-TSNE runs Algorithm 1, which picks its fusion operator from sparsity alone with no reference to the score. c-TSNE leads every re-run baseline on all four datasets. It reaches 1.000 *±* 0.000 on Kolod and 0.863 *±* 0.015 on Usoskin, the sparsest of the four. The Euclidean-based methods vary widely across datasets, with KPCA at 0.896 on Kolod and 0.336 on Usoskin. SIMLR is the one broadly competitive baseline, and c-TSNE exceeds it on Kolod and Usoskin while remaining transparent and simple to implement, with no multi-kernel learning to tune. We attribute this consistency to fusing complementary, data-matched views rather than relying on a single dissimilarity.

**Table 6:** Clustering NMI of c-TSNE and its baselines, as mean *±* SD over 10 random seeds under a single fixed protocol. c-TSNE follows Algorithm 1, whose fusion operator is selected from sparsity alone. Best among the re-run methods in **bold**. ^*†*^SIMLR is carried over from the original single-run protocol and was not re-run here, so it is reported for reference and not bolded.

| Dataset | *c*-TSNE (Alg. 1) | t-SNE | KPCA | UMAP | SIMLR<sup>†</sup> |
| --- | --- | --- | --- | --- | --- |
| Buettner | **$0.821 \pm 0.032$** | $0.303 \pm 0.010$ | $0.556 \pm 0.000$ | $0.282 \pm 0.008$ | 0.89 |
| Kolod | **$1.000 \pm 0.000$** | $0.654 \pm 0.079$ | $0.896 \pm 0.000$ | $0.679 \pm 0.020$ | 0.99 |
| Pollen | **$0.918 \pm 0.021$** | $0.890 \pm 0.015$ | $0.725 \pm 0.003$ | $0.849 \pm 0.009$ | 0.95 |
| Usoskin | **$0.863 \pm 0.015$** | $0.545 \pm 0.034$ | $0.336 \pm 0.003$ | $0.556 \pm 0.014$ | 0.74 |

### 4.5. Entry-usage: a fusion-level explanation of heterogeneity

To open the fusion box, we measure the *entry-usage percentage*: under max-fusion, the fraction of entries of the final affinity contributed by each source. For *n* cells, the Yule entry-usage is

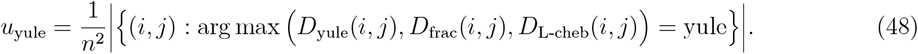

This reveals which biological factor dominates the fused distance and how sparsity shapes the fusion.

Table 7 shows a clear sparsity dependence. Kolod, the least sparse dataset, is dominated by L-Chebyshev (41.5%): when zeros are few, the magnitudes of top-expressed genes carry the most discriminative information. By contrast Usoskin, the sparsest dataset, is dominated by Yule (41.7%): when zeros abound, *whether* a gene is detected matters most. This directly motivates the fusion rule: prefer max-fusion (and up-weight Yule) for high-sparsity data, sum-fusion for low-sparsity data. The usage shares move with sparsity in the direction the rule assumes, across the full range of these datasets (Table 7). Entry-usage thus turns the fusion into an explanation: it states, per dataset, which biological axis the embedding relied on.

**Table 7:** Entry-usage (%) of each source in the max-fused distance.

| Dataset | $u_{\text{yule}}$ | $u_{\text{frac}}$ | $u_{\text{L-cheb}}$ |
| --- | --- | --- | --- |
| Buettner | 34.3 | 34.8 | 30.9 |
| Kolod | 25.6 | 32.9 | 41.5 |
| Pollen | 26.7 | 38.2 | 35.1 |
| Usoskin | 41.7 | 23.6 | 34.7 |

### 4.6. Robustness to dropout and noise

Robustness to dropout and noise is intrinsic to scRNA-seq yet rarely tested. Because c-TSNE down-weights transient, technical variation in favor of persistent, biologically meaningful signal, it should be more robust than classic t-SNE.

#### Dropout

We treat each dataset as a population and randomly remove 5%–50% of nonzero entries to simulate additional dropout, tracking NMI as sparsity rises.

#### Noise

We perturb each measurement *x*_*ij*_ with independent Gaussian noise *ϵ*_*ij*_ ~ *N* (0, *σ*_*ij*_), 0.1 ≤ *σ*_*ij*_ *<* 0.7, giving 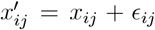. The range 0.1–0.7 stress-tests c-TSNE without fully erasing the signal. Gaussian noise is a standardized perturbation, not a model of the true expression distribution.

Across all datasets, c-TSNE substantially and consistently outperforms t-SNE in NMI under both perturbations (Fig. 4). On Kolod, c-TSNE’s NMI is nearly flat while t-SNE’s swings sharply; on Buettner under noise, c-TSNE’s NMI never drops below 0.70 while t-SNE’s never exceeds 0.30. We attribute this to the cell-driven fusion, which dampens noise by combining redundant views. Sum-fusion is generally more noise-robust, while max-fusion is advantageous for highly sparse data such as Usoskin.

**Figure 4:**
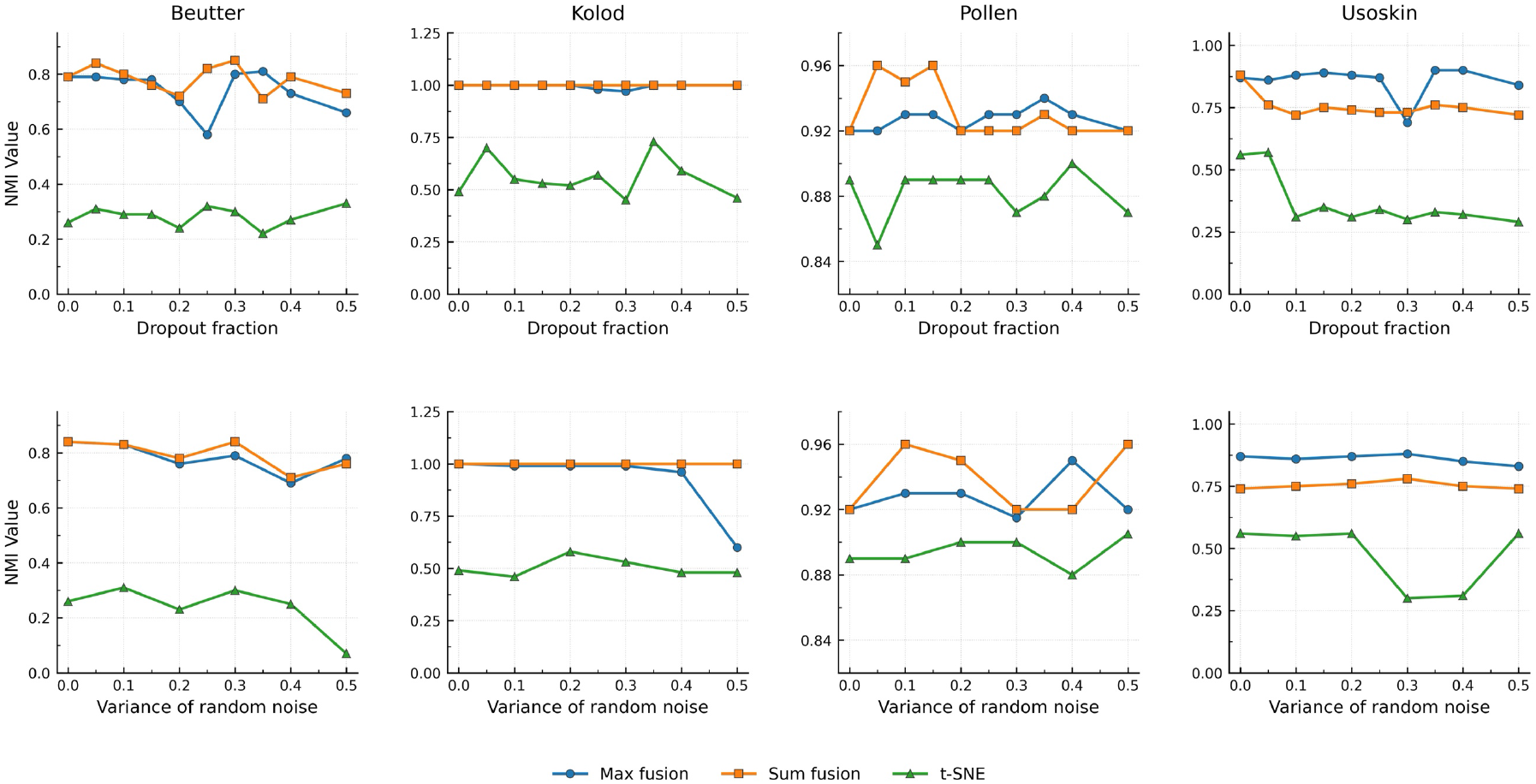
Robustness of c-TSNE versus classic t-SNE across dropout fractions and noise levels on Buettner, Kolod, Pollen, and Usoskin. c-TSNE maintains high, stable NMI; t-SNE fluctuates sharply, most visibly on the high-sparsity Usoskin. Under noise, sum-fusion is generally the more robust c-TSNE variant.

#### Statistical significance

We compared c-TSNE against t-SNE with the paired Wilcoxon signed-rank test over all (dataset *×* perturbation-level) points. Under dropout (*n* = 40), the sum-of-positive-ranks statistic is *W* ^+^ = 820 with *p* ≈ 1.8 *×* 10^−12^; under noise (*n* = 24), *W* ^+^ = 300 with *p* ≈ 1.2 *×* 10^−7^ (statistics recomputed from the released result tables). Both *p*-values are far below *α* = 0.05, so we reject the null and conclude that c-TSNE is significantly more robust than t-SNE to both dropout and noise. This robustness underpins its biological interpretability: the embeddings remain reliable under the technical perturbations typical of scRNA-seq.

### 4.7. Performance on additional datasets (Patel, Goolam)

We add two higher-sparsity benchmarks, Patel and Goolam (Table 8) [14, 15]. Goolam contains 124 blastomeres across 5 mouse-embryo stages (2-, 4-, 8-, 16-, 32-cell) [15]. Patel contains 543 cells from five primary glioblastomas, grouped into 5 transcriptional classes [14].

**Table 8:** Additional benchmark datasets (computed from raw data).

| Dataset | Cells | Genes | Classes | Sparsity (%) |
| --- | --- | --- | --- | --- |
| Patel | 543 | 5,948 | 5 | 53.3 |
| Goolam | 124 | 41,480 | 5 | 68.6 |

On Patel, Algorithm 1 reads a sparsity of 53.3% and up-weights the presence/absence view in the sum operator, using *w*_yule_ = 0.8, *w*_frac_ = 0.1, and *w*_L-cheb_ = 0.1 (Fig. 5). This gives NMI 0.771 *±* 0.009, ahead of every Euclidean-geometry baseline (Table 9). The presence/absence view on its own reaches 0.876 *±* 0.006 here, which is the behavior the sparsity-adaptive rule anticipates: as sparsity rises, gene detection becomes the most informative axis and the operator concentrates weight on it.

**Figure 5:**
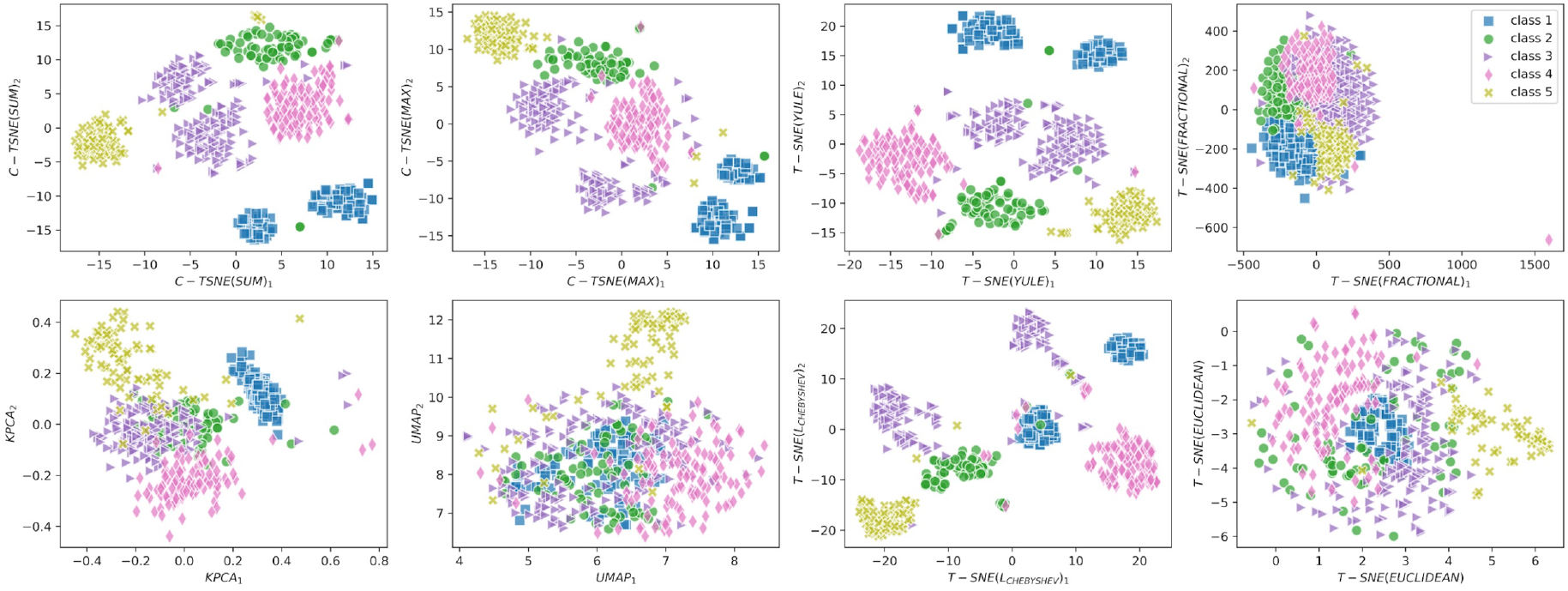
c-TSNE versus baselines on Patel: t-SNE under the individual cell-driven and Euclidean dissimilarities, KPCA, and UMAP. For this high-sparsity dataset, c-TSNE (sum- and max-fusion) and t-SNE-Yule separate the five clusters best.

**Table 9:** Clustering NMI on Patel and Goolam, as mean *±* SD over 10 random seeds under a single fixed protocol. c-TSNE follows Algorithm 1. Best among the re-run methods in bold. ^*†*^SIMLR is carried over from the original single-run protocol.

| Dataset | c-TSNE | t-SNE under a single view |  |  |  |  |  |
| --- | --- | --- | --- | --- | --- | --- | --- |
|  | (Alg. 1) | Yule | frac. | L-cheb | Eucl. | KPCA | UMAP |
| Patel | $0.771 \pm 0.009$ | **$0.876 \pm 0.006$** | $0.583 \pm 0.076$ | $0.623 \pm 0.024$ | $0.329 \pm 0.030$ | $0.565 \pm 0.003$ | $0.228 \pm 0.019$ |
| Goolam | $0.660 \pm 0.011$ | **$0.776 \pm 0.000$** | $0.586 \pm 0.047$ | $0.258 \pm 0.007$ | $0.241 \pm 0.006$ | $0.302 \pm 0.004$ | $0.269 \pm 0.022$ |

SIMLR<sup>†</sup>: Patel 0.82, Goolam 0.69.

The same pattern holds on Goolam at 68.6% sparsity (Fig. 6). Algorithm 1 reaches 0.660 *±* 0.011, ahead of every Euclidean-geometry baseline, while the presence/absence view alone reaches 0.776 *±* 0.000. Both datasets support the recommendation of Section 6: give the presence/absence view dominant weight once sparsity is high.

**Figure 6:**
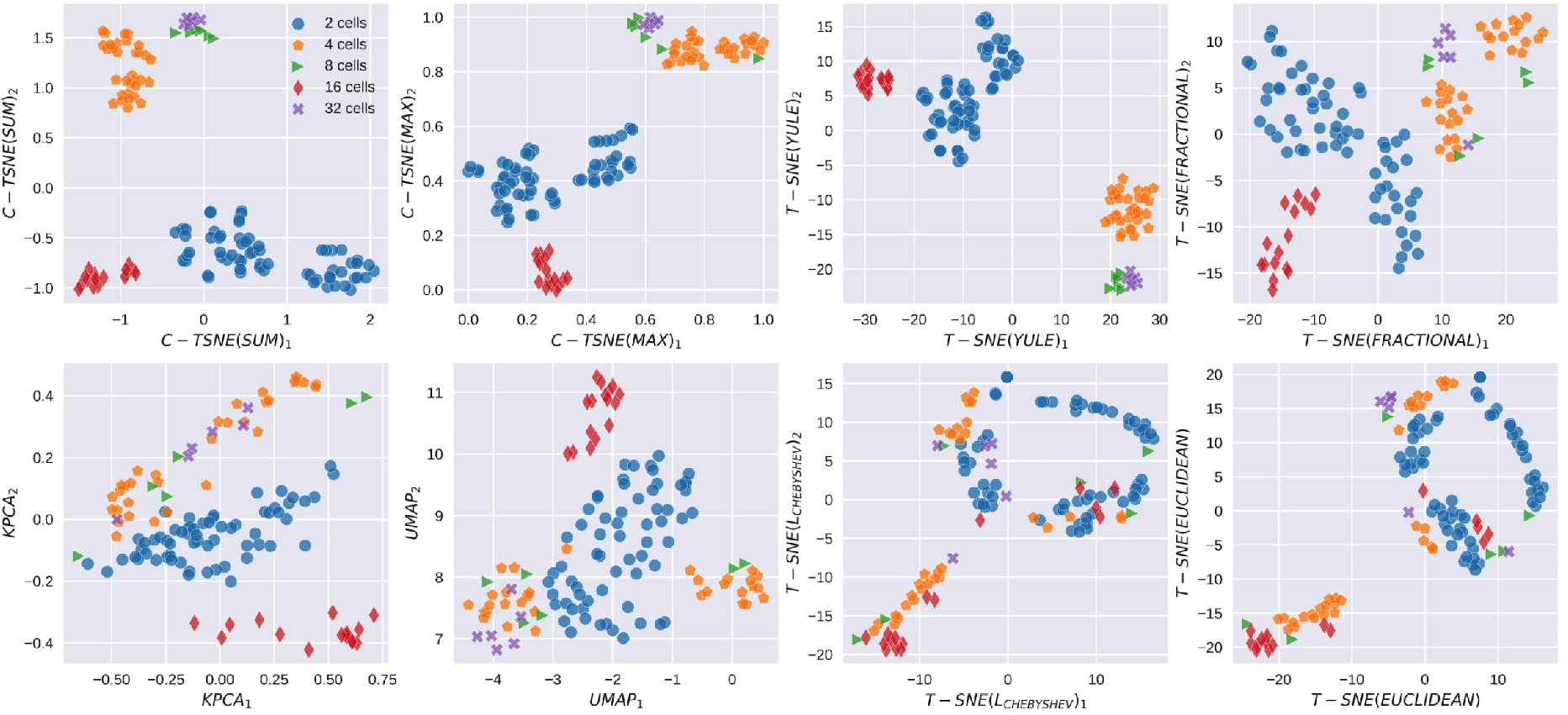
c-TSNE versus baselines on Goolam: t-SNE under the three cell-driven and default dissimilarities, KPCA, and UMAP. c-TSNE (sum-fusion) and t-SNE under the Yule view give the clearest separations.

Across all six benchmarks, c-TSNE under Algorithm 1 leads every Euclidean-geometry baseline, and its margin over t-SNE with Euclidean distance is consistent throughout.

### 4.8. Atlas-scale validation: 29 datasets, 11 baselines, two protocols

The second part of our evaluation moves from the plate-based benchmarks of Sections 4.3–4.7 to the regime single-cell studies actually face today: **large, highly sparse atlases**. We assembled **29 publicly available scRNA-seq datasets**, each processed under two protocols (Raw and PP, below) and the two cortex datasets read additionally at fine label resolution, giving **62 evaluation entries**, and compared c-UMAP against **11 baselines** under **two metrics** (NMI and AMI). The collection spans **225 to 1**,**136**,**218 cells**, a 5,000-fold range; 11,684 to 61,497 genes; **3 to 180** ground-truth classes; and **45.0% to 98.0%** sparsity. It covers two species, twelve source studies, and three protocol families: plate-based (Fluidigm C1, Smart-seq2), droplet-based (Drop-seq, inDrop, 10x, DroNc-seq, sNuc-DropSeq) and microwell-based (Microwell-seq). Table 10 lists every dataset, grouped by family. Supplementary Tables S1 to S8 and the 56 per-dataset embedding figures referenced throughout this section are contained in the accompanying appendix document, which is submitted as Supplementary Material; the full per-dataset catalogue is Table C.20.

**Table 10:** The 29-dataset benchmark collection, dataset by dataset. Cells, genes and sparsity are measured on the raw count matrix; *C* is the number of ground-truth classes. Families are separated by rules. The two cortex datasets are additionally read at fine label resolution (*C* = 93 and *C* = 81), giving the 62 evaluation entries of Section 4.10.

| Family | Dataset | Cells | Genes | $C$ | Sparsity (%) |
| --- | --- | --- | --- | --- | --- |
| Cell lines and stem cells | Tian (Drop-seq) | 225 | 15,127 | 3 | 62.1 |
|  | Kumar | 246 | 45,159 | 3 | 45.8 |
|  | Koh | 531 | 48,981 | 9 | 72.1 |
|  | Tian (10x) | 902 | 16,468 | 3 | 45.0 |
|  | Chu | 1,018 | 19,097 | 7 | 49.8 |
|  | Tian (10x, 5 lines) | 3,918 | 11,786 | 5 | 63.0 |
| Pancreatic islets | Baron mouse 1 | 822 | 14,878 | 13 | 90.5 |
|  | Baron mouse 2 | 1,064 | 14,878 | 13 | 87.8 |
|  | Baron human 4 | 1,303 | 20,125 | 14 | 89.0 |
|  | Baron human 2 | 1,724 | 20,125 | 14 | 90.6 |
|  | Baron human 1 | 1,937 | 20,125 | 14 | 90.4 |
|  | Enge | 2,282 | 23,460 | 6 | 86.0 |
|  | Baron human 3 | 3,605 | 20,125 | 14 | 91.3 |
| Tissue atlases | Wu kidney (DroNc-seq) | 2,769 | 19,713 | 13 | 94.3 |
|  | Wu kidney (sNuc-DropSeq) | 3,011 | 19,713 | 13 | 95.4 |
|  | Duodenum | 5,200 | 30,172 | 10 | 91.2 |
|  | MCA fetal intestine | 6,076 | 20,115 | 13 | 96.7 |
|  | MCA fetal stomach | 6,192 | 19,720 | 16 | 97.4 |
|  | HCL peripheral blood | 7,160 | 21,267 | 13 | 98.0 |
|  | HCL muscle | 7,775 | 22,961 | 17 | 97.9 |
|  | HCL adrenal gland | 8,114 | 23,842 | 13 | 97.7 |
|  | HCL kidney | 9,153 | 25,601 | 18 | 97.5 |
|  | HCL trachea | 9,949 | 24,990 | 14 | 97.1 |
|  | Lung | 10,360 | 16,327 | 13 | 96.6 |
| Large atlases | ZhengMix (PBMC) | 11,749 | 11,684 | 8 | 95.3 |
|  | DropViz cortex posterior | 46,346 | 27,720 | 14 | 93.5 |
|  | DropViz cortex frontal | 71,639 | 29,463 | 14 | 93.2 |
|  | Tabula Sapiens (0.5M) | 508,731 | 60,606 | 42 | 95.5 |
|  | Tabula Sapiens (1.1M) | 1,136,218 | 61,497 | 180 | 94.8 |
| **Total** | **29 datasets** | **225–1,136,218** | **11,684–61,497** | **3–180** | **45.0–98.0** |

#### How this evaluation is designed

Four choices govern the evaluation, each made to test the fusion rule rather than to present it favorably. First, the weights are fixed once per protocol and never adjusted per dataset, so a win cannot come from tuning (Section 4.9). Second, the collection spans a 5,000-fold range in cell count, which tests whether one rule holds from a plate of 225 cells to a million-cell atlas. Third, every comparison is scored by two indices, since NMI alone rewards large cluster counts and AMI corrects for that. Fourth, we measure what the preprocessing we avoid actually removes (Section 4.12), so the preprocessing-free design is tested rather than asserted. Readers who wish to check any single number will find the per-dataset tables in Tables C.21–C.24.

#### Baseline methods

We compare against **11** baselines spanning four families. *Classical dimension reduction:* t-SNE [6], UMAP [8], kernel PCA [9] and PHATE [32]. *Similarity learning:* SIMLR [4]. *Deep learning:* scDCCA [34], scGAC [35] and scAGCL [47]. *Consensus and pretrained models:* scICE [33] and the single-cell foundation model scGPT [46], clustered both by K-means (scGPT-K) and by Leiden (scGPT-L). Every baseline runs at its published defaults, and every method that requires a cluster count receives the true *C*.

#### Two protocols

**Raw**: all methods, c-UMAP included, receive the same unprocessed count matrix. **PP**: each baseline receives a standard preprocessing pipeline (quality filtering, normalization, log-transform, 2,000 highly-variable-gene selection, scaling), while c-UMAP continues to operate on raw counts. Reporting both protocols lets the reader separate the contribution of the fusion from the contribution of preprocessing, and Section 4.12 quantifies what the PP pipeline removes.

### 4.9. One fixed fusion weight setting across all 29 datasets

A fusion rule is only useful if it transfers. We therefore fix the weights once per protocol. The same values are then applied unchanged to all 29 datasets, rather than tuning (*w*_*y*_, *w*_*f*_, *w*_*c*_) for each one.

| Protocol | Fixed weights $[w_{\text{yule}}, w_{\text{frac}}, w_{\text{L-cheb}}]$ |
| --- | --- |
| Raw (no preprocessing) | [0.90, 0.00, 0.10] |
| PP (standard preprocessing) | [0.80, 0.10, 0.10] |

The presence/absence view carries most of the weight in both protocols, which is what Section 4.5 predicts for data reaching 98% sparsity. The weight assigned to each view tracks sparsity, and this is the sparsity-adaptive principle of Section 3.6 acting on real data. The holistic view is weighted most heavily at moderate sparsity, where it is the single strongest view on Buettner (Table 5), and it retains weight under PP. Under Raw, at the high sparsity of this collection, the detection signal is the most informative axis and the weight concentrates there accordingly. The holistic view keeps a share in the degenerate regime described next, where it is weighted at 0.20 against 0.00 for detection. A view therefore earns weight in the sparsity regime it was designed to describe, rather than in every regime at once.

One explicit, label-free safeguard covers the degenerate case. Yule’s coefficient is exactly zero whenever one cell’s set of detected genes is nested inside the other’s. It therefore stops ordering cells once feature selection leaves too few genes per cell. We therefore monitor *z*_15_, the fraction of cells having at least 15 partners at Yule distance zero, and switch to [0.00, 0.20, 0.80] when *z*_15_ *>* 0.8. Across all 62 entries exactly one triggers this rule: the Tabula Sapiens [48] full atlas under PP (*z*_15_ = 1.000, against 0.000 for the other 61 entries), where highly-variable-gene selection leaves a median of 21 detected genes per cell. The same two weight vectors govern datasets differing 5,000-fold in size, from 3 to 180 classes, and across three protocol families. The performance reported below therefore reflects the fusion principle itself, not per-dataset tuning.

#### Clustering

c-UMAP clusters the fused graph with Leiden under the CPM objective; K-means on the two-dimensional embedding is used when a dataset has at least 6,000 cells and at most 10 classes. Leiden takes a resolution rather than a cluster count. We therefore sweep a geometric grid and record the number of clusters each resolution yields. The resolution whose count comes closest to *C* is kept, with ties broken toward the larger resolution.

### 4.10. Clustering accuracy across 62 entries

**c-UMAP attains the highest mean score in all four settings**, covering both indices under both protocols (Table 11). It is ahead of every one of the 11 baselines on both indices. One two-sided Wilcoxon signed-rank test was run per baseline per index, giving 22 tests in total. Every test returns *p <* 0.001, and every test stays significant after Bonferroni correction. The per-dataset tables (NMI and AMI, Raw and PP) are Tables C.21–C.24.

**Table 11:** Summary over the 29-dataset benchmark. Columns 2–5 are mean scores across datasets within each protocol (higher is better). “Wins” and *p* are computed on Raw and PP pooled; the denominator is the number of paired entries the baseline produced, since entries a baseline could not produce are dropped from its test rather than scored as zero. All 22 tests are significant after Bonferroni correction. Best per column in **bold**.

| Method | Mean NMI |  | Mean AMI |  | NMI vs. c-UMAP |  | AMI vs. c-UMAP |  |
| --- | --- | --- | --- | --- | --- | --- | --- | --- |
| | Raw | PP | Raw | PP | Wins | $p$ | Wins | $p$ |
| **c-UMAP** | **0.618** | **0.696** | **0.657** | **0.728** | n/a | n/a | n/a | n/a |
| scICE | 0.499 | 0.632 | 0.573 | 0.707 | 42/57 | $2.5 \times 10^{-5}$ | 39/57 | $9.5 \times 10^{-4}$ |
| scAGCL | 0.579 | 0.579 | 0.618 | 0.626 | 45/54 | $1.5 \times 10^{-6}$ | 47/54 | $2.0 \times 10^{-7}$ |
| UMAP | 0.473 | 0.646 | 0.509 | 0.695 | 45/58 | $8.7 \times 10^{-8}$ | 48/58 | $3.1 \times 10^{-8}$ |
| PHATE | 0.402 | 0.661 | 0.430 | 0.697 | 46/58 | $1.3 \times 10^{-7}$ | 47/58 | $2.6 \times 10^{-8}$ |
| t-SNE | 0.464 | 0.563 | 0.502 | 0.608 | 48/58 | $9.0 \times 10^{-9}$ | 50/58 | $2.0 \times 10^{-9}$ |
| scGPT-L | 0.490 | 0.514 | 0.543 | 0.573 | 47/59 | $2.6 \times 10^{-6}$ | 46/59 | $1.6 \times 10^{-6}$ |
| scGPT-K | 0.449 | 0.480 | 0.489 | 0.521 | 49/59 | $4.7 \times 10^{-9}$ | 52/59 | $1.7 \times 10^{-9}$ |
| scDCCA | 0.363 | 0.594 | 0.381 | 0.633 | 52/58 | $7.1 \times 10^{-9}$ | 53/58 | $1.8 \times 10^{-9}$ |
| scGAC | 0.378 | 0.505 | 0.413 | 0.557 | 39/46 | $8.3 \times 10^{-8}$ | 40/46 | $4.7 \times 10^{-8}$ |
| SIMLR | 0.365 | 0.581 | 0.378 | 0.597 | 52/55 | $1.2 \times 10^{-9}$ | 53/55 | $5.6 \times 10^{-10}$ |
| KPCA | 0.163 | 0.521 | 0.168 | 0.561 | 55/58 | $6.7 \times 10^{-11}$ | 55/58 | $6.4 \times 10^{-11}$ |

#### Reading the two protocols together

The lead holds whether or not the baselines are given their preferred input. Under **Raw**, where every method sees the same unprocessed counts, c-UMAP’s mean NMI of 0.618 leads the field, and the margin over the Euclidean-geometry methods is large (t-SNE 0.464, UMAP 0.473, KPCA 0.163). Under **PP**, where the baselines receive their recommended pipelines and c-UMAP still reads raw counts, c-UMAP’s mean NMI rises to 0.696 and remains the highest, as does its mean AMI of 0.728. The cell-driven fusion therefore delivers on its intended input without requiring the normalization, log-transform and gene-selection steps its competitors depend on. AMI, which corrects the chance agreement that inflates NMI at large *K*, ranks the methods the same way, so the conclusion does not depend on the choice of index.

#### A reference that no pipeline produced

Both indices score a clustering against a reference annotation, and most published annotations were themselves produced by a standard pipeline that begins with highly variable gene selection. Section 4.12 shows what that step removes, so a reader is entitled to ask whether the comparison rewards agreement with the very pipeline this work argues against. One dataset settles the question on its own terms. ZhengMix is assembled from populations separated by fluorescence-activated cell sorting. Its eight labels were therefore fixed by a physical assay before any sequencing. No computational pipeline of any kind stands between the cells and their classes. Under Raw, c-UMAP reaches NMI 0.756 there against 0.723 for the strongest baseline, scGPT with Leiden, and separates all eight immune populations from unprocessed counts (Figure 10). The ordering on that dataset is the same as the ordering over the benchmark as a whole, so the advantage does not depend on labels that a preprocessing pipeline helped to draw.

#### Resolution of the ground truth

Two further readings, Cortex P (*C* = 93) and Cortex F (*C* = 81), score the two cortex datasets against the fine-grained DropViz sub-clusters rather than the 14 global classes. These are the same cells read at a higher label resolution, and c-UMAP retains its advantage there (NMI 0.542 and 0.536 under Raw), showing the gain is not an artifact of a coarse label set.

### 4.11. Scalability: reaching 1.14 million cells

Scalability is part of the method, not an afterthought: real atlases hold 10^5^ to 10^6^ cells. Complexity must be read over the whole pipeline rather than over the layer a method emphasizes. Table 12 gives the overall complexity of each method it lists and the step that dominates it.

**Table 12:** Overall complexity and dominating step. c-UMAP’s lean implementation contains no *N × N* step at any point, which is what lets it reach 1.14 *×* 10^6^ cells.

| Method | Overall complexity | Dominating step |
| --- | --- | --- |
| t-SNE (BH/FFT) | $O(n \log n)$ | gradient iterations |
| UMAP | $\approx O(n^{1.14})$ | $k$-NN graph |
| PHATE | $O(n^2)$ | diffusion operator |
| KernelPCA | $O(n^2)$–$O(n^3)$ | $N \times N$ kernel |
| SIMLR | $O(n^2)$ | $N \times N$ similarity |
| scGAC | $O(n^2)$ | $N \times N$ attention |
| scDCCA | $O(n)$ per epoch | ZINB autoencoder |
| scAGCL | $O(n^2)$ | pairwise graph construction |
| scICE | $O(n^2)$ overall | scLENS cell×cell eigendecomposition |
| c-UMAP (dense) | $O(n^2)$ | pairwise distances |
| **c-UMAP (lean)** | $\approx O(nk)$ | approximate $k$-NN; **no $N \times N$ step** |

#### Three implementations, one fusion

The same three distances and the same weights are used at every scale; only the way they are computed changes. Below 10^4^ cells all three distance matrices are formed densely and fused at the distance level. Between 10^4^ and 10^5^ cells they are precomputed and cached, or, under PP, replaced by three UMAP graphs built on 100 principal components and fused at the graph level. Above 10^5^ cells a dense matrix is no longer possible, because 5 *×* 10^5^ cells would already require 1 TB of memory. Neighbors are therefore found by approximate *k*-NN on the GPU. The switch is forced by memory, and the fusion weights are identical before and after it.

#### Measured wall-clock time

c-UMAP runs faster than the strongest baseline at every size tested, and the margin widens as datasets grow. Table 13 reports end-to-end time for c-UMAP and for scICE on the same machine and configuration, where end to end means graph construction together with clustering. The speed-up is 2 to 7*×* on the pancreas sets and 9*×* on cortex frontal. On ZhengMix it reaches 22*×*. At 508,731 cells c-UMAP finishes in 2.4 hours. We re-measured every entry end to end rather than quoting published figures. The reason is visible on ZhengMix under Raw: scICE’s clustering step takes 440 s while building its graph takes 5,297 s, so graph construction is 92% of the total.

**Table 13:** End-to-end wall-clock time (graph construction plus clustering) for c-UMAP and scICE on identical hardware. ^*†*^For the cortex rows only scICE’s graph-construction step could be timed, so those speed-ups are lower bounds.

| Dataset | Cells | c-UMAP | scICE | Speed-up |
| --- | --- | --- | --- | --- |
| *Raw (no preprocessing)* |  |  |  |  |
| H1 | 1,937 | 69.0 s | 136 s | $2.0\times$ |
| H3 | 3,605 | 92.5 s | 185 s | $2.0\times$ |
| ZhengMix | 11,749 | 482 s | 5,737 s | **$11.9\times$** |
| Tabula Sapiens | 508,731 | **2.4 h** | n/a | n/a |
| *PP (standard preprocessing)* |  |  |  |  |
| H1 | 1,925 | 19.2 s | 116 s | $6.0\times$ |
| H3 | 3,600 | 31.7 s | 222 s | $7.0\times$ |
| ZhengMix | 11,749 | 75.4 s | 1,677 s | **$22.2\times$** |
| Cortex posterior | 46,341 | 699 s | 3,050 s<sup>†</sup> | $4.4\times$ |
| Cortex frontal | 71,594 | 1,076 s | 9,685 s<sup>†</sup> | $9.0\times$ |

#### Measured scaling exponent

The measured cost of c-UMAP grows almost linearly in the number of cells. Because a complexity table is an argument rather than a measurement, we fit the exponent directly. Cortex frontal was subsampled to a series of sizes with dimension and class structure held fixed, so that only *n* varies. The fit is *t* ∝ *n*^1.01^ under PP across five sizes from 5k to 71.6k cells, and *t*∝ *n*^1.07^ under Raw. This is a within-dataset fit; fitting across datasets, where dimension, sparsity and class count all vary at once, gives about *n*^1.35^. The two numbers measure different things and should not be quoted interchangeably.

#### Reaching the million-cell scale

c-UMAP produces embeddings and clusterings for both Tabula Sapiens atlases: **508**,**731 cells with 42 classes** and **1**,**136**,**218 cells with 180 classes**, attaining NMI 0.590 and 0.677 and AMI 0.622 and 0.684 under Raw. This is the largest scale in the comparison. c-UMAP reaches it with the same fusion and the same weights used on the 225-cell Tian Drop-seq set, and Figure 7 shows both atlas embeddings, the 508,731-cell immune compartment in Figure 7a and the 1,136,218-cell full atlas in Figure 7b. At this scale the claim is feasibility rather than a margin, because no baseline in the comparison produces an embedding there at all.

**Figure 7:**
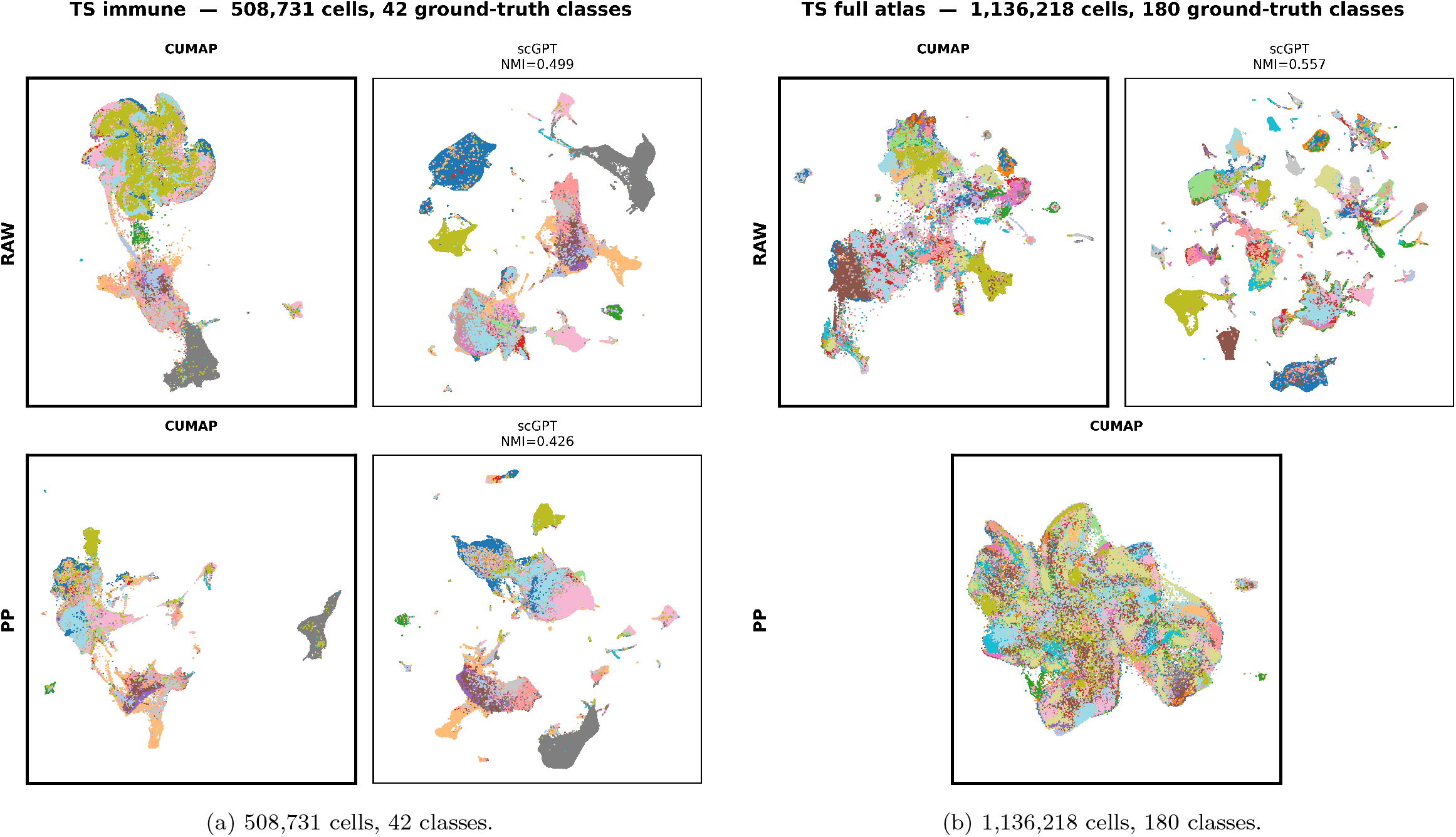
Atlas scale, Raw protocol. c-UMAP reads the unprocessed count matrix of the two Tabula Sapiens atlases under the same fused distance and the same fixed weights applied to the 225-cell Tian Drop-seq benchmark. Point colour is the published cell-type label. The two panels stand alone because no baseline in the comparison produces an embedding at either scale.

The reason is architectural rather than incidental, and it differs by family. The similarity-learning and kernel methods are bounded by an explicit *n × n* object: SIMLR forms a dense multiple-kernel similarity matrix. At 508k cells that is 1,913 GB of doubles against the 502 GB available. Exact kernel PCA needs the 508,731^2^ RBF kernel, about 2 TB. The graph-attention model fails one level up for the same reason: scGAC materializes an *N × N* attention matrix. Attention over cells therefore costs what a dense similarity costs. Minibatching over cells does not avoid that while the attention stays global. scAGCL, which contrasts augmented views of one graph, exhausts memory at the million-cell scale and hits our two-hour cap on the cortex atlases. The deep generative model fails differently: scDCCA’s zero-inflated negative-binomial likelihood went numerically undefined in the first epoch on the full atlas and stayed so for all 100 epochs. Every cell collapsed into one cluster. The obstacle there is the loss, not the memory. The classical methods are bounded by their own exact form, *O*(*n*^2^) for t-SNE, and reach this scale only through GPU or landmark approximations that are no longer the method being compared. One further baseline, scICE, is bounded by single-core graph construction: its 508k-cell run produced nothing after seven days.

Low published complexity is therefore an incomplete guide to what a pipeline can run, and the per-entry reasons are recorded with Tables C.21–C.24. c-UMAP reaches the million-cell scale because its lean path builds no *N × N* object at any step: the fusion is evaluated only on the edges of an approximate *k*-nearest-neighbour graph.

### 4.12. What preprocessing discards: the case for a preprocessing-free method

The framework’s preprocessing-free design is motivated by a measurable claim, which we test directly. The standard pipeline keeps 2,000 genes out of tens of thousands, and makes that choice from variance alone without ever seeing the labels. The statistical basis of such selection, and of the normalizations that accompany it, has been examined at length [1, 17]. We quantified how many *label-informative* genes it drops. Three criteria are used, each averaged over the ground-truth classes of a dataset.

A. **DE genes lost**. For each class we take its 50 top-ranked differentially expressed genes, ranked by a Wilcoxon rank-sum test on the raw matrix. We report the percentage not among the 2,000.
B. **Marker genes lost**. For each class we take the union of its curated markers in PanglaoDB [44] and CellMarker 2.0 [45]. We report the percentage not among the 2,000.
C. **Intersection lost**. We report the percentage of the DE-confirmed markers, meaning the genes counted by both A and B, that are not among the 2,000.

Across the 29 datasets, criterion A has a median of **46.9%** (range 20.7–94.7%), criterion B ranges 44.7–88.7%, and criterion C ranges 10.1% on Pancreas H3 to 83.3% on the Tabula Sapiens full atlas. In total the selection drops **15**,**121** (dataset, class, gene) instances that rank among the top differentially expressed genes of their class. These come from **4**,**988 distinct genes**. Of those instances, **2**,**032** are also a curated marker of the same class, covering **970 distinct genes**. Per-dataset values are in Table C.26.

Table 14 makes the point concrete. Each listed gene is both a top-ranked differentially expressed gene of its class *and* a curated marker of that same class, and each is discarded before any method sees the data.

**Table 14:** Canonical marker genes discarded by variance-based feature selection. Each is a top-ranked DE gene of its class and a curated marker of the same class. “Rank” is its position among that class’s DE genes. A fuller list is in Table C.27.

| Gene | Dataset | Class | Role | Rank |
| --- | --- | --- | --- | --- |
| CTRB1 | Pancreas H3 | acinar | acinar digestive-enzyme marker | 1 |
| IAPP | Pancreas H2 | beta | amylin, co-secreted with insulin | 2 |
| GCG | Pancreas H4 | alpha | proglucagon, defining alpha-cell hormone | 2 |
| CD3D | ZhengMix | memory T | delta subunit of the TCR-CD3 complex | 9 |
| PLP1 | Cortex frontal | oligodendrocyte | main structural protein of myelin | 1 |
| SLC1A2 | Cortex frontal | astrocyte | principal astrocytic glutamate transporter | 8 |
| MS4A1 | TS immune | B cell | CD20, defining B-cell surface antigen | 3 |
| FOXP3 | TS immune | regulatory T cell | master transcription factor of Tregs | 3 |
| FCGR3A | TS immune | non-classical monocyte | CD16, defines non-classical monocytes | 1 |

A pipeline that removes *IAPP* before clustering beta cells, or *FOXP3* before resolving regulatory T cells, is discarding precisely the evidence a biologist would use to name the resulting cluster. This is the concrete cost that the preprocessing-free design avoids: by reading raw counts, c-UMAP keeps every marker gene available to the fusion, and the entry-usage diagnostic of Section 4.5 can then attribute each cluster boundary to a biological axis computed over the complete transcriptome.

### 4.13. Embeddings

Scores alone do not show what a method does to the data. Figures 8–11 place every method side by side on the two largest non-atlas benchmarks under both protocols. Point color gives the ground-truth cell type, and the coloring is identical across panels, so they can be read against one another. Compact, well-separated blocks of one color indicate recovery of the true cell types. Figure 12 adds a smaller, densely annotated example in the human-pancreas donor H1, where the endocrine and exocrine populations separate into distinct groups. A representative subset is shown here; the full set for all 29 datasets under both protocols is released with the code.

**Figure 8:**
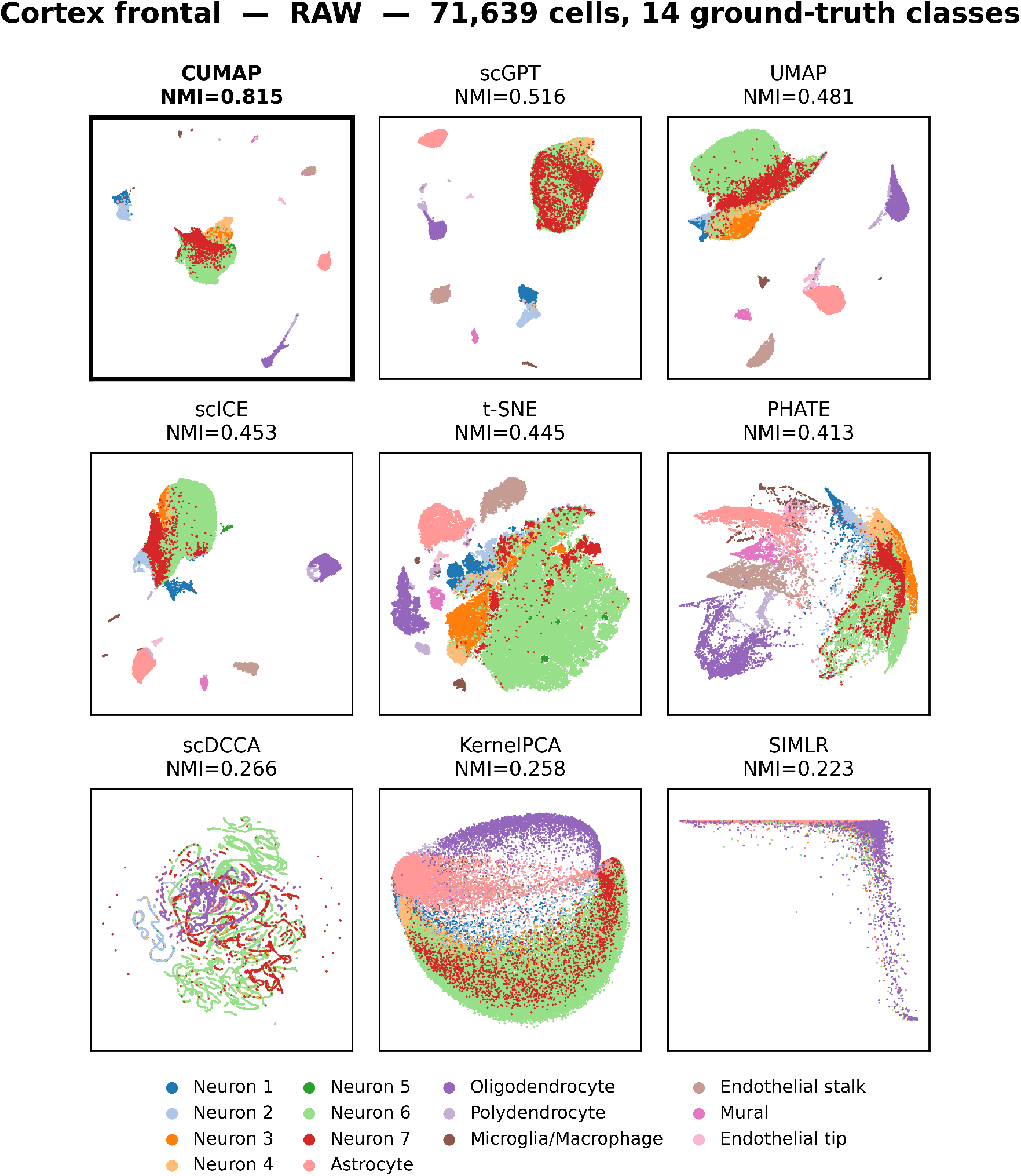
Cortex frontal, Raw protocol. (71,639 cells, 14 ground-truth classes). Every method receives the same unprocessed count matrix. c-UMAP resolves the cortical cell types into compact, separated groups while operating directly on raw counts.

**Figure 9:**
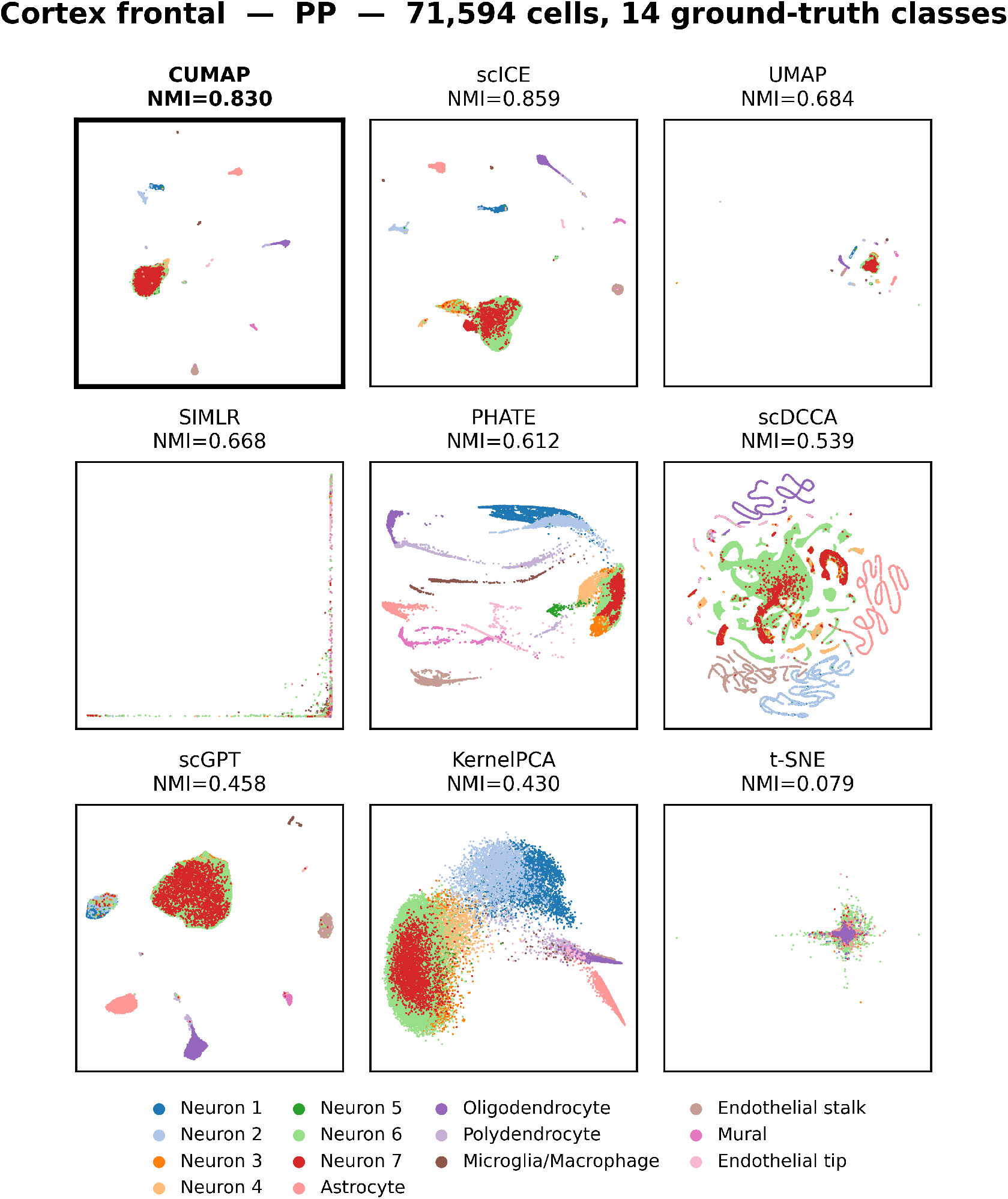
Cortex frontal, PP protocol. (71,594 cells, 14 ground-truth classes). Each baseline runs under its own recommended preprocessing pipeline, while c-UMAP continues to read raw counts and retains compact, well-separated clusters. This is the one panel in which a baseline scores higher: scICE reaches 0.859 against c-UMAP’s 0.830 once scICE is given its preferred input. The box marks the cell-driven method, not the highest score. Over the benchmark as a whole c-UMAP leads under PP as well (Table 11).

**Figure 10:**
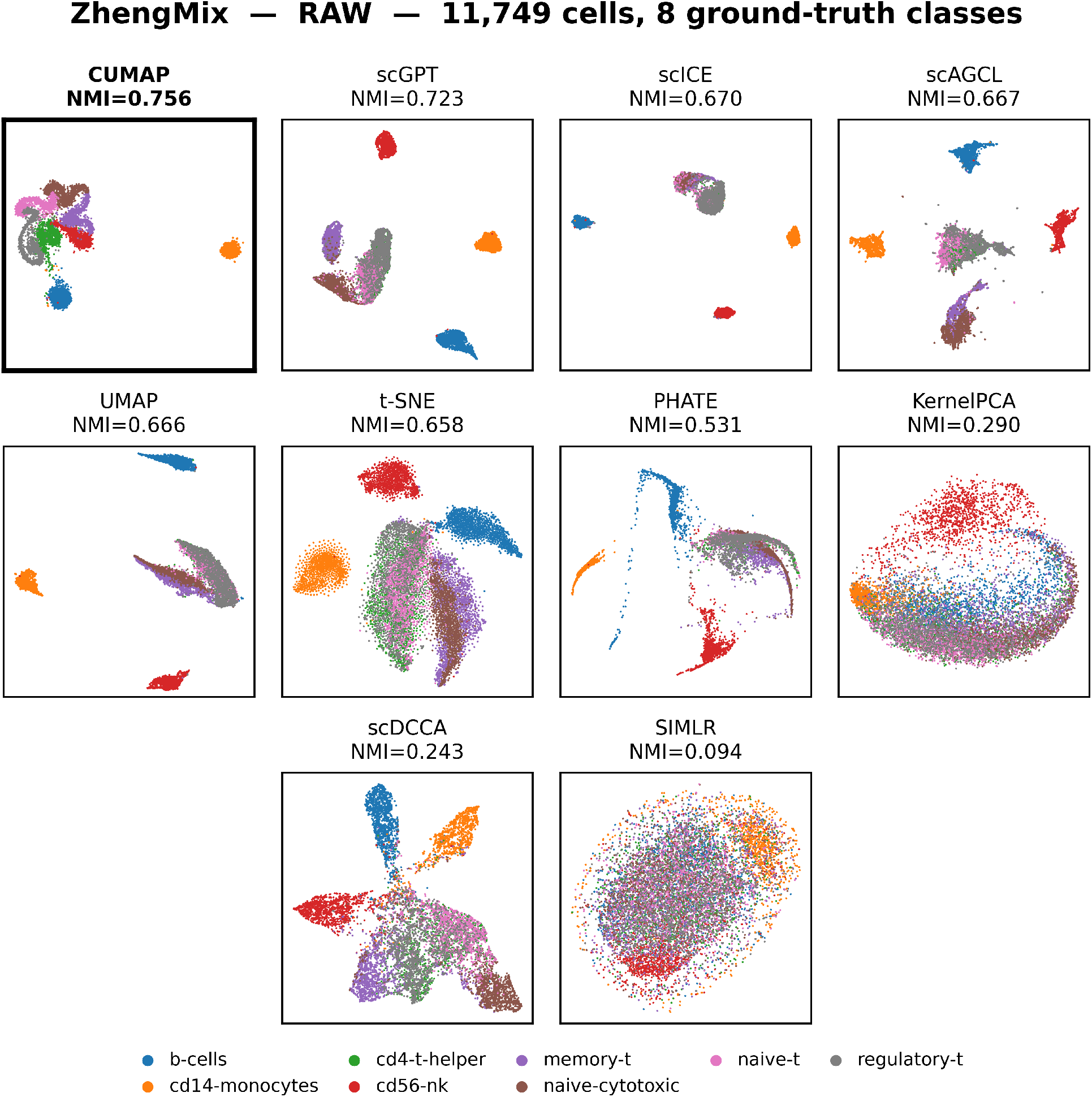
ZhengMix (PBMC), Raw protocol. (11,749 cells, 8 immune cell types). Labels come from FACS sorting; see the text.

**Figure 11:**
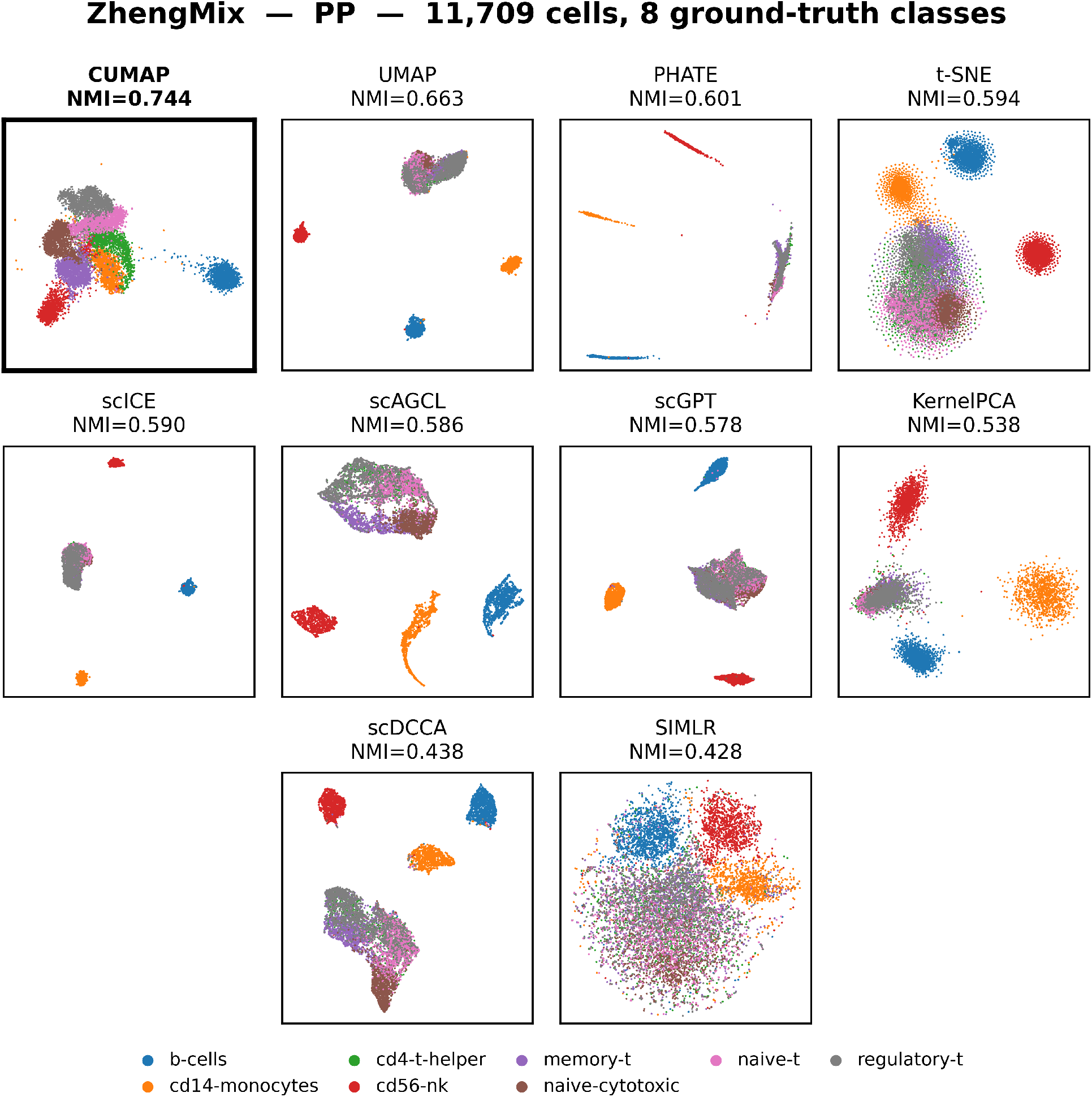
ZhengMix (PBMC), PP protocol. (11,749 cells, 8 immune cell types). Each baseline runs with its recommended preprocessing; c-UMAP reads raw counts throughout.

**Figure 12:**
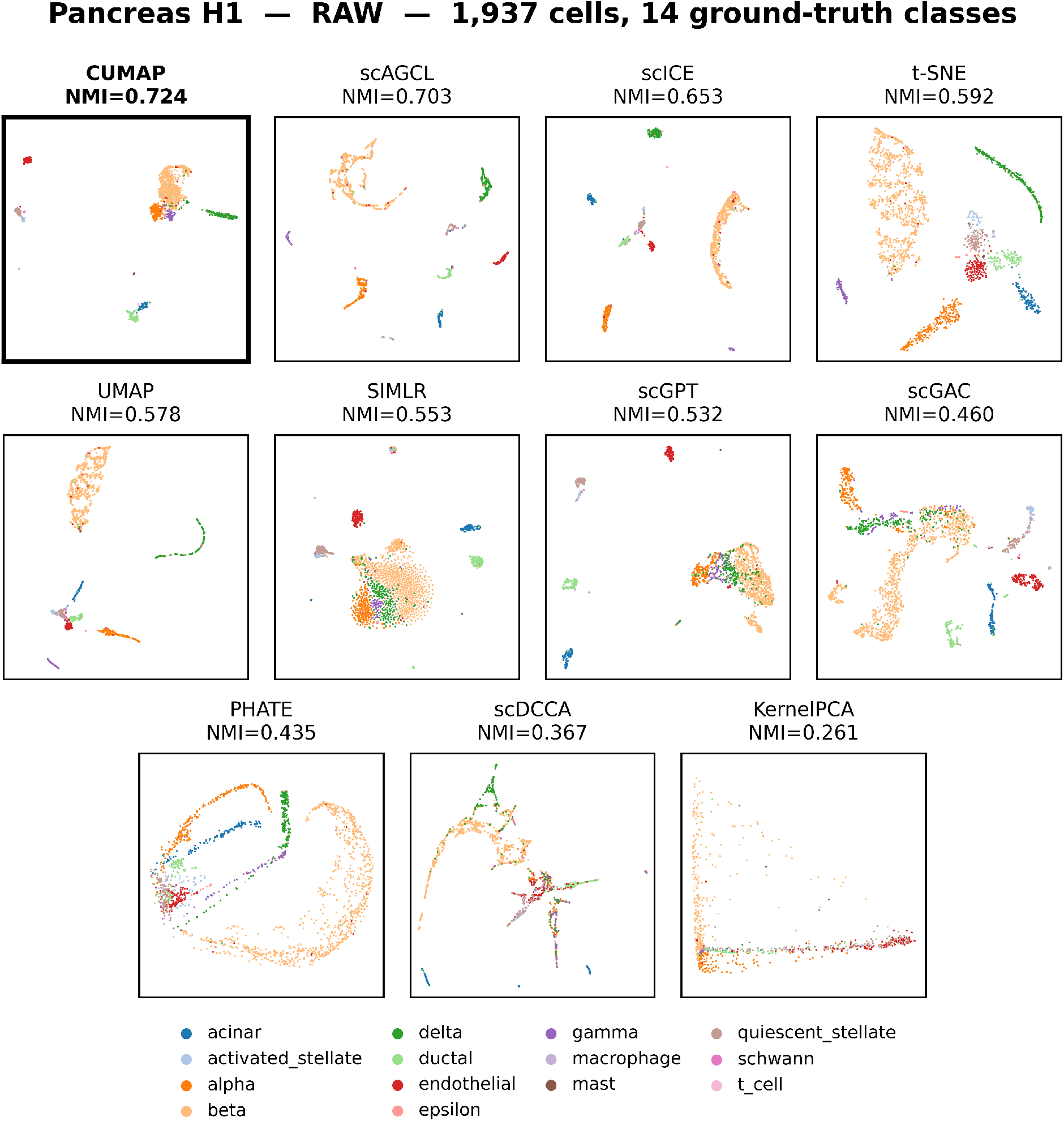
GSE84133 H1 (human pancreas), Raw protocol. (1,937 cells, 14 annotated cell types). c-UMAP places the endocrine (*α, β, δ, γ*) and exocrine (acinar, ductal) populations in distinct groups, with rare immune and supporting types at the periphery.

Together these results show that the cell-driven fusion principle carries from c-TSNE at small to medium scale to c-UMAP at atlas scale. A single fixed weight setting per protocol suffices for all 29 datasets. Under that setting c-UMAP attains the highest mean NMI and AMI in every evaluated case, and it leads all 11 baselines on both indices after Bonferroni correction. It also scales to 1.14 million cells.

## 5. Discussion

The central result of this work is that one fixed fusion rule, reading raw counts, attains the highest mean NMI and AMI across 29 datasets under both protocols while scaling to 1.14 million cells. That sentence bundles several distinct claims, and each deserves separate treatment. We take them in turn below.

### The rule transfers without tuning

A fusion rule earns its place only if it works on data it was not adjusted for. We therefore fixed the weights once per protocol and applied them unchanged to datasets differing 5,000-fold in cell count, spanning 3 to 180 classes, and drawn from three protocol families. Per-dataset tuning would have been the easier path, and also the less informative one, because the tuning signal is exactly what a discovery setting does not provide. That one setting suffices indicates the sparsity-adaptive weighting captures a stable property of scRNA-seq geometry rather than an artifact of any one dataset. The single documented exception reinforces the point, since the entry where the presence/absence view degenerates is identified by a label-free statistic computed from the data itself.

### Preprocessing carries a measurable cost

The case for reading raw counts is usually made on principle. Section 4.12 makes it quantitative instead. Variance-based selection of 2,000 genes removes a median 46.9% of each class’s top-ranked differentially expressed genes. Among them are the markers that define the populations being sought: *IAPP* for beta cells, *FOXP3* for regulatory T cells, *MS4A1* for B cells. A cluster formed after such a step can no longer be annotated by the gene that would have named it. Working from raw counts keeps the full transcriptome available to the fusion. The entry-usage diagnostic can then attribute a boundary to a biological axis computed over every gene, not over a variance-selected subset.

### Scalability belongs to the method, not to the engineering

Published asymptotics describe the layer an author chooses to analyze rather than the pipeline a user runs. Measuring end to end changes the picture: Table 13 shows graph construction accounting for 92% of the strongest baseline’s total time on ZhengMix. Because the lean path of c-UMAP builds no *N × N* matrix at any point, the same fusion and the same weights carry from 225 cells to 1.14 million, with a measured within-dataset exponent of *n*^1.01^. Scale here is a property of the formulation rather than of the implementation.

### Explainability and accuracy move together

c-TSNE and c-UMAP challenge the assumed trade-off between interpretability and performance [7]. Matching the dissimilarity to the data improves clustering and robustness at the same time, without the multi-kernel learning or feature selection that make competing pipelines harder to audit [4]. The entry-usage diagnostic is what turns the fusion itself into an explanation, reporting for each dataset which biological axis the embedding relied on and tying that axis to sparsity (Table 7).

### Implications for cancer research

On Patel, c-TSNE separates glioblastoma subtypes, which connects the method to a concrete clinical problem. The same fusion should extend to other heterogeneous cancers, for instance in isolating leukemic stem cells or resolving triple-negative breast-cancer subtypes. Precise identification of therapeutically relevant populations is the practical benefit, and it supports a broader role in precision medicine.

### Beyond single-cell analysis

The principle generalizes past the application that motivated it. Wherever high-dimensional measurements are sparse and their zeros are informative, similarity can be decomposed into named views, fused by an operator tuned to the sparsity, and audited afterward by asking which view supplied each entry. Mass cytometry, spatial transcriptomics, microbiome count tables, and electronic health records share that structure. For the information-fusion community specifically, the entry-usage diagnostic offers a general template: a fusion operator can be made accountable by reporting its per-entry provenance, at negligible cost and without weakening the fusion.

### Complexity and speedup

The naive *O*(*n*^3^) cost of c-TSNE comes from the full eigen-decomposition used for normalization and from the dense SVD of the L-Chebyshev view, not from t-SNE itself. Section 3.9 removes both cubic terms by building the fused distance only on an approximate *k*-NN graph, with fidelity bounded by Theorem 3. The result is Fast c-TSNE at *O*(*n* log *n p*) time and *O*(*nk* + nnz(*X*)) memory (Table 2), which is near-linear in *n* and matches the c-UMAP asymptotics while retaining the Student-*t* embedding.

## 6. Conclusion

We presented cell-driven information fusion, an explainable framework for single-cell embedding. It builds cell similarity from three named biological views read directly off raw counts and combines them with a sparsity-adaptive power-mean operator. Two instantiations cover the practical range of the problem. c-TSNE leads every Euclidean-geometry baseline on six established benchmarks and is significantly more robust than t-SNE to dropout and noise. c-UMAP carries the same fusion to atlas scale.

Our strongest result concerns a single fixed weight vector per protocol, reading raw counts. It attains the highest mean NMI and AMI across 29 datasets under both protocols and leads all 11 baselines on both indices after Bonferroni correction. The same setting scales to 1.14 million cells. This matters for single-cell biology because it removes two costs analysts now accept as unavoidable. One is tuning an embedding per dataset. The other is discarding marker genes by variance-based selection before clustering begins. It matters beyond single-cell biology because the same construction applies wherever sparse high-dimensional measurements carry informative zeros, from spatial transcriptomics and mass cytometry to microbiome and clinical count data. For information fusion as a field, the entry-usage diagnostic shows that an operator can report the provenance of every fused entry at negligible cost, making accountability a design feature rather than an afterthought.

Taken together, these results indicate that explainability need not be traded against performance, nor against the ability to run at atlas scale. Matching the dissimilarity to the data improves clustering and interpretability at the same time [7, 20]. We recommend sum-fusion as the default at moderate sparsity, with the presence/absence view given dominant weight for highly sparse data. Several directions follow naturally. A principled nonlinear weighting scheme would extend the operator, and adaptive fusion could be driven by statistics beyond sparsity such as data entropy [22]. On the application side, the framework invites use in single-cell classification [21] and pairing with density-based clustering [23, 30].

## Data and code availability

The cumap package, the per-dataset scores behind every table, and the analysis scripts are at https://github.com/hank08819/cumap under the MIT License (`pip install cumap`, Python 3.9). The scores cover c-UMAP and all eleven baselines over the 62 evaluation entries, with each dataset’s cell count, gene count, sparsity, class count and fixed weights, so Tables 10 and 11 can be rebuilt from the release alone; audit.py does so on a CPU in under a minute. All datasets are public, drawn from GSE84133 [36], GSE116470 [37], SRP073767 [38] and further sources; accessions and label sets are in Table C.20.

## CRediT authorship contribution statement

**Henry Han:** Conceptualization, Methodology, Supervision, Software, Writing – original draft, Writing – review and editing. **Tianyu Zhang:** Methodology, Validation, Data curation, Software. **Luyu Niu:** Software, Data curation, Visualization, Validation. **Jiacun Wang:** Investigation, Validation. **Fei Han:** Investigation, Validation. **Chun Li:** Investigation, Validation. **Junyi Li:** Investigation, Validation. **Huaixin Cao:** Investigation, Validation.

## Declaration of generative AI in the writing process

During the preparation of this work the authors used a generative AI assistant solely to polish the English and grammar.

## Declaration of competing interest

The authors declare that they have no competing interests.

## Appendix A. Supporting analyses

### Appendix A.1. SNF comparison: cell-driven fusion vs. cross-diffusion fusion

Information-fusion reviewers expect a comparison against an established *fusion* operator, not only against embedding/clustering baselines. We therefore benchmark against Similarity Network Fusion (SNF) [31] under a protocol that isolates the fusion operator. SNF and our cell-driven fusion are given the identical three cell-driven views, so any difference is attributable to the fusion rule alone.

#### Protocol

For each dataset we (1) compute the three cell-driven dissimilarities, each median-normalized; (2) for SNF, convert each to an affinity with the scaled-exponential kernel on *K*-NN graphs (*K* = 20, *µ* = 0.5, *t* = 20 cross-diffusion iterations); (3) embed both fused distances by t-SNE and cluster by K-means at the ground-truth class number.

#### Result

Under this fully matched pipeline (Table A.15), cell-driven fusion and SNF are *statistically comparable overall* (paired Wilcoxon *p* = 0.12, *n* = 7). The advantage is concentrated on the structurally harder, higher-sparsity datasets, where cell-driven fusion wins on all four: Zeisel (+0.13), Patel (+0.09), Goolam (+0.14), Usoskin (+0.03).

**Table A.15:** SNF fusion baseline under a fully matched pipeline. Both fusions use the same three cell-driven views and the same downstream (t-SNE+K-means). This comparison median-normalizes the views so that both operators receive identical inputs, so its absolute values sit on a different scale from the eigenvalue-normalized protocol of Table 6. What it measures is the difference between the two columns. Best of the matched columns in **bold**. Rows ordered by sparsity.

| Dataset | Sparsity (%) | Ours (t-SNE+KM) | SNF (t-SNE+KM) | SNF (spec.) |
| --- | --- | --- | --- | --- |
| Kolod | 27.9 | **1.000** | **1.000** | 0.972 |
| Buettner | 37.9 | 0.732 | **0.831** | 0.839 |
| Zeisel | 46.0 | **0.637** | 0.509 | 0.628 |
| Pollen | 51.0 | 0.912 | **0.932** | 0.956 |
| Patel | 53.3 | **0.845** | 0.757 | 0.621 |
| Goolam | 68.6 | **0.654** | 0.514 | 0.527 |
| Usoskin | 78.1 | **0.711** | 0.683 | 0.492 |
| Mean |  | **0.784** | 0.747 | 0.719 |

### Appendix A.2. Per-distance ablation

To validate the contribution of each cell-driven view, and to motivate the fixed weights adopted in Section 4.9, we evaluate the framework under each subset of the three metrics on GSE84133 H1 (Table A.16). The ranking is the informative quantity here: the combination that maximizes NMI is (*w*_*y*_, *w*_*f*_, *w*_*c*_) = (0.9, 0, 0.1), which is exactly the fixed Raw weight vector applied unchanged to all 29 datasets in the atlas-scale benchmark. The ablation therefore selects the same operator that the benchmark then validates without further tuning.

**Table A.16:** Per-distance ablation on GSE84133 H1, run under the exact dense-affinity pipeline used to choose the operator. Y = Yule, F = fractional, C = L-Chebyshev. Best in **bold**. Absolute values follow this selection pipeline and are not on the same scale as the fixed-weight benchmark of Table 11; the ordering of the rows is what motivates the weight choice.

| Distance combination | $(w_y, w_f, w_c)$ | Routes | NMI |
| --- | --- | --- | --- |
| Yule only | (1, 0, 0) | 1 | 0.7321 |
| Fractional only | (0, 1, 0) | 1 | 0.4187 |
| L-Chebyshev only | (0, 0, 1) | 1 | 0.7104 |
| Y+F | (0.5, 0.5, 0) | 2 | 0.7193 |
| F+C | (0, 0.5, 0.5) | 2 | 0.6921 |
| Y+F+C (full) | (0.7, 0.1, 0.2) | 3 | 0.7843 |
| **Y+C (used)** | **(0.9, 0, 0.1)** | **2** | **0.7942** |

### Appendix A.3. Cortex Leiden-CPM ablation

To support the choice of K-means for the imbalanced cortex atlases, we also run a Leiden CPM resolution scan on the c-UMAP embedding (Table A.17).

**Table A.17:** Cortex Leiden-CPM ablation. c-UMAP wins under either downstream clustering.

| Dataset | c-UMAP + K-means | c-UMAP + Leiden | scICE |
| --- | --- | --- | --- |
| Cortex Posterior | **0.4919** | 0.4421 | 0.3818 |
| Cortex Frontal | **0.5485** | 0.4807 | 0.4529 |

### Appendix A.4. Fusion-weight grid and the eigengap rule

The eigengap rule of Section 3.13 selects (*α, w*) label-free on a per-dataset basis and is the general form of the operator. For the atlas-scale benchmark we deliberately give up that freedom and fix the weights once per protocol (Section 4.9): [0.90, 0.00, 0.10] under Raw and [0.80, 0.10, 0.10] under PP, applied unchanged to all 29 datasets. Both settings place the dominant weight on the presence/absence view, consistent with the 87–98% sparsity of these data and with the entry-usage analysis of Section 4.5. Reporting the fixed setting is the stricter test: it shows the advantage in Table 11 is a property of the fusion rather than of per-dataset selection.

### Appendix A.5. Fast c-TSNE benchmark (near-linear scaling)

We implemented Fast c-TSNE approximate *k*-NN under the cell-driven views via NN-descent (pynndescent) feeding a Barnes–Hut t-SNE and the exact variant. A least-squares fit gives *n*^2.00^ for the exact variant and *n*^0.92^ **for Fast c-TSNE**, which is empirically near-linear and confirms the *O*(*n* log *np*) analysis. Table A.19 reports the measured wall-clock times, where the speedup reaches 13.7 **at** *× n* = 3005.

**Table A.18:** Fidelity of Fast c-TSNE (*k* = 60) against exact c-TSNE, NMI_max_ averaged over 3 seeds. Both columns use the uniform-weight sum operator throughout, rather than the sparsity routing of Algorithm 1, so that they differ only in the *k*-NN truncation. Absolute values are therefore on a different scale from Table 6. The truncation is lossless, with Fast c-TSNE matching or exceeding exact c-TSNE on every dataset.

| Dataset | $n$ | exact | Fast c-TSNE | gap |
| --- | --- | --- | --- | --- |
| Goolam | 124 | 0.664 | 0.664 | +0.00 |
| Buettner | 182 | 0.547 | 0.572 | +0.03 |
| Pollen | 249 | 0.925 | 0.925 | +0.00 |
| Patel | 543 | 0.739 | 0.749 | +0.01 |
| Usoskin | 622 | 0.671 | 0.785 | +0.11 |
| Kolod | 704 | 1.000 | 1.000 | +0.00 |
| Mean |  | 0.758 | **0.782** | +0.02 |

## Appendix B. Representative embeddings

## Appendix C. Per-dataset tables

### Appendix C.1. Per-dataset catalogue of the 29 benchmark datasets

**Table C.20:** The 29 datasets used in this study.

| Dataset | Species | Tissue | Source | Cells | Genes | $K$ | Sparsity (%) |
| --- | --- | --- | --- | --- | --- | --- | --- |
| TS full atlas | Human | Whole body | TS v2 | 1,136,218 | 61,497 | 180 | 94.8 |
| TS immune | Human | Immune | TS v2 | 508,731 | 60,606 | 42 | 95.5 |
| Cortex frontal | Mouse | Brain cortex | GSE116470 | 71,639 | 29,463 | 14 | 93.2 |
| Cortex posterior | Mouse | Brain cortex | GSE116470 | 46,346 | 27,720 | 14 | 93.5 |

Table C.20 continued from previous page
| Dataset | Species | Tissue | Source | Cells | Genes | $K$ | Sparsity (%) |
| --- | --- | --- | --- | --- | --- | --- | --- |
| ZhengMix | Human | PBMC | SRP073767 | 11,749 | 11,684 | 8 | 95.3 |
| Lung atlas | Human | Lung | GSE130148 | 10,360 | 16,327 | 13 | 96.6 |
| HCL trachea | Human | Trachea | GSE134355 | 9,949 | 24,990 | 14 | 97.1 |
| HCL kidney | Human | Kidney | GSE134355 | 9,153 | 25,601 | 18 | 97.5 |
| HCL adrenal | Human | Adrenal gland | GSE134355 | 8,114 | 23,842 | 13 | 97.7 |
| HCL muscle | Human | Skeletal muscle | GSE134355 | 7,775 | 22,961 | 17 | 97.9 |
| HCL blood | Human | Peripheral blood | GSE134355 | 7,160 | 21,267 | 13 | 98.0 |
| HCL duodenum | Human | Duodenum | GSE134355 | 5,200 | 30,172 | 10 | 91.2 |
| MCA fetal stomach | Mouse | Fetal stomach | GSE108097 | 6,192 | 19,720 | 16 | 97.4 |
| MCA fetal intestine | Mouse | Fetal intestine | GSE108097 | 6,076 | 20,115 | 13 | 96.7 |
| Tian sc 10x 5-line | Human | Lung cell lines | GSE118767 | 3,918 | 11,786 | 5 | 63.0 |
| Tian sc 10x | Human | Lung cell lines | GSE118767 | 902 | 16,468 | 3 | 45.0 |
| Tian sc Drop-seq | Human | Lung cell lines | GSE118767 | 225 | 15,127 | 3 | 62.1 |
| Pancreas H1 | Human | Pancreatic islet | GSE84133 | 1,937 | 20,125 | 14 | 90.4 |
| Pancreas H2 | Human | Pancreatic islet | GSE84133 | 1,724 | 20,125 | 14 | 90.6 |
| Pancreas H3 | Human | Pancreatic islet | GSE84133 | 3,605 | 20,125 | 14 | 91.3 |
| Pancreas H4 | Human | Pancreatic islet | GSE84133 | 1,303 | 20,125 | 14 | 89.0 |
| Pancreas M1 | Mouse | Pancreatic islet | GSE84133 | 822 | 14,878 | 13 | 90.5 |
| Pancreas M2 | Mouse | Pancreatic islet | GSE84133 | 1,064 | 14,878 | 13 | 87.8 |
| Wu sNuc-DropSeq | Mouse | Kidney | GSE119531 | 3,011 | 19,713 | 13 | 95.4 |
| Wu DroNc-seq | Mouse | Kidney | GSE119531 | 2,769 | 19,713 | 13 | 94.3 |
| Enge | Human | Pancreatic islet | GSE81547 | 2,282 | 23,460 | 6 | 86.0 |
| Chu | Human | hESC diff. | GSE75748 | 1,018 | 19,097 | 7 | 49.8 |
| Koh | Human | hESC/mesoderm | Duò 2018 | 531 | 48,981 | 9 | 72.1 |
| Kumar | Mouse | mESC | Duò 2018 | 246 | 45,159 | 3 | 45.8 |

**Figure B.13:**
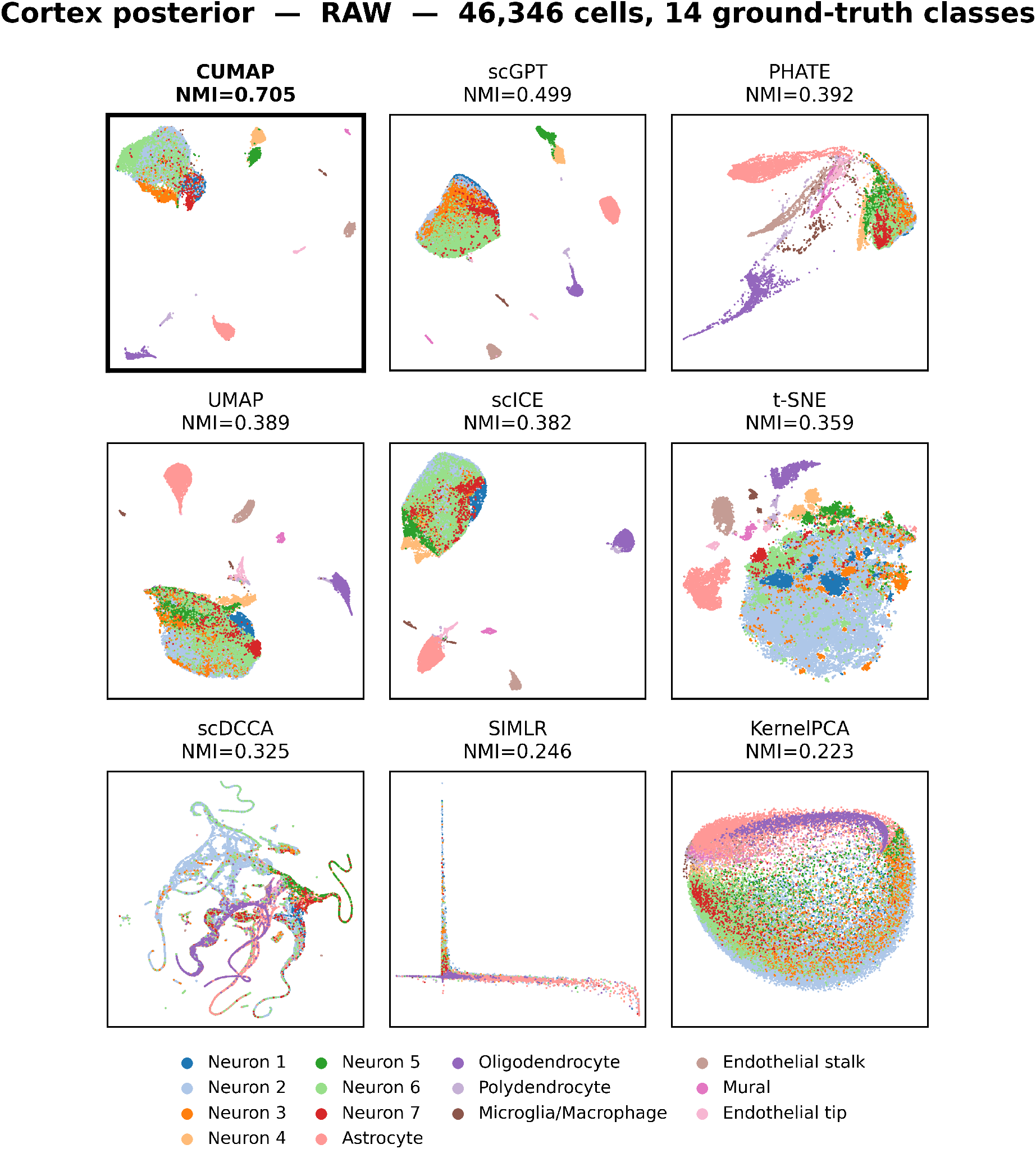
DropViz Cortex Posterior, Raw protocol. (46,346 cells, 14 ground-truth classes). Each panel is one method in its own embedding, colored by ground-truth cell type.

**Figure B.14:**
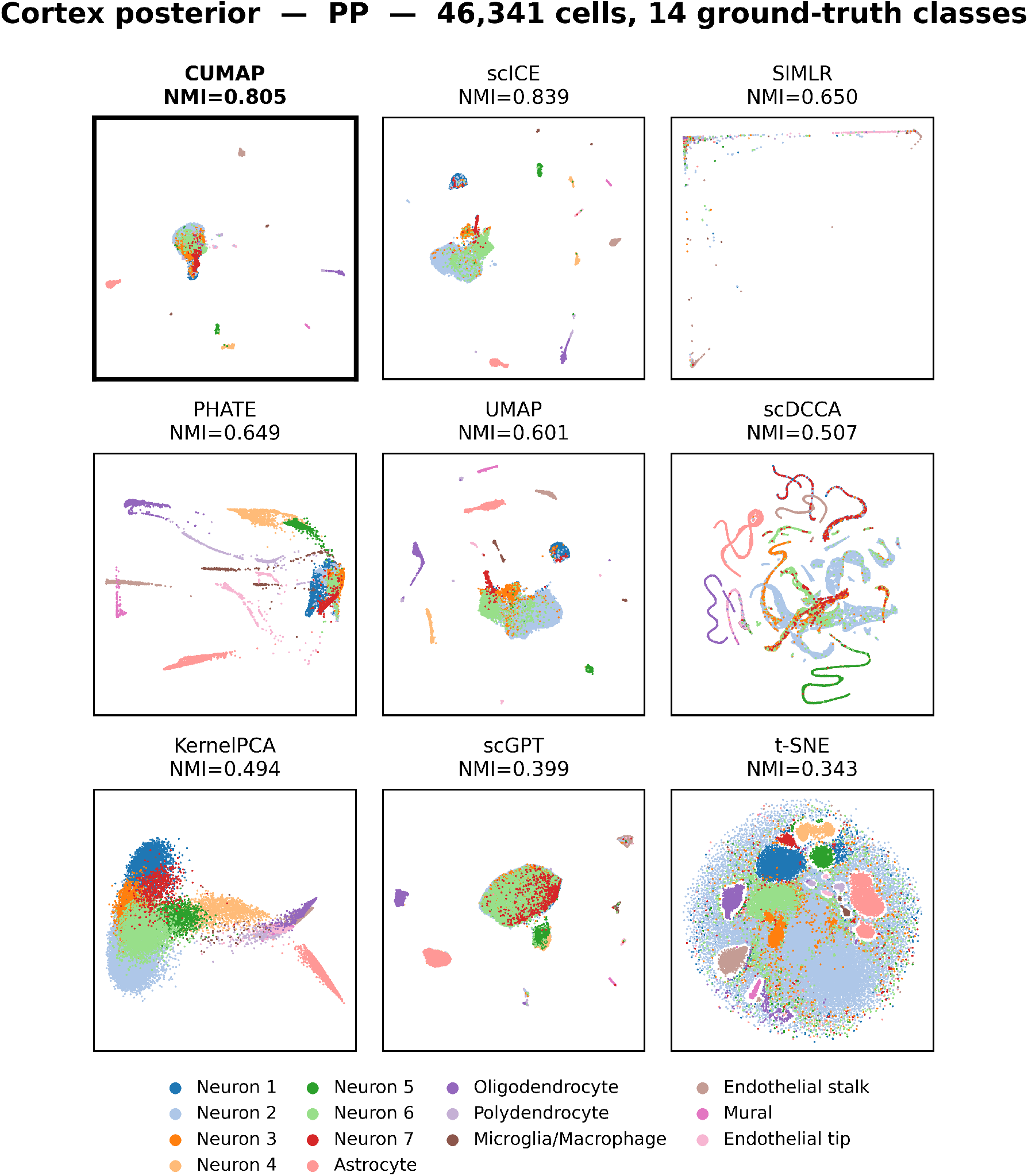
DropViz Cortex Posterior, PP protocol. (46,341 cells, 14 ground-truth classes). Each panel is one method in its own embedding, colored by ground-truth cell type.

**Figure B.15:**
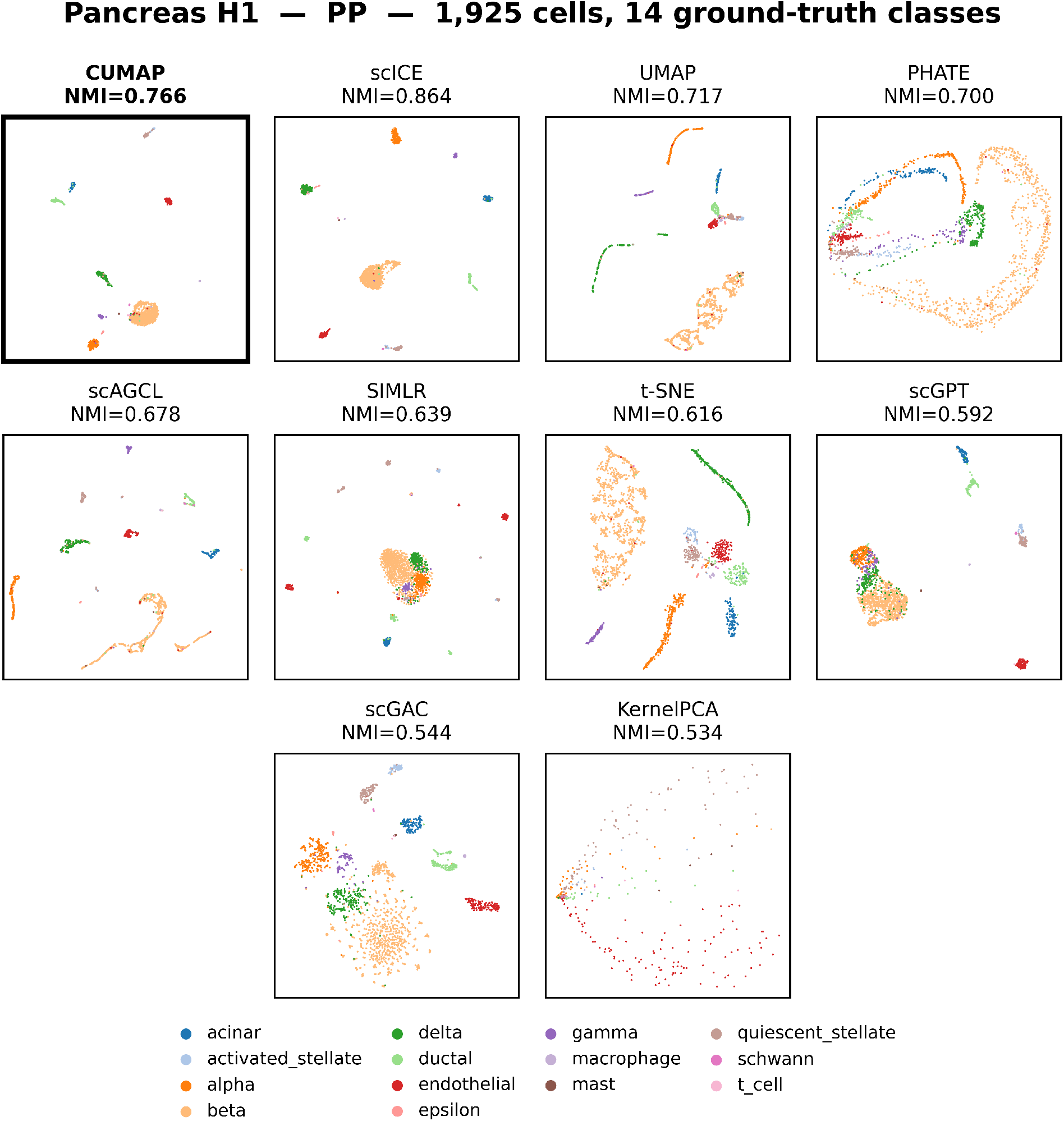
GSE84133 H1 (human pancreas), PP protocol. (1,925 cells, 14 ground-truth classes). Each panel is one method in its own embedding, colored by ground-truth cell type.

**Table A.19:** Measured wall-clock (s) of Fast c-TSNE vs. exact c-TSNE on subsampled Zeisel (*D* = 4412 genes), single CPU node. Fast c-TSNE scales as *n*^0.92^ vs. *n*^2.00^ for exact, a 13.7*×* speedup at *n* = 3005.

| $n$ | exact (s) | Fast (s) | speedup |
| --- | --- | --- | --- |
| 300 | 4.8 | 4.6 | $1.0\times$ |
| 600 | 16.2 | 7.4 | $2.2\times$ |
| 1200 | 57.1 | 14.3 | $4.0\times$ |
| 2000 | 183.1 | 24.3 | $7.5\times$ |
| 3005 | 505.3 | 36.9 | **$13.7\times$** |
| fit | $n^{2.00}$ | **$n^{0.92}$** | |

### Appendix C.2. Per-dataset NMI, Raw protocol

**Table C.21:** NMI on the RAW condition (no preprocessing), fixed weights [0.9, 0.0, 0.1]. Best per row in bold; a dash (–) marks an entry the method could not produce, with a superscript giving the reason (legend in the text). *CUMAP wins* and *Wilcoxon p* are computed on RAW and PP combined, so these two rows are identical in the RAW and PP tables of the same metric; the denominator is the number of paired entries the baseline actually produced.

| Dataset | Cells | *K* | CUMAP | scICE | tSNE | UMAP | KPCA | PHATE | SIMLR | scDCCA | scGAC | scAGCL | sGPT<sub>K</sub> | sGPT<sub>L</sub> |
| --- | --- | --- | --- | --- | --- | --- | --- | --- | --- | --- | --- | --- | --- | --- |
| Tian_sc_dropseq | 225 | 3 | **0.8632** | 0.7554 | 0.3800 | 0.3563 | 0.0346 | 0.3502 | 0.7222 | 0.7766 | 0.3942 | 0.7316 | 0.3081 | 0.2621 |
| kumar | 246 | 3 | 0.9262 | 0.9586 | 0.9235 | 0.9385 | 0.0258 | 0.9385 | 0.9647 | 0.8968 | **1.0000** | 0.8951 | 0.3329 | 0.2753 |
| koh | 531 | 9 | **0.7489** | 0.6842 | 0.4671 | 0.4453 | 0.0565 | 0.4431 | 0.5550 | 0.1343 | 0.5226 | 0.6834 | 0.3768 | 0.3472 |
| M1 | 822 | 13 | 0.6091 | **0.6984** | 0.4943 | 0.4962 | 0.2256 | 0.4304 | 0.4980 | 0.2971 | 0.4395 | 0.5481 | 0.5114 | 0.5918 |
| Tian_sc_10x | 902 | 3 | **0.9809** | 0.9743 | 0.9688 | 0.9743 | 0.0045 | 0.9743 | 0.9729 | 0.5624 | **0.9809** | 0.9743 | 0.5077 | 0.3736 |
| Chu | 1,018 | 7 | **0.8701** | 0.7795 | 0.6054 | 0.6953 | 0.0626 | 0.5198 | 0.5234 | 0.7339 | 0.6293 | 0.8214 | 0.6130 | 0.6073 |
| M2 | 1,064 | 13 | 0.6138 | 0.5899 | 0.4656 | 0.4846 | 0.1990 | 0.4070 | 0.3795 | 0.1665 | 0.4417 | **0.6188** | 0.3942 | 0.3999 |
| H4 | 1,303 | 14 | **0.7797** | 0.5938 | 0.5087 | 0.5177 | 0.2503 | 0.4546 | 0.5103 | 0.3699 | 0.3631 | 0.6022 | 0.4409 | 0.5004 |
| H2 | 1,724 | 14 | **0.7338** | 0.6369 | 0.5539 | 0.5698 | 0.2822 | 0.4544 | 0.5161 | 0.3829 | 0.5015 | 0.6586 | 0.4906 | 0.5206 |
| H1 | 1,937 | 14 | **0.7236** | 0.6530 | 0.5918 | 0.5775 | 0.2608 | 0.4350 | 0.5529 | 0.3671 | 0.4603 | 0.7030 | 0.5395 | 0.5317 |
| Enge | 2,282 | 6 | 0.6655 | **0.7526** | 0.5704 | 0.5962 | 0.0089 | 0.5963 | 0.3207 | 0.4337 | 0.3628 | 0.5342 | 0.4025 | 0.3838 |
| wu_kidney_droncseq | 2,769 | 13 | **0.4148** | 0.3509 | 0.1208 | 0.1192 | 0.0577 | 0.0952 | 0.3562 | 0.1545 | 0.2329 | 0.2938 | 0.4058 | 0.4039 |
| wu_kidney_snucdropseq | 3,011 | 13 | 0.3839 | 0.2541 | 0.1324 | 0.1078 | 0.0355 | 0.0441 | 0.1324 | 0.1266 | 0.0710 | 0.2378 | 0.3942 | **0.4080** |
| H3 | 3,605 | 14 | **0.8078** | 0.6387 | 0.5526 | 0.5829 | 0.3548 | 0.5271 | 0.4816 | 0.4517 | 0.4814 | 0.7118 | 0.5289 | 0.6126 |
| Tian_sc_10x_5cl | 3,918 | 5 | **0.9370** | 0.9365 | 0.7686 | 0.8713 | 0.0214 | 0.6668 | 0.9261 | 0.7108 | 0.6249 | 0.9318 | 0.3231 | 0.4305 |
| duodenum | 5,200 | 10 | 0.3279 | 0.4714 | 0.3592 | 0.3758 | 0.3156 | 0.3471 | 0.5636 | 0.5151 | 0.5172 | 0.5609 | **0.5800** | 0.5179 |
| mca_fetalintestine | 6,076 | 13 | 0.3435 | 0.1882 | 0.2472 | 0.2746 | 0.1245 | 0.2154 | 0.0502 | 0.1380 | 0.1635 | 0.2510 | 0.4252 | **0.5147** |
| mca_fetalstomach | 6,192 | 16 | 0.1914 | 0.0910 | 0.2407 | 0.2168 | 0.1101 | 0.1257 | 0.0266 | 0.1582 | 0.0005 | 0.3375 | 0.3473 | **0.4573** |
| hcl_peripheralblood4 | 7,160 | 13 | 0.4693 | 0.4142 | 0.4886 | 0.4959 | 0.3611 | 0.4489 | 0.0328 | 0.1749 | 0.1265 | 0.5022 | 0.4711 | **0.5601** |
| hcl_muscle1 | 7,775 | 17 | 0.2917 | 0.3261 | 0.3180 | 0.3285 | 0.1508 | 0.2623 | 0.0138 | 0.1500 | 0.0641 | 0.2997 | 0.3430 | **0.4773** |
| hcl_adrenalgland2 | 8,114 | 13 | 0.3674 | <sup>g</sup> | 0.3646 | 0.3756 | 0.2061 | 0.2848 | <sup>h</sup> | 0.2886 | 0.1254 | 0.4772 | 0.3958 | **0.5063** |
| hcl_kidney2 | 9,153 | 18 | 0.4334 | 0.1249 | 0.2998 | 0.2949 | 0.1228 | 0.1352 | 0.0501 | 0.3064 | 0.0618 | 0.4402 | 0.4354 | **0.5448** |
| hcl_trachea2 | 9,949 | 14 | **0.6171** | 0.0376 | 0.4554 | 0.4655 | 0.1596 | 0.3472 | 0.0748 | 0.3797 | 0.1375 | 0.5141 | 0.4524 | 0.5599 |
| Lung | 10,360 | 13 | 0.4419 | 0.1141 | 0.3919 | 0.3712 | 0.1407 | 0.1823 | 0.0659 | 0.1624 | <sup>c</sup> | 0.4846 | 0.4536 | **0.5742** |
| ZhengMix | 11,749 | 8 | **0.7560** | 0.6702 | 0.6583 | 0.6663 | 0.2900 | 0.5307 | 0.0943 | 0.2431 | <sup>c</sup> | 0.6671 | 0.7004 | 0.7234 |
| Cortex_P | 46,346 | 14 | **0.7051** | 0.3818 | 0.3591 | 0.3891 | 0.2227 | 0.3922 | 0.2456 | 0.3253 | <sup>c</sup> | <sup>f</sup> | 0.4436 | 0.4995 |
| Cortex_F | 71,639 | 14 | **0.8153** | 0.4529 | 0.4452 | 0.4813 | 0.2584 | 0.4129 | 0.2226 | 0.2657 | <sup>c</sup> | <sup>f</sup> | 0.4919 | 0.5163 |
| Cortex_P (K=93) | 46,346 | 93 | **0.5421** | 0.1839 | 0.3603 | 0.3123 | 0.1895 | 0.2923 | 0.1595 | 0.3765 | <sup>c</sup> | <sup>f</sup> | 0.3801 | 0.3961 |
| Cortex_F (K=81) | 71,639 | 81 | **0.5362** | 0.2662 | 0.3680 | 0.3391 | 0.2067 | 0.3493 | 0.2048 | 0.4830 | <sup>c</sup> | <sup>f</sup> | 0.3828 | 0.4158 |
| TS50W_raw | 508,731 | 42 | **0.5904** | <sup>b</sup> | <sup>e</sup> | <sup>e</sup> | <sup>c</sup> | <sup>e</sup> | <sup>c</sup> | <sup>d</sup> | <sup>c</sup> | <sup>c</sup> | 0.4991 | 0.5403 |
| TS100W_raw | 1,136,218 | 180 | 0.6770 | <sup>b</sup> | <sup>e</sup> | <sup>e</sup> | <sup>c</sup> | <sup>e</sup> | <sup>c</sup> | <sup>d</sup> | <sup>c</sup> | <sup>c</sup> | 0.5574 | **0.7505** |
| Mean |  |  | 0.6182 | 0.4993 | 0.4641 | 0.4731 | 0.1634 | 0.4022 | 0.3649 | 0.3632 | 0.3784 | 0.5792 | 0.4493 | 0.4904 |
| CUMAP wins |  |  | – | 42/57 | 48/58 | 45/58 | 55/58 | 46/58 | 52/55 | 52/58 | 39/46 | 45/54 | 49/59 | 47/59 |
| Wilcoxon p |  |  | – | 2.5e-5 | 9.0e-9 | 8.7e-8 | 6.7e-11 | 1.3e-7 | 1.2e-9 | 7.1e-9 | 8.3e-8 | 1.5e-6 | 4.7e-9 | 2.6e-6 |

### Appendix C.3. Per-dataset NMI, PP protocol

**Table C.22:**
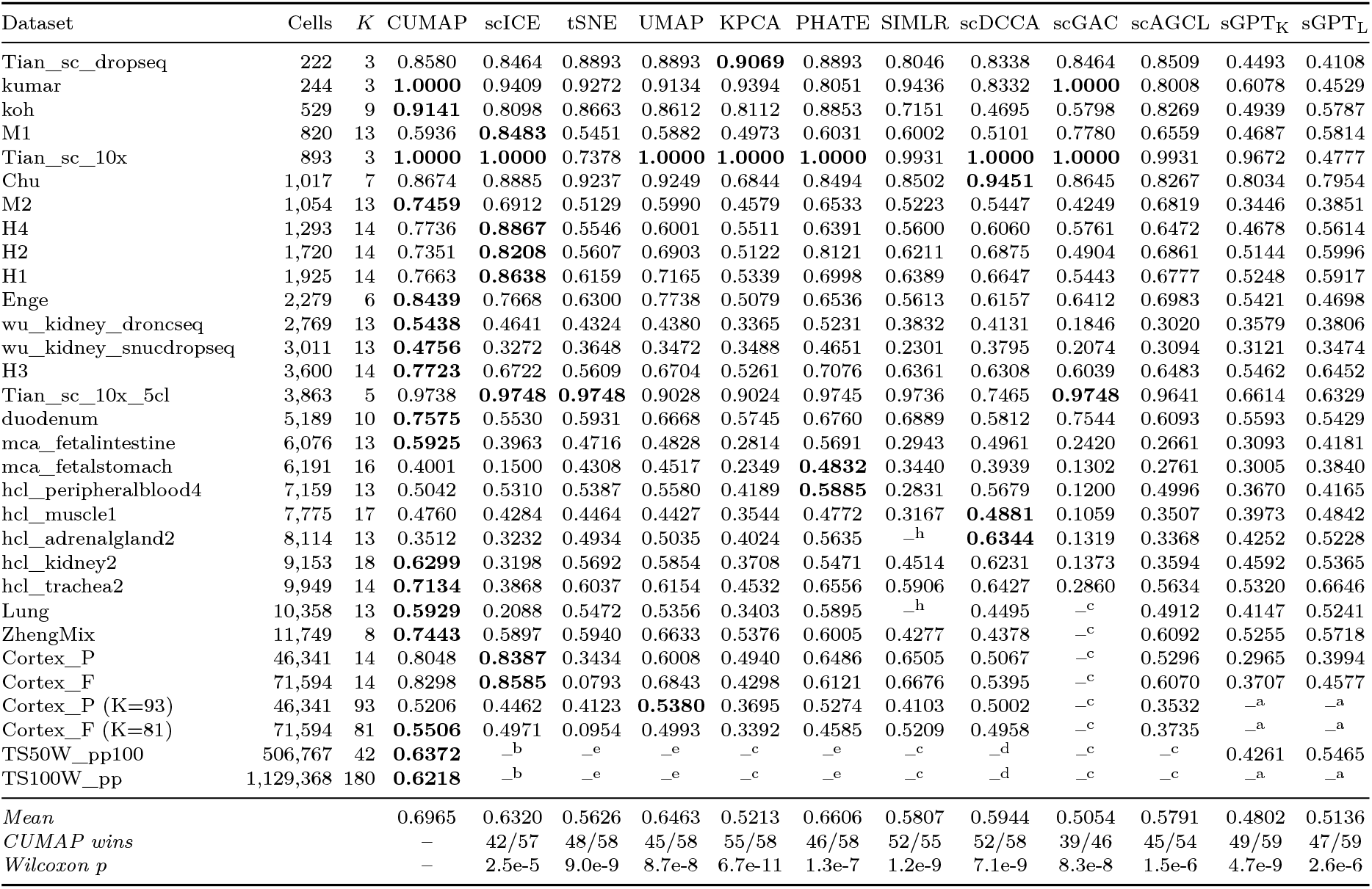
NMI on the PP condition (standard preprocessing), fixed weights [0.8, 0.1, 0.1] (TS full atlas: [0.00, 0.20, 0.80], see Section 4.9). Best per row in bold; a dash (–) marks an entry the method could not produce, with a superscript giving the reason (legend in the text). *CUMAP wins* and *Wilcoxon p* are computed on RAW and PP combined, so these two rows are identical in the RAW and PP tables of the same metric; the denominator is the number of paired entries the baseline actually produced.

### Appendix C.4. Per-dataset AMI, Raw protocol

**Table C.23:** AMI on the RAW condition (no preprocessing), fixed weights [0.9, 0.0, 0.1]. Best per row in bold; a dash (–) marks an entry the method could not produce, with a superscript giving the reason (legend in the text). *CUMAP wins* and *Wilcoxon p* are computed on RAW and PP combined, so these two rows are identical in the RAW and PP tables of the same metric; the denominator is the number of paired entries the baseline actually produced.

| Dataset | Cells | K | CUMAP | scICE | tSNE | UMAP | KPCA | PHATE | SIMLR | scDCCA | scGAC | scAGCL | sGPT<sub>K</sub> | sGPT<sub>L</sub> |
| --- | --- | --- | --- | --- | --- | --- | --- | --- | --- | --- | --- | --- | --- | --- |
| Tian_sc_dropseq | 225 | 3 | **0.8652** | 0.7868 | 0.3751 | 0.3530 | 0.0410 | 0.3464 | 0.7199 | 0.7786 | 0.5244 | 0.7326 | 0.3040 | 0.3213 |
| kumar | 246 | 3 | 0.9272 | 0.9591 | 0.9233 | 0.9386 | 0.0286 | 0.9386 | 0.9644 | 0.8966 | **1.0000** | 0.8947 | 0.3343 | 0.3301 |
| koh | 531 | 9 | **0.7496** | 0.6782 | 0.4514 | 0.4284 | 0.0497 | 0.4273 | 0.5415 | 0.1116 | 0.6078 | 0.7032 | 0.3595 | 0.3559 |
| M1 | 822 | 13 | 0.7077 | **0.7562** | 0.5876 | 0.5842 | 0.2233 | 0.5031 | 0.5667 | 0.3355 | 0.4930 | 0.6463 | 0.5973 | 0.6500 |
| Tian_sc_10x | 902 | 3 | **0.9810** | 0.9743 | 0.9688 | 0.9743 | 0.0038 | 0.9743 | 0.9728 | 0.5704 | **0.9810** | 0.9743 | 0.5089 | 0.4714 |
| Chu | 1,018 | 7 | **0.8896** | 0.7958 | 0.6033 | 0.6940 | 0.0790 | 0.5180 | 0.5180 | 0.7240 | 0.6487 | 0.8615 | 0.6119 | 0.6408 |
| M2 | 1,064 | 13 | 0.6798 | 0.6520 | 0.5562 | 0.5754 | 0.2075 | 0.4797 | 0.4338 | 0.1775 | 0.4839 | **0.7113** | 0.4538 | 0.4634 |
| H4 | 1,303 | 14 | **0.7955** | 0.7032 | 0.6183 | 0.6223 | 0.2308 | 0.5452 | 0.6055 | 0.4384 | 0.4018 | 0.7156 | 0.5132 | 0.5609 |
| H2 | 1,724 | 14 | **0.8050** | 0.7349 | 0.6611 | 0.6752 | 0.2677 | 0.5369 | 0.5858 | 0.4385 | 0.5088 | 0.7516 | 0.5563 | 0.5845 |
| H1 | 1,937 | 14 | 0.7746 | 0.7403 | 0.6846 | 0.6636 | 0.2567 | 0.4968 | 0.6103 | 0.4148 | 0.5063 | **0.7780** | 0.5983 | 0.6112 |
| Enge | 2,282 | 6 | 0.7022 | **0.7517** | 0.6216 | 0.6501 | 0.0065 | 0.6496 | 0.3183 | 0.4708 | 0.3635 | 0.5821 | 0.4359 | 0.5065 |
| wu_kidney_droncseq | 2,769 | 13 | 0.4376 | 0.3796 | 0.1264 | 0.1236 | 0.0543 | 0.0960 | 0.3470 | 0.1709 | 0.2573 | 0.3154 | 0.4376 | **0.4501** |
| wu_kidney_snucdropseq | 3,011 | 13 | 0.3815 | 0.3336 | 0.1389 | 0.1103 | 0.0284 | 0.0385 | 0.1229 | 0.1338 | 0.1170 | 0.2550 | 0.4192 | **0.4252** |
| H3 | 3,605 | 14 | **0.8398** | 0.7329 | 0.6572 | 0.6865 | 0.3611 | 0.6128 | 0.5545 | 0.5288 | 0.5112 | 0.7743 | 0.6220 | 0.6899 |
| Tian_sc_10x_5cl | 3,918 | 5 | **0.9405** | 0.9395 | 0.7753 | 0.8714 | 0.0361 | 0.6730 | 0.9260 | 0.6374 | 0.6207 | 0.9350 | 0.3247 | 0.5101 |
| duodenum | 5,200 | 10 | 0.3804 | 0.6223 | 0.3928 | 0.4051 | 0.3340 | 0.3605 | 0.5616 | 0.5296 | 0.5648 | 0.6297 | **0.6581** | 0.6333 |
| mca_fetalintestine | 6,076 | 13 | 0.4459 | 0.2840 | 0.2654 | 0.2901 | 0.1245 | 0.2223 | 0.0446 | 0.1478 | 0.1800 | 0.2666 | 0.4513 | **0.5413** |
| mca_fetalstomach | 6,192 | 16 | 0.2460 | 0.1495 | 0.2556 | 0.2275 | 0.1044 | 0.1205 | 0.0185 | 0.1559 | 0.0002 | 0.3525 | 0.3685 | **0.4657** |
| hcl_peripheralblood4 | 7,160 | 13 | 0.5494 | 0.5368 | 0.5419 | 0.5455 | 0.3639 | 0.4564 | 0.0276 | 0.1927 | 0.1706 | 0.5463 | 0.5154 | **0.6055** |
| hcl_muscle1 | 7,775 | 17 | 0.3568 | 0.4426 | 0.3533 | 0.3623 | 0.1583 | 0.2805 | 0.0054 | 0.1768 | 0.0932 | 0.3287 | 0.3777 | **0.4847** |
| hcl_adrenalgland2 | 8,114 | 13 | 0.4663 | –<sup>g</sup> | 0.3921 | 0.4043 | 0.2068 | 0.2900 | –<sup>h</sup> | 0.2998 | 0.1631 | 0.5036 | 0.4238 | **0.5241** |
| hcl_kidney2 | 9,153 | 18 | 0.5051 | 0.2100 | 0.3179 | 0.3124 | 0.1233 | 0.1297 | 0.0438 | 0.3137 | 0.0803 | 0.4551 | 0.4585 | **0.5686** |
| hcl_trachea2 | 9,949 | 14 | **0.6631** | 0.0718 | 0.4960 | 0.5031 | 0.1699 | 0.3733 | 0.0711 | 0.3886 | 0.2123 | 0.5504 | 0.4884 | 0.5840 |
| Lung | 10,360 | 13 | 0.4734 | 0.1844 | 0.4133 | 0.3840 | 0.1447 | 0.1989 | 0.0628 | 0.1645 | –<sup>c</sup> | 0.4962 | 0.4729 | **0.6347** |
| ZhengMix | 11,749 | 8 | **0.7568** | 0.7452 | 0.6583 | 0.6668 | 0.2946 | 0.5583 | 0.1336 | 0.2448 | –<sup>c</sup> | 0.6955 | 0.7110 | 0.7504 |
| Cortex_P | 46,346 | 14 | **0.7567** | 0.5423 | 0.4269 | 0.4580 | 0.2623 | 0.4343 | 0.2476 | 0.3802 | –<sup>c</sup> | –<sup>f</sup> | 0.5149 | 0.5773 |
| Cortex_F | 71,639 | 14 | **0.8271** | 0.6150 | 0.4976 | 0.5485 | 0.3058 | 0.5343 | 0.2381 | 0.3140 | –<sup>c</sup> | –<sup>f</sup> | 0.5761 | 0.6284 |
| Cortex_P (K=93) | 46,346 | 93 | **0.5661** | 0.3074 | 0.3763 | 0.3223 | 0.1849 | 0.2886 | 0.1425 | 0.4057 | –<sup>c</sup> | –<sup>f</sup> | 0.3945 | 0.4118 |
| Cortex_F (K=81) | 71,639 | 81 | **0.5782** | 0.4178 | 0.4273 | 0.3919 | 0.2315 | 0.3916 | 0.1946 | 0.5013 | –<sup>c</sup> | –<sup>f</sup> | 0.4409 | 0.4751 |
| TS50W_raw | 508,731 | 42 | 0.6224 | –<sup>b</sup> | –<sup>e</sup> | –<sup>e</sup> | –<sup>c</sup> | –<sup>e</sup> | –<sup>c</sup> | –<sup>d</sup> | –<sup>c</sup> | –<sup>c</sup> | 0.5889 | **0.6237** |
| TS100W_raw | 1,136,218 | 180 | 0.6843 | –<sup>b</sup> | –<sup>e</sup> | –<sup>e</sup> | –<sup>c</sup> | –<sup>e</sup> | –<sup>c</sup> | –<sup>d</sup> | –<sup>c</sup> | –<sup>c</sup> | 0.6381 | **0.7503** |
| Mean |  |  | 0.6566 | 0.5731 | 0.5022 | 0.5094 | 0.1684 | 0.4302 | 0.3778 | 0.3808 | 0.4126 | 0.6182 | 0.4889 | 0.5429 |
| CUMAP wins |  |  | – | 39/57 | 50/58 | 48/58 | 55/58 | 47/58 | 53/55 | 53/58 | 40/46 | 47/54 | 52/59 | 46/59 |
| Wilcoxon p |  |  | – | 9.5e-4 | 2.0e-9 | 3.1e-8 | 6.4e-11 | 2.6e-8 | 5.6e-10 | 1.8e-9 | 4.7e-8 | 2.0e-7 | 1.7e-9 | 1.6e-6 |

### Appendix C.5. Per-dataset AMI, PP protocol

**Table C.24:** AMI on the PP condition (standard preprocessing), fixed weights [0.8, 0.1, 0.1] (TS full atlas: [0.00, 0.20, 0.80], see Section 4.9). Best per row in bold; a dash (–) marks an entry the method could not produce, with a superscript giving the reason (legend in the text). *CUMAP wins* and *Wilcoxon p* are computed on RAW and PP combined, so these two rows are identical in the RAW and PP tables of the same metric; the denominator is the number of paired entries the baseline actually produced.

| Dataset | Cells | K | CUMAP | scICE | tSNE | UMAP | KPCA | PHATE | SIMLR | scDCCA | scGAC | scAGCL | sGPT<sub>K</sub> | sGPT<sub>L</sub> |
| --- | --- | --- | --- | --- | --- | --- | --- | --- | --- | --- | --- | --- | --- | --- |
| Tian_sc_dropseq | 222 | 3 | 0.8597 | 0.8484 | 0.8908 | 0.8908 | **0.9083** | 0.8908 | 0.8030 | 0.8361 | 0.8453 | 0.8527 | 0.4456 | 0.5222 |
| kumar | 244 | 3 | **1.0000** | 0.9490 | 0.9282 | 0.9147 | 0.9391 | 0.8065 | 0.9432 | 0.8324 | 0.9793 | 0.8013 | 0.6066 | 0.5295 |
| koh | 529 | 9 | **0.9317** | 0.8586 | 0.8643 | 0.8574 | 0.8068 | 0.8841 | 0.7063 | 0.4547 | 0.6144 | 0.8304 | 0.4829 | 0.5841 |
| M1 | 820 | 13 | 0.7029 | **0.8724** | 0.6553 | 0.7015 | 0.5822 | 0.7017 | 0.6788 | 0.6018 | 0.7948 | 0.7173 | 0.5526 | 0.6412 |
| Tian_sc_10x | 893 | 3 | **1.0000** | **1.0000** | 0.7549 | **1.0000** | **1.0000** | **1.0000** | 0.9931 | **1.0000** | **1.0000** | 0.9931 | 0.9673 | 0.6419 |
| Chu | 1,017 | 7 | 0.8981 | 0.9252 | 0.9233 | 0.9244 | 0.6942 | 0.8989 | 0.8487 | **0.9468** | 0.8950 | 0.8786 | 0.8319 | 0.8136 |
| M2 | 1,054 | 13 | **0.8127** | 0.7777 | 0.6210 | 0.7035 | 0.5367 | 0.7232 | 0.5708 | 0.6174 | 0.5593 | 0.7534 | 0.4007 | 0.4450 |
| H4 | 1,293 | 14 | 0.8519 | **0.9136** | 0.6776 | 0.7273 | 0.6467 | 0.7509 | 0.6371 | 0.7129 | 0.6793 | 0.7574 | 0.5598 | 0.6285 |
| H2 | 1,720 | 14 | 0.8316 | **0.8767** | 0.6720 | 0.7947 | 0.5934 | 0.8724 | 0.6622 | 0.7718 | 0.5827 | 0.7758 | 0.5942 | 0.6681 |
| H1 | 1,925 | 14 | 0.8343 | **0.9023** | 0.7156 | 0.8089 | 0.5946 | 0.7654 | 0.6677 | 0.7539 | 0.6656 | 0.7582 | 0.5991 | 0.6651 |
| Enge | 2,279 | 6 | **0.8542** | 0.8126 | 0.6906 | 0.7828 | 0.5441 | 0.6971 | 0.5597 | 0.6725 | 0.7438 | 0.7484 | 0.5845 | 0.5890 |
| wu_kidney_droncseq | 2,769 | 13 | **0.5617** | 0.4801 | 0.4806 | 0.4876 | 0.3618 | 0.5356 | 0.3745 | 0.4358 | 0.2569 | 0.3312 | 0.3956 | 0.4169 |
| wu_kidney_snucdropseq | 3,011 | 13 | **0.4839** | 0.4111 | 0.4006 | 0.3818 | 0.3766 | 0.4772 | 0.2210 | 0.4047 | 0.2326 | 0.3162 | 0.3369 | 0.3775 |
| H3 | 3,600 | 14 | **0.8392** | 0.7816 | 0.6683 | 0.7791 | 0.6142 | 0.7843 | 0.6884 | 0.7139 | 0.6376 | 0.7370 | 0.6431 | 0.7135 |
| Tian_sc_10x_5cl | 3,863 | 5 | 0.9751 | **0.9760** | **0.9760** | 0.9154 | 0.9031 | 0.9757 | 0.9736 | 0.7531 | **0.9760** | 0.9656 | 0.6682 | 0.7639 |
| duodenum | 5,189 | 10 | **0.7811** | 0.7025 | 0.6770 | 0.7347 | 0.6566 | 0.7496 | 0.6874 | 0.6523 | 0.7466 | 0.6802 | 0.6429 | 0.6560 |
| mca_fetalintestine | 6,076 | 13 | 0.5909 | 0.5461 | 0.5101 | 0.5218 | 0.2950 | **0.5985** | 0.2903 | 0.5355 | 0.3148 | 0.2793 | 0.3302 | 0.4480 |
| mca_fetalstomach | 6,191 | 16 | 0.4295 | 0.2544 | 0.4626 | 0.4753 | 0.2497 | **0.4906** | 0.3386 | 0.3972 | 0.1931 | 0.2846 | 0.3158 | 0.3863 |
| hcl_peripheralblood4 | 7,159 | 13 | 0.5852 | 0.6328 | 0.5980 | 0.6110 | 0.4666 | **0.6381** | 0.2792 | 0.5947 | 0.1695 | 0.5392 | 0.4047 | 0.4767 |
| hcl_muscle1 | 7,775 | 17 | **0.5432** | 0.5364 | 0.4983 | 0.4920 | 0.3908 | 0.4995 | 0.3116 | 0.5188 | 0.1456 | 0.3718 | 0.4375 | 0.5014 |
| hcl_adrenalgland2 | 8,114 | 13 | 0.4270 | 0.4473 | 0.5305 | 0.5394 | 0.4294 | 0.5817 | <sup>h</sup> | **0.6537** | 0.2028 | 0.3518 | 0.4540 | 0.5350 |
| hcl_kidney2 | 9,153 | 18 | **0.6395** | 0.4525 | 0.6061 | 0.6196 | 0.3887 | 0.5796 | 0.4478 | 0.6307 | 0.2051 | 0.3703 | 0.4853 | 0.5724 |
| hcl_trachea2 | 9,949 | 14 | **0.7306** | 0.5235 | 0.6542 | 0.6636 | 0.4908 | 0.6843 | 0.5891 | 0.6858 | 0.3739 | 0.6010 | 0.5736 | 0.6892 |
| Lung | 10,358 | 13 | **0.5986** | 0.3336 | 0.5691 | 0.5582 | 0.3539 | 0.5977 | <sup>h</sup> | 0.4426 | <sup>c</sup> | 0.5118 | 0.4351 | 0.5711 |
| ZhengMix | 11,749 | 8 | **0.7457** | 0.6851 | 0.5939 | 0.6633 | 0.5846 | 0.6293 | 0.5462 | 0.4482 | <sup>c</sup> | 0.6256 | 0.5375 | 0.5911 |
| Cortex_P | 46,341 | 14 | 0.8295 | **0.8518** | 0.4080 | 0.6922 | 0.5765 | 0.6918 | 0.6904 | 0.5955 | <sup>c</sup> | 0.6177 | 0.3499 | 0.4685 |
| Cortex_F | 71,594 | 14 | 0.8729 | **0.9076** | 0.0845 | 0.7627 | 0.5064 | 0.6788 | 0.6946 | 0.6418 | <sup>c</sup> | 0.6898 | 0.4421 | 0.5432 |
| Cortex_P (K=93) | 46,341 | 93 | 0.5118 | **0.5998** | 0.4309 | 0.5687 | 0.3812 | 0.5230 | 0.3975 | 0.5004 | <sup>c</sup> | 0.4944 | <sup>a</sup> | <sup>a</sup> |
| Cortex_F (K=81) | 71,594 | 81 | 0.5803 | **0.6566** | 0.0902 | 0.5774 | 0.3841 | 0.5132 | 0.5145 | 0.5510 | <sup>c</sup> | 0.5225 | <sup>a</sup> | <sup>a</sup> |
| TS50W_pp100 | 506,767 | 42 | **0.6401** | <sup>b</sup> | <sup>e</sup> | <sup>e</sup> | <sup>c</sup> | <sup>e</sup> | <sup>c</sup> | <sup>d</sup> | <sup>c</sup> | <sup>c</sup> | 0.5047 | 0.6003 |
| TS100W_pp | 1,129,368 | 180 | **0.6261** | <sup>b</sup> | <sup>e</sup> | <sup>e</sup> | <sup>c</sup> | <sup>e</sup> | <sup>c</sup> | <sup>d</sup> | <sup>c</sup> | <sup>c</sup> | <sup>a</sup> | <sup>a</sup> |
| Mean |  |  | 0.7280 | 0.7074 | 0.6080 | 0.6948 | 0.5606 | 0.6972 | 0.5969 | 0.6330 | 0.5571 | 0.6261 | 0.5208 | 0.5728 |
| CUMAP wins |  |  | – | 39/57 | 50/58 | 48/58 | 55/58 | 47/58 | 53/55 | 53/58 | 40/46 | 47/54 | 52/59 | 46/59 |
| Wilcoxon p |  |  | – | 9.5e-4 | 2.0e-9 | 3.1e-8 | 6.4e-11 | 2.6e-8 | 5.6e-10 | 1.8e-9 | 4.7e-8 | 2.0e-7 | 1.7e-9 | 1.6e-6 |

### Appendix C.6. Implementations and the scale at which each is required

**Table C.25:** The three implementations and the scale at which each is required.

| Scale | Implementation | Reason |
| --- | --- | --- |
| $\leq 10^4$ cells | Dense distances → distance-level fusion | Exact distances are affordable |
| $10^4$–$10^5$ | Cached dense / PCA-100 graph-level fusion | Dense $N \times N$ no longer fits |
| $> 10^5$ | Approximate $k$NN (lean) on GPU | 500K cells → 1 TB; 1.1M → 5 TB |

### Appendix C.7. Label-informative genes discarded by feature selection

**Table C.26:** Label-informative genes discarded by feature selection.

| Dataset | A: DE lost | B: marker lost | C: intersection lost |
| --- | --- | --- | --- |
| TS full atlas | 91.2% | 88.7% | 83.3% |
| TS immune | 80.2% | 77.5% | 67.3% |
| Cortex frontal | 50.1% | 71.5% | 40.9% |
| Cortex posterior | 49.7% | 70.4% | 37.5% |
| ZhengMix | 64.0% | 74.7% | 46.3% |
| Lung atlas | 50.3% | 65.7% | 29.9% |
| HCL trachea | 44.1% | 62.9% | 23.9% |
| HCL kidney | 51.8% | 65.8% | 12.8% |
| HCL adrenal | 50.6% | 58.8% | 28.8% |
| HCL muscle | 39.8% | 61.8% | 21.1% |
| HCL blood | 56.5% | 66.7% | 24.6% |
| HCL duodenum | 48.2% | 63.4% | 24.9% |
| MCA fetal stomach | 45.8% | 66.2% | 11.0% |
| MCA fetal intestine | 38.3% | 74.3% | 19.4% |
| Tian sc 10x 5-line | 27.6% | N/A | N/A |
| Tian sc 10x | 20.7% | N/A | N/A |
| Tian sc Drop-seq | 38.0% | N/A | N/A |
| Pancreas H1 | 37.0% | 54.5% | 16.6% |
| Pancreas H2 | 38.6% | 50.8% | 13.2% |
| Pancreas H3 | 38.6% | 49.6% | 10.1% |
| Pancreas H4 | 36.4% | 44.7% | 16.1% |
| Pancreas M1 | 48.8% | 64.9% | 24.1% |
| Pancreas M2 | 46.9% | 58.9% | 18.9% |
| Wu sNuc-DropSeq | 39.7% | N/A | N/A |
| Wu DroNc-seq | 34.6% | N/A | N/A |
| Enge | 51.3% | 63.6% | 39.6% |
| Chu | 37.7% | N/A | N/A |
| Koh | 80.0% | N/A | N/A |
| Kumar | 94.7% | N/A | N/A |

### Appendix C.8. Discarded genes that are both top-ranked DE genes and curated markers

*Failure-case legend.*. The superscripts in Tables C.21–C.24 are:

a. **scGPT: not run**. scGPT was given the RAW count matrix for every dataset and the PP matrix for 28 of the 29; the full atlas is the one dataset the PP pass does not cover. The two fine-grained cortex rows would need a *K* = 81 and a *K* = 93 clustering of the PP embedding, which we did not run; their RAW counterparts are reported.
b. **scICE, Tabula Sapiens: no output**. Its graph construction is single-core; the 508k-cell run was still running after 7 days and produced nothing.
c. **Out of memory (***n*^2^ **or** *N × N* ). scGAC builds an *N × N* attention matrix; SIMLR builds a dense multiple-kernel similarity matrix (1,913 GB of *n*^2^ doubles at 508k cells against 502 GB available); exact KernelPCA needs the 508,731^2^ RBF kernel at ≈2 TB; scAGCL runs out of memory at the million-cell scale.
d. **scDCCA, Tabula Sapiens: diverged**. The zero-inflated negative-binomial loss went numerically undefined within the first epoch and stayed so for all 100 epochs, collapsing every cell into one cluster. We report the failure rather than the degenerate score.
e. **Exact form infeasible at** ≥ 5 *×* 10^5^ **cells**. t-SNE is *O*(*n*^2^); UMAP and PHATE only reach this scale through GPU or landmark approximations, which are no longer the textbook method being compared.
f. **scAGCL, cortex RAW: wall-clock cap**. These two cells hit our 2-hour limit (7,200 s) without finishing. This is a timeout, *not* an out-of-memory failure, and we label it separately.
g. **scICE, adrenal gland RAW: no stable clustering**. The run finished, but its stability filter kept none of the candidate cluster numbers, so the output holds the cell identifiers and no labels at all. On the same dataset after preprocessing the filter keeps four candidates.
h. **SIMLR: crash in its final t-SNE step**. The method itself completes: the PCA, the *k*-nearest-neighbour search, the multiple kernels, the five iterations and the *K*-means all finish. The process then aborts inside the t-SNE call that produces the visualisation, with heap corruption reported by the allocator. We rediagnosed all three cells by rerunning them and each failed at the same point.

**Table C.27:**
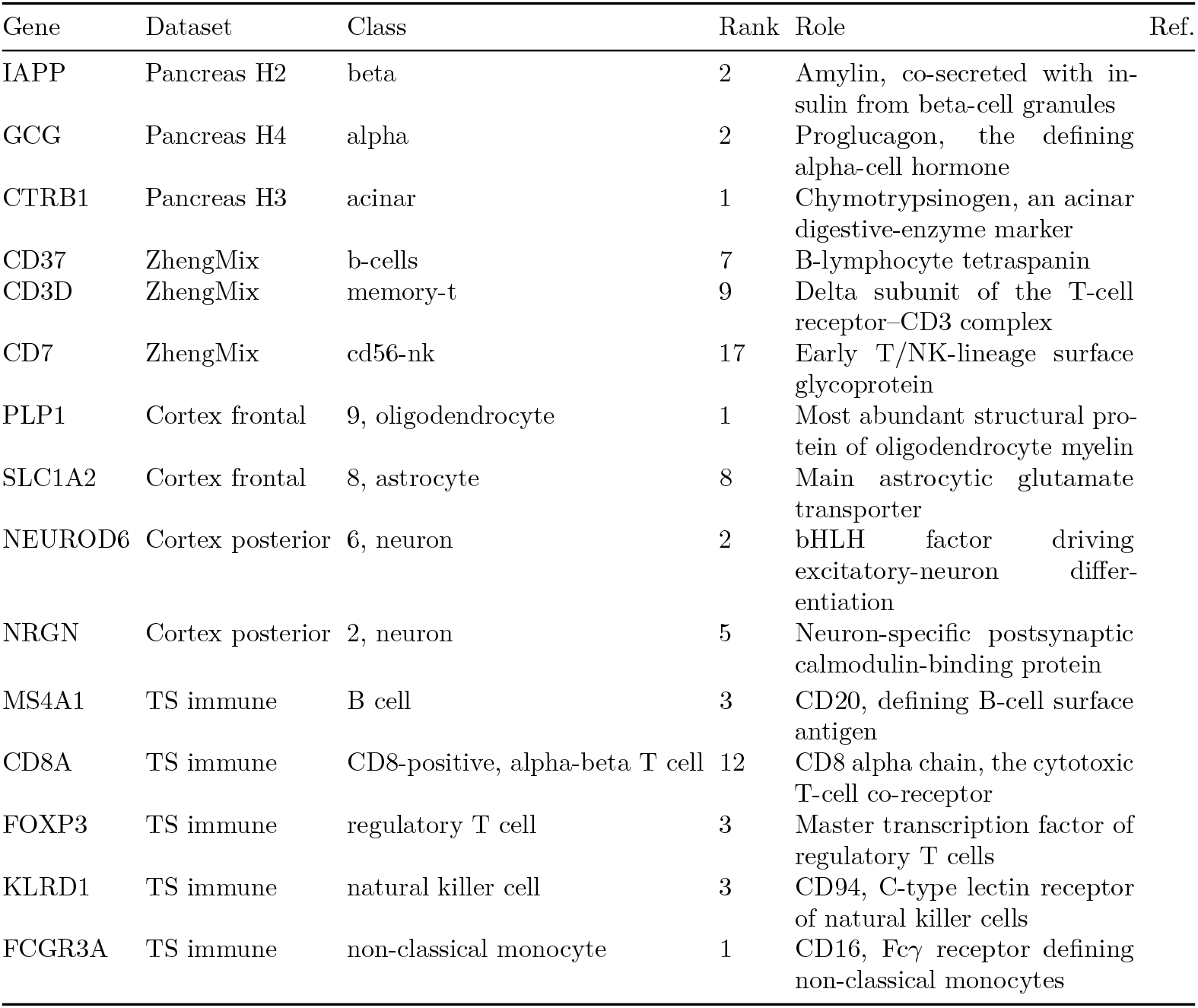
Discarded genes that are both top-ranked DE genes and curated markers.

| Gene | Dataset | Class | Rank | Role | Ref. |
| --- | --- | --- | --- | --- | --- |
| IAPP | Pancreas H2 | beta | 2 | Amylin, co-secreted with insulin from beta-cell granules |  |
| GCG | Pancreas H4 | alpha | 2 | Proglucagon, the defining alpha-cell hormone |  |
| CTRB1 | Pancreas H3 | acinar | 1 | Chymotrypsinogen, an acinar digestive-enzyme marker |  |
| CD37 | ZhengMix | b-cells | 7 | B-lymphocyte tetraspanin |  |
| CD3D | ZhengMix | memory-t | 9 | Delta subunit of the T-cell receptor-CD3 complex |  |
| CD7 | ZhengMix | cd56-nk | 17 | Early T/NK-lineage surface glycoprotein |  |
| PLP1 | Cortex frontal | 9, oligodendrocyte | 1 | Most abundant structural protein of oligodendrocyte myelin |  |
| SLC1A2 | Cortex frontal | 8, astrocyte | 8 | Main astrocytic glutamate transporter |  |
| NEUROD6 | Cortex posterior | 6, neuron | 2 | bHLH factor driving excitatory-neuron differentiation |  |
| NRGN | Cortex posterior | 2, neuron | 5 | Neuron-specific postsynaptic calmodulin-binding protein |  |
| MS4A1 | TS immune | B cell | 3 | CD20, defining B-cell surface antigen |  |
| CD8A | TS immune | CD8-positive, alpha-beta T cell | 12 | CD8 alpha chain, the cytotoxic T-cell co-receptor |  |
| FOXP3 | TS immune | regulatory T cell | 3 | Master transcription factor of regulatory T cells |  |
| KLRD1 | TS immune | natural killer cell | 3 | CD94, C-type lectin receptor of natural killer cells |  |
| FCGR3A | TS immune | non-classical monocyte | 1 | CD16, Fcγ receptor defining non-classical monocytes | |

<sup>c</sup> **Out of memory ( $n^2$ or $N \times N$ ).** scGAC builds an $N \times N$ attention matrix; SIMLR builds a dense multiple-kernel similarity matrix (1,913 GB of $n^2$ doubles at 508k cells against 502 GB available); exact KernelPCA needs the $508,731^2$ RBF kernel at $\approx 2$ TB; scAGCL runs out of memory at the million-cell scale.
<sup>d</sup> **scDCCA, Tabula Sapiens: diverged.** The zero-inflated negative-binomial loss went numerically undefined within the first epoch and stayed so for all 100 epochs, collapsing every cell into one cluster. We report the failure rather than the degenerate score.
<sup>e</sup> **Exact form infeasible at $\geq 5 \times 10^5$ cells.** t-SNE is $O(n^2)$ ; UMAP and PHATE only reach this scale through GPU or landmark approximations, which are no longer the textbook method being compared.
<sup>f</sup> **scAGCL, cortex RAW: wall-clock cap.** These two cells hit our 2-hour limit (7,200 s) without finishing. This is a timeout, *not* an out-of-memory failure, and we label it separately.
<sup>g</sup> **scICE, adrenal gland RAW: no stable clustering.** The run finished, but its stability filter kept none of the candidate cluster numbers, so the output holds the cell identifiers and no labels at all. On the same dataset after preprocessing the filter keeps four candidates.

## Appendix D. Additional embedding comparisons

### Appendix D.1. Each dataset under both protocols

Each dataset below is shown under both the **Raw** protocol (all methods receive the same unprocessed count matrix) and the **PP** protocol (each baseline uses its own recommended pipeline; c-UMAP continues to read raw counts). Point color is the ground-truth cell type and is identical across panels within a figure. These complement the main-text Cortex Frontal (Figs. 8–9) and ZhengMix (Figs. 10–11) comparisons; the complete set of 56 figures covering all 29 datasets under both protocols is released with the appendix package (Section 4.8). The figures below give the pancreas donors H2 to H4, the mouse donors M1 and M2, and the posterior cortex atlas; H1 and the frontal cortex atlas are in the manuscript.

**Figure D.16:**
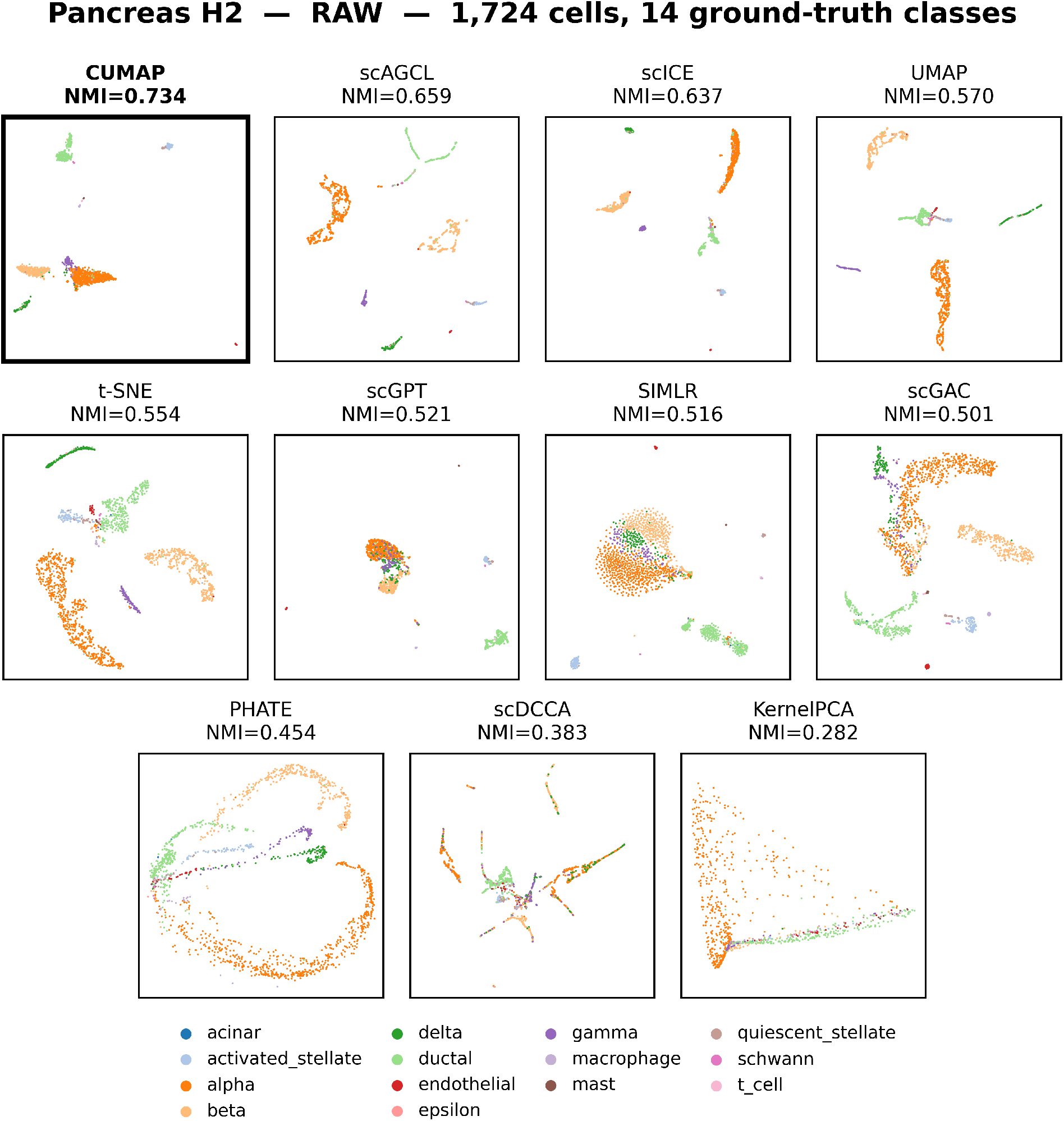
GSE84133 H2 (human pancreas), Raw protocol. (1,724 cells, 14 ground-truth classes). Each panel is one method in its own embedding, colored by ground-truth cell type.

**Figure D.17:**
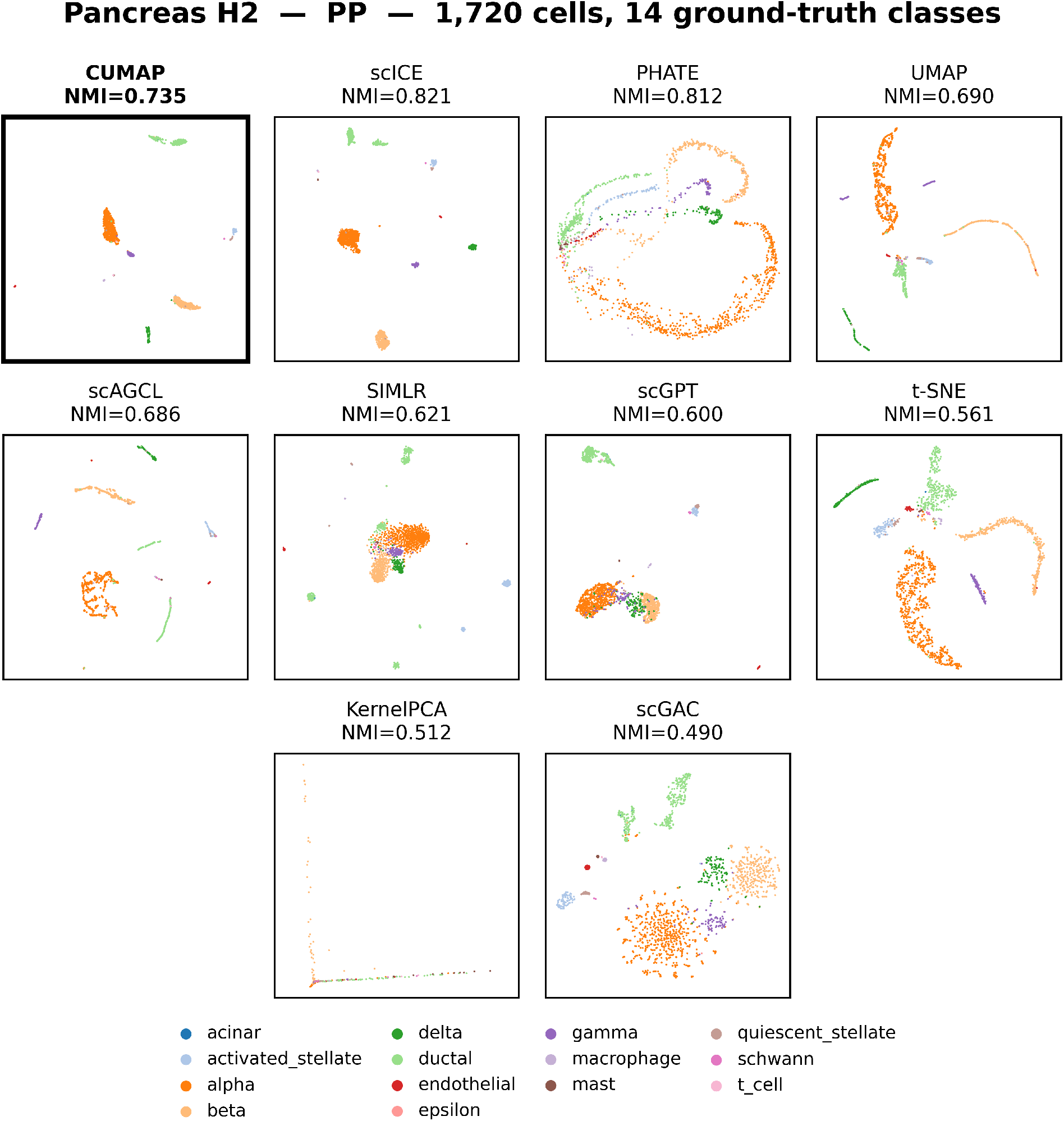
GSE84133 H2 (human pancreas), PP protocol. (1,724 cells, 14 ground-truth classes). Each panel is one method in its own embedding, colored by ground-truth cell type.

**Figure D.18:**
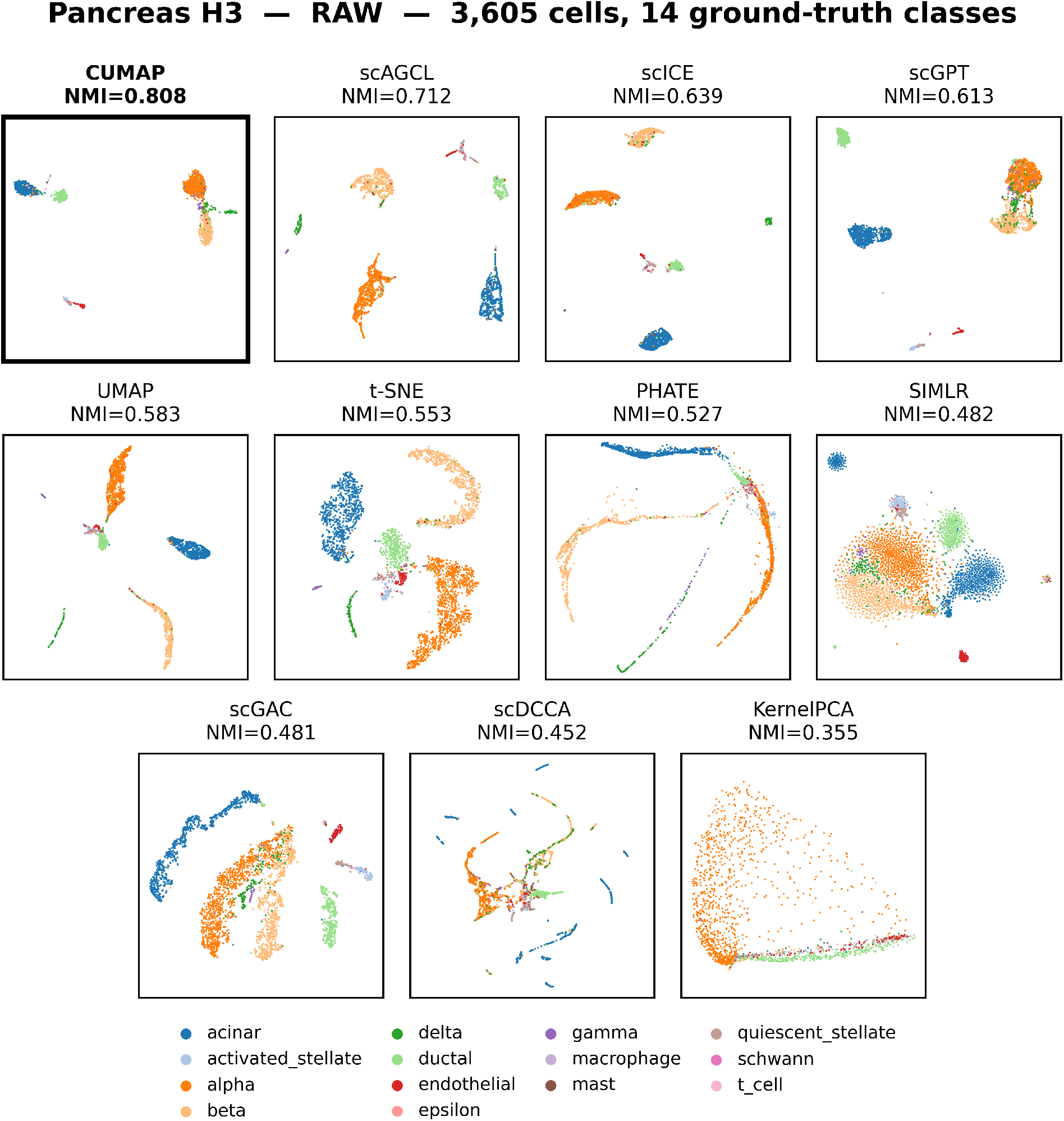
GSE84133 H3 (human pancreas), Raw protocol. (3,605 cells, 14 ground-truth classes). Each panel is one method in its own embedding, colored by ground-truth cell type.

**Figure D.19:**
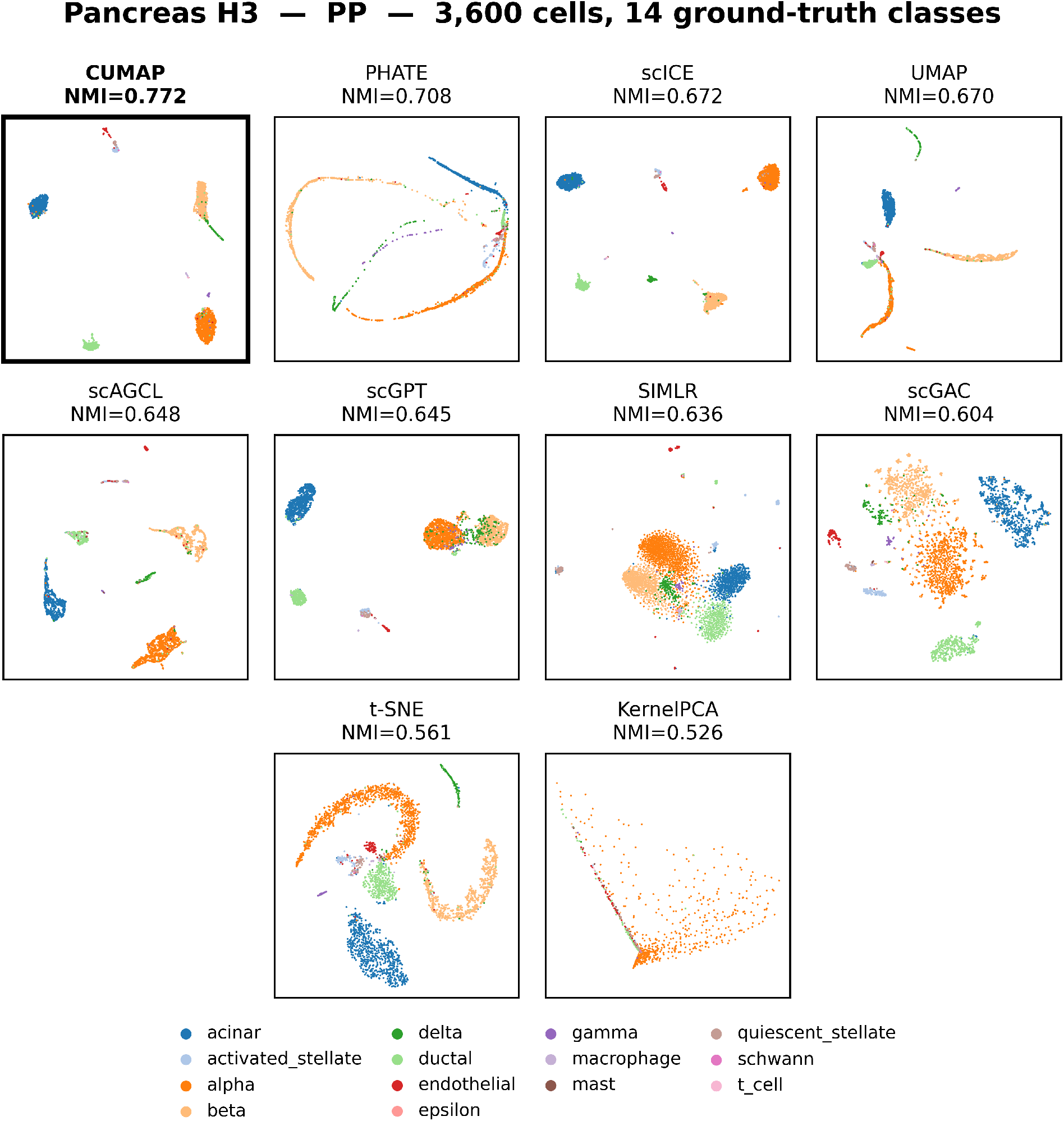
GSE84133 H3 (human pancreas), PP protocol. (3,600 cells, 14 ground-truth classes). Each panel is one method in its own embedding, colored by ground-truth cell type.

**Figure D.20:**
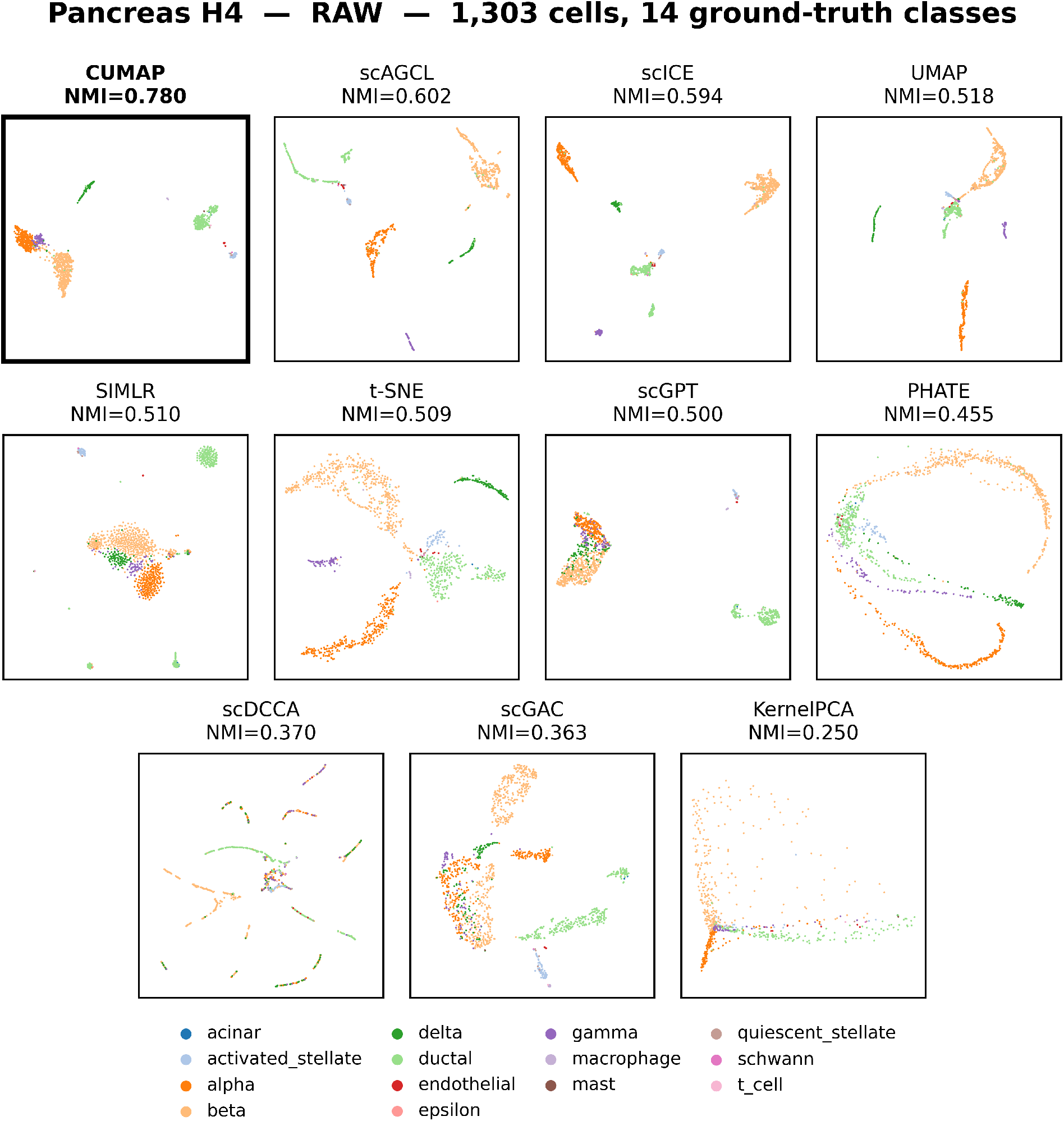
GSE84133 H4 (human pancreas, T2D donor), Raw protocol. (1,303 cells, 14 ground-truth classes). Each panel is one method in its own embedding, colored by ground-truth cell type.

**Figure D.21:**
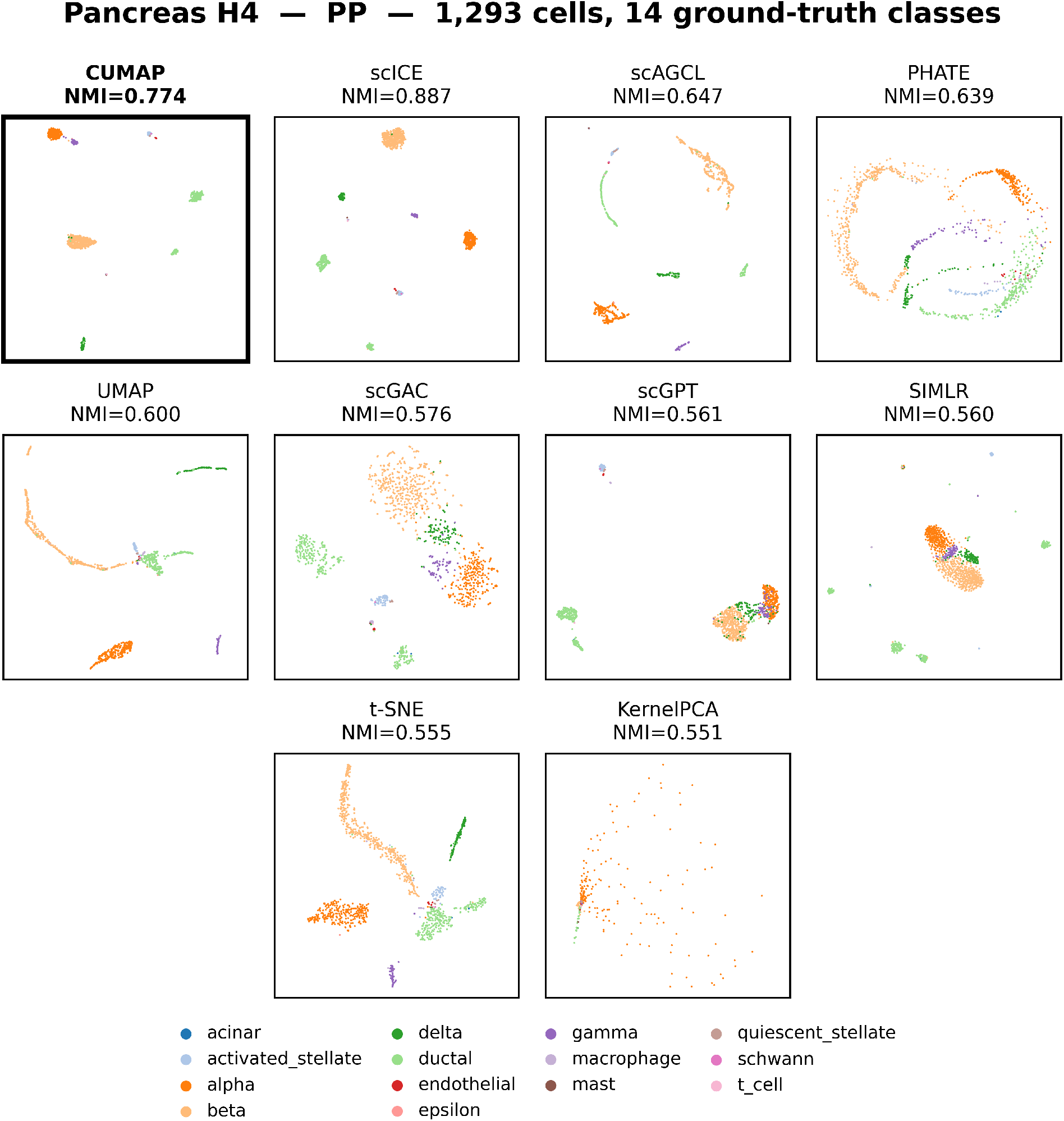
GSE84133 H4 (human pancreas, T2D donor), PP protocol. (1,303 cells, 14 ground-truth classes). Each panel is one method in its own embedding, colored by ground-truth cell type.

**Figure D.22:**
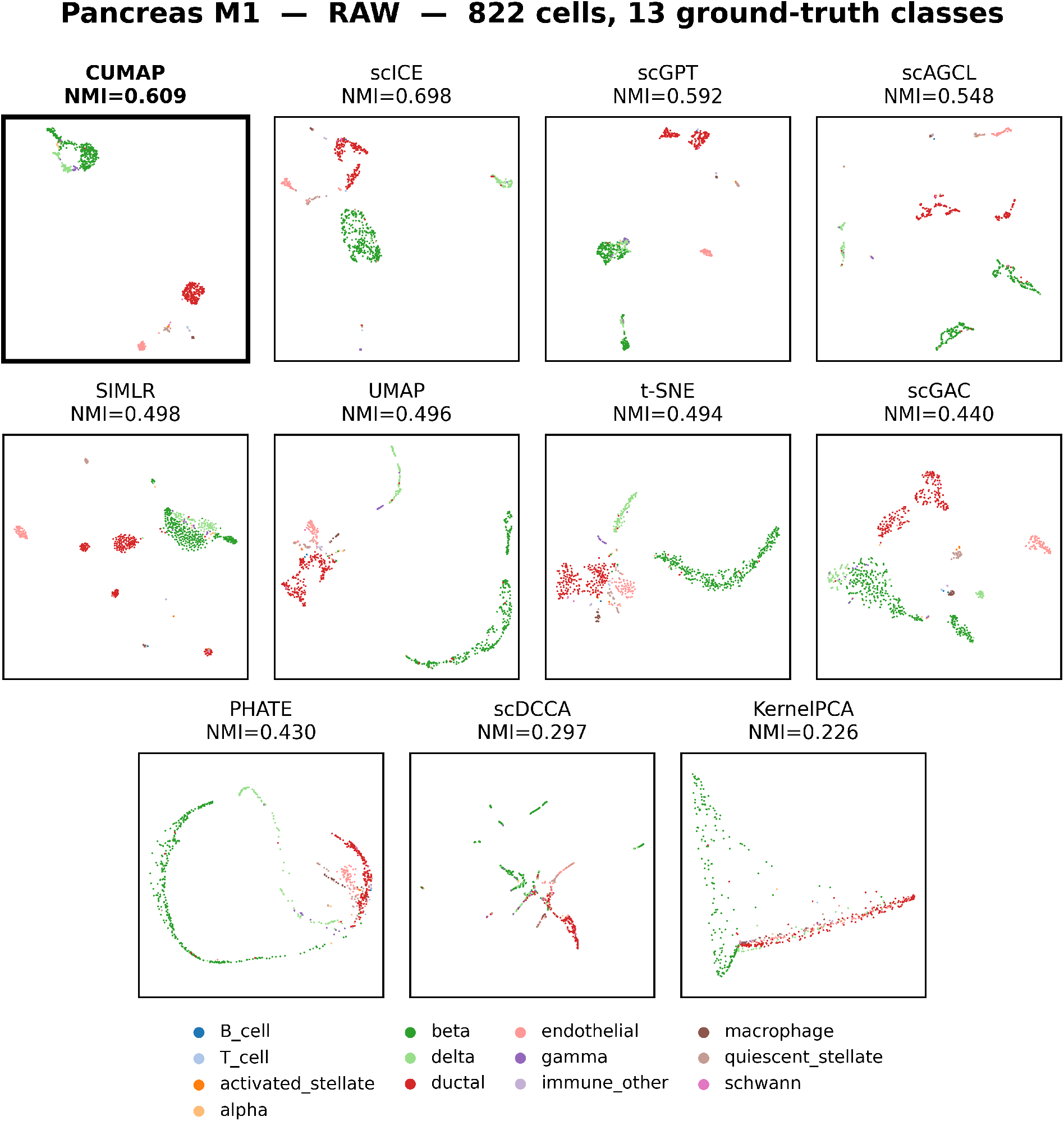
GSE84133 M1 (mouse pancreas), Raw protocol. (822 cells, 13 ground-truth classes). Each panel is one method in its own embedding, colored by ground-truth cell type.

**Figure D.23:**
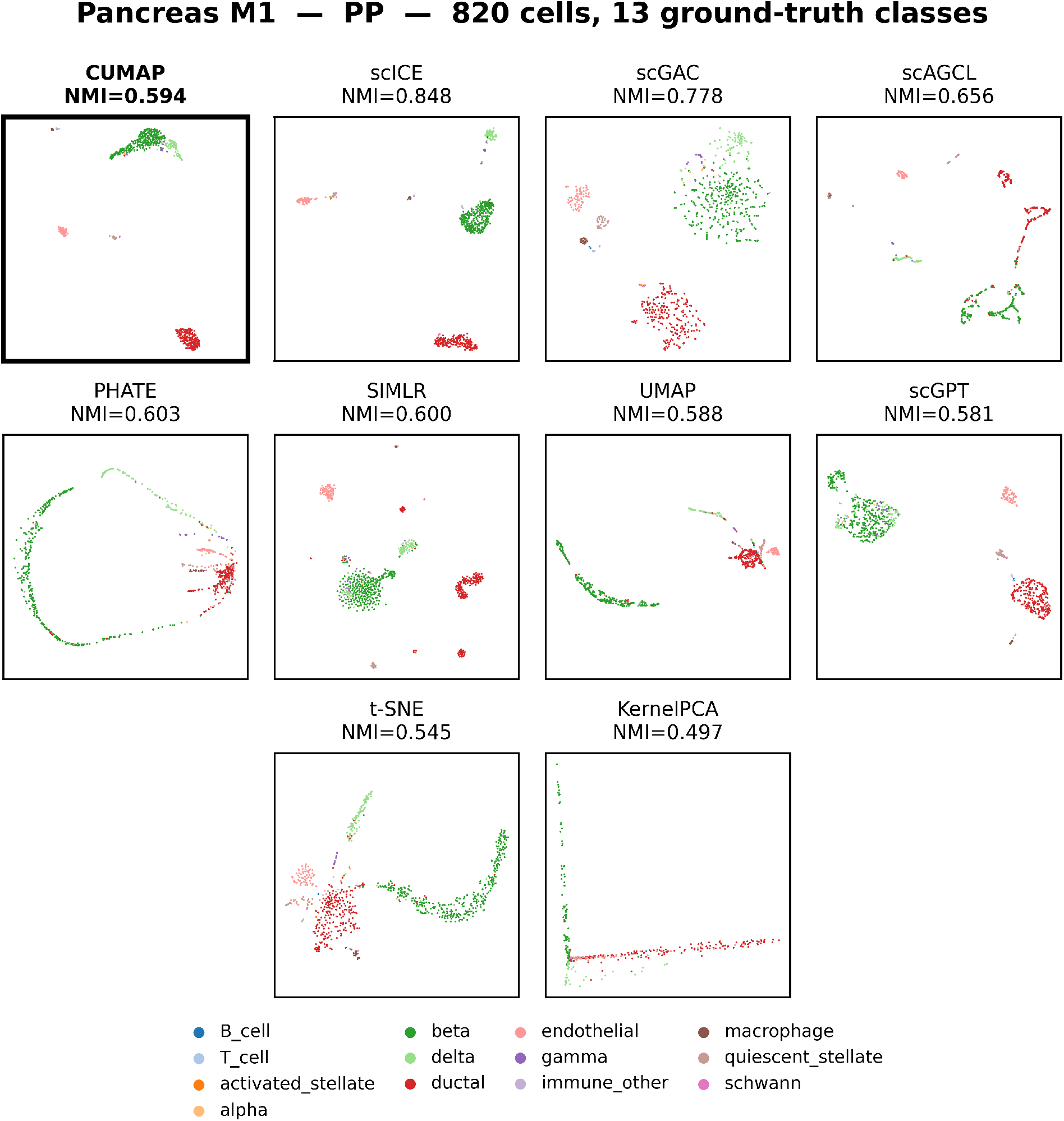
GSE84133 M1 (mouse pancreas), PP protocol. (822 cells, 13 ground-truth classes). Each panel is one method in its own embedding, colored by ground-truth cell type.

**Figure D.24:**
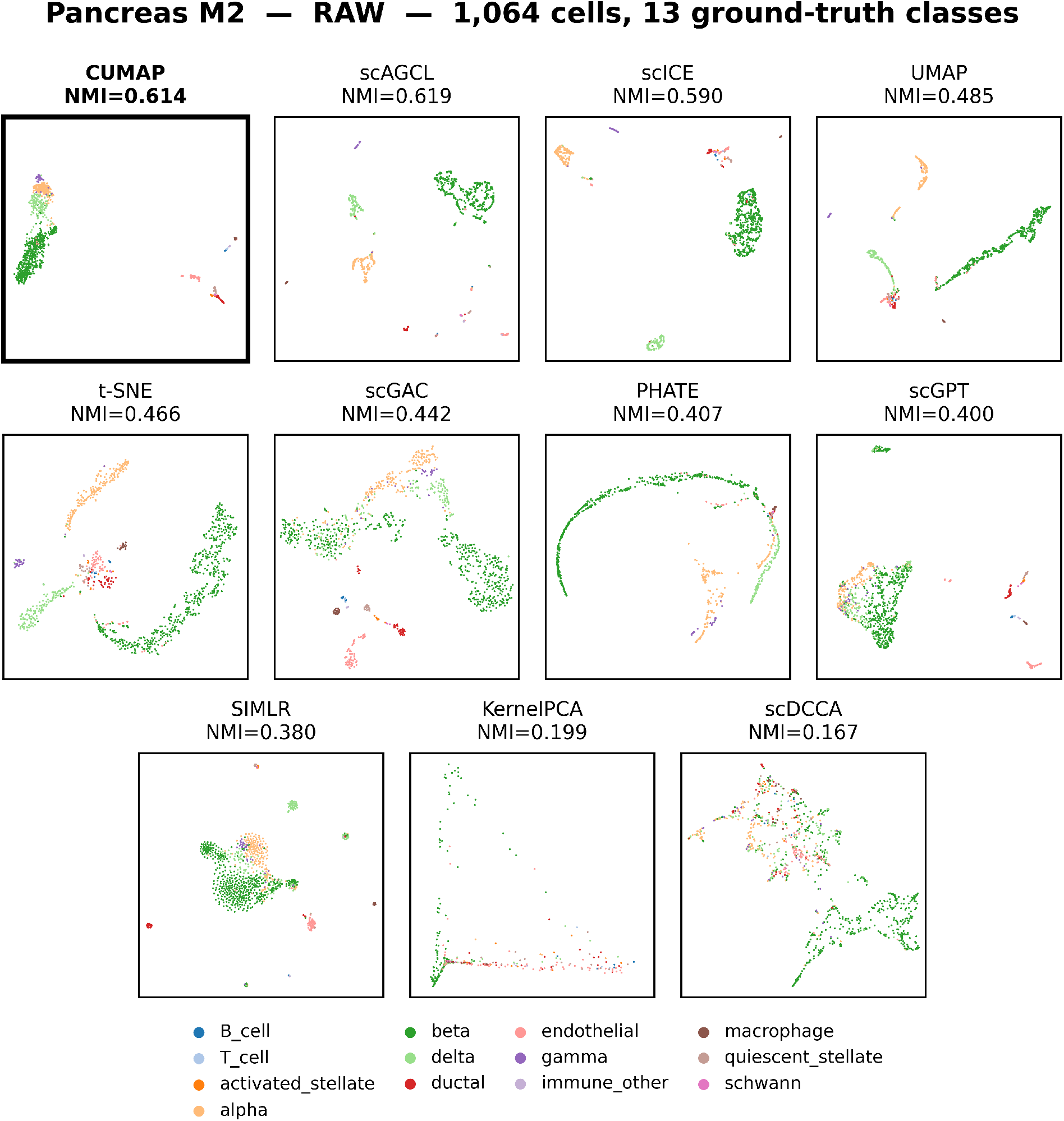
GSE84133 M2 (mouse pancreas), Raw protocol. (1,064 cells, 13 ground-truth classes). Each panel is one method in its own embedding, colored by ground-truth cell type.

**Figure D.25:**
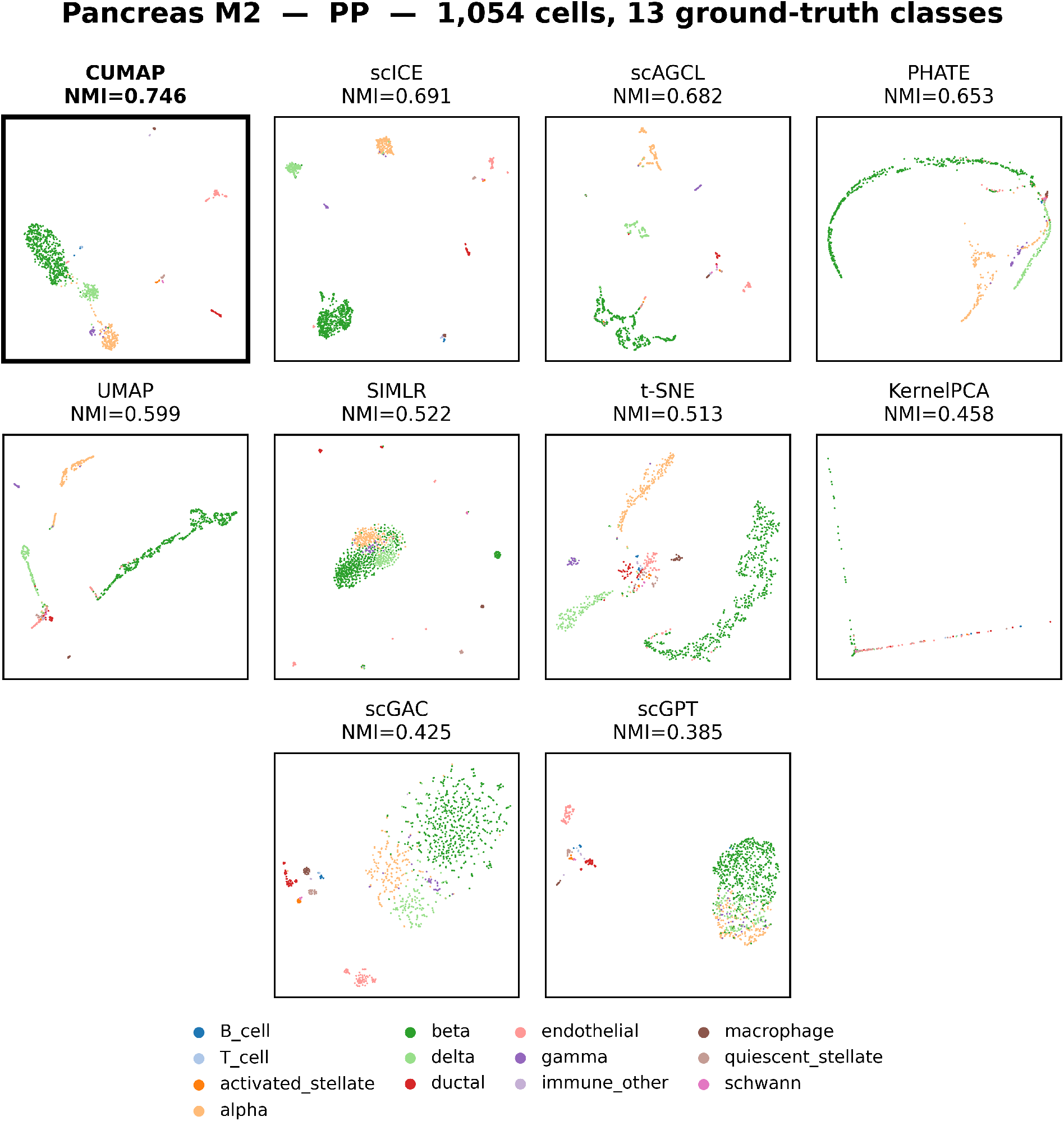
GSE84133 M2 (mouse pancreas), PP protocol. (1,064 cells, 13 ground-truth classes). Each panel is one method in its own embedding, colored by ground-truth cell type.

## Notes

https://tabula-sapiens.sf.czbiohub.org/

### Competing Interest Statement

The authors have declared no competing interest.

